# SEQUENCE SLIDER: integration of structural and genetic data to characterize isoforms from natural source

**DOI:** 10.1101/2021.09.16.460517

**Authors:** Rafael J. Borges, Guilherme H. M. Salvador, Daniel C. Pimenta, Lucilene D. dos Santos, Marcos R. M. Fontes, Isabel Usón

## Abstract

Proteins isolated from natural source can be composed of a mixture of isoforms with similar physicochemical properties that coexists in final steps of purification, toxins being prominent examples. Sequence composition is enforced throughout structural studies even when unsubstantiated. Herein, we propose a novel perspective to address the usually neglected heterogeneity of natural products by integrating biophysical, genetic and structural data in our program SEQUENCE SLIDER.

The aim is to assess the evidence supporting chemical composition in structure determination. Locally, we interrogate the experimental map to establish which side chains are supported by the structural data and the genetic information relating sequence conservation is integrated in this statistic. Hence, we build a constrained peptide database, containing most probable sequences to interpret mass spectrometry data (MS). In parallel, we perform MS *de novo* sequencing with genomic-based algorithms that foresee point mutations.

We calibrated SLIDER with *Gallus gallus* lysozyme, for which sequence is unequivocally established and numerous natural isoforms are reported. We used SLIDER to characterise a metalloproteinase and a phospholipase A2-like protein from the venom of *Bothrops moojeni* and a crotoxin from *Crotalus durissus collilineatus*. This integrated approach offers a more realistic structural descriptor to characterize macromolecules isolated from natural source.

**Key points:**

- The method SEQUENCE SLIDER integrates biophysical, genetic and structural data to assign sequence.
- It joins crystallography, mass spectrometry and phylogenetic data to characterize isoforms.
- Sequence heterogeneity of four proteins purified directly from snake venom was established.

## INTRODUCTION

Structural biology has been invaluable to unravel the molecular mechanisms of macromolecules and their complexes, as their structures convey function. The observation of different states and chemical reaction intermediates has provided the base for major breakthroughs in the last six decades with X-ray crystallography being the most used structural technique. A validated crystallographic model from high resolution data with global indicators conforming to the norm is usually unquestioned by the scientific community. It will be adopted to interpret biochemical data and to perform theoretical studies, such as homology modelling, molecular dynamics, docking and virtual screening, even if some local features included in the model may be undetermined (1). Since the resolution revolution, single particle cryoelectron microscopy (cryoEM) has reached atomic resolution with apoferritin (2, 3). Homogenous samples, favoured by high symmetry and low flexibility represent particularly favourable cases, whereas the typical resolution in a cryoEM reconstruction varies locally between 3-20 Å. In regions with resolution better than 3 Å, side chain modelling becomes possible, but it is subject to specific issues leading to side chains strongly affected by radiation damage or carrying net negative charges frequently becoming invisible in the electrostatic potential map (4, 5).

CryoEM obviates the need for crystallization, relieving the difficulty to study complex structures such as ribosomes, ionic channels and viral proteins. Various of these samples are obtained directly from the natural source, such as the trimeric spike protein isolated from the severe acute respiratory syndrome coronavirus 2 (6), ryanodine receptor isoform 1 purified from rabbit skeletal muscle (7–9) and its isoform 2 from dog heart ventricles (10), Inositol 1,4,5-trisphosphate receptor type 1 from rat cerebellum (EMD-6369) and Nav1.4-β1 Complex from electric eel (11).

Prior knowledge of the sample composition is essential for building the macromolecule in the typical scenario where structural data do not reach atomic resolution. For proteins produced recombinantly, sequence knowledge comes beforehand, whereas investigating structures obtained from natural sources, isoforms that share physicochemical properties may coexist in final steps of purification. Lysozyme, the most studied protein in crystallography for example (12), has been seen in at least four isoforms in bovine cartilage (13) and three in the medicinal leech (14). Most relevant is the case of natural products where concerted action of isoforms enhances their function, such as natural venoms. They are composed by a cocktail of toxins characterized by one of the most rapid evolutionary divergence and variability in any category of proteins (15). In fact, variability in venom composition within a single species may be related to ontogeny (16), diet (17), seasonality (18), geographical location (19) and gender (20, 21). For the snake venom phospholipases A2 (PLA_2_s), at least 16 isoforms of β-bungarotoxin (22) and 16 isoforms of crotoxin (23, 24) have been identified and characterized. The precise isoform may determine activity, as an example, two single point mutation Phe-Ile124 (phenylalanine to isoleucine in residue position 124) and Lys-Glu128 in the ammodytoxin are enough to change their toxicity, anticoagulant properties and toxin targets (25). Common to crystallography and cryoEM structures, researchers may assume their sample coincides with one entry on the sequence and structural repositories, therefore, bias propagation of established sequences is a concern in the field of natural products. Moreover, repositories are not error-free as they are curated at certain level mostly with global validating statistics and the information ultimately depends on depositors.

For non-recombinant samples, crystallographic and cryoEM determination has been effectively complemented by independent genetic information. A sophisticated analysis of electron density or potential volumes was implemented in the method *findMySequence,* which aims to identify the most plausible protein sequence in a database given a density map (45). It implements a sequence alignment based on a machine-learning residue-type classifier (45). Relevant genetic information can be derived from evolutionary conservation and amino acid frequencies from phylogenetic relations between homologues (26, 27), as residues related to function are conserved and structural stability restricts concerted pairwise changes (28–31). Such information has also proven useful for predicting protein structures (implemented in AlphaFold (32) and 3Dseq (33)) and using fragments from these for *ab initio* phasing of crystallographic structures using Molecular Replacement (implemented in Ample (34)), as they infer nearby residues analysing evolutionary covariance (32–34).

Mass spectrometry (MS), Edman degradation and cDNA sequencing are the complementary experimental techniques that allow determination of protein sequences directly purified from natural sources (35). MS does not require purity and has been a breakthrough characterizing complex samples in the advent of the omics, proteo, genome and transcriptomics (36, 37). Improvement was done to the particular application of omics fields to characterize venom, naming it as venomics, that may characterize the variability between different isoforms with amounts as small 0.05% (37, 38). Conventional identification approaches require concordance between the fragmented peptides from MS/MS with the *in-silico* digested sequences from a database, although unknown sequences or proteins with strong polymorphism are not revealed (reviews in (39–41)). Alternatives are *de novo* sequencing, that relies only on the experimental data, usually at the price of compromising coverage, and algorithms that tolerates mismatches, point mutations and post-translation modifications (PTM). Determining side chain composition in structures of samples containing multiple isoforms will not be trivial and will require the conjunction of different biophysical techniques, including MS.

The integrated use of MS and structural determination has proven invaluable in multiple ways comprising protein-construct design, purification, crystallization, phasing and model building (42). MS can define compact domains and mobile regions guiding truncation to improve sample preparation, assess the components present in a crystal, measure number of heavy atoms in a crystal and address issues in modelling, topology and side-chain proximity (42). Diemer *et al* characterized a Phosphate Binding Protein purified from the human plasma absent in eukaryotic genome databases by tandem use of crystallography and MS data combining multiple alternative digestions to obtain the correct sequence (43). Guo *et al.* (44) characterized the sequence haemoglobins from two endangered felines by use of crystallography, MS data from a single proteolysis and evaluation of homologue sequences (44). The side chain evaluator method called SEQUENCE SLIDER (SLIDER) provides a framework to model this complex task of integrating structural and genetic information. SLIDER (46) was first created in the *ab initio* phasing scope of ARCIMBOLDO method (47), in which small fragments are identified in the asymmetric unit using Molecular Replacement (48) and the rest of the structure is revealed through density modification and automatic map interpretation (49). SLIDER uses available sequence information, secondary structure prediction or alignment between remote homologues, to build most probable side-chain atoms into partial solution usually composed of polyalanine fragments, a discrimination in global statistics of the agreement between data and model may reveal true sequence hypothesis (46).

Herein, we describe SLIDER, implemented to integrate structure determination, mass spectrometry and genetic information, to address the heterogeneity of complex samples purified from natural sources building a probability of amino acids per residue basis. We calibrate SLIDER using a crystallographic dataset of the known hen egg-white lysozyme from *Gallus gallus* and we apply it to elucidate a metalloproteinase (BmooMP-I) (50) and a PLA_2_-like protein (MjTX-I) from *Bothrops moojeni* and a crotoxin from *Crotalus durissus collineatus* (CBCol), revealing sequences not yet seen in literature. We offer the structural community a methodology that consistently assigns amino acids integrating available information for macromolecules purified from natural source.

## MATERIALS AND METHODS

### Protein purification

Lyophilized lysozyme from *Gallus gallus* was purchased by Sigma-Aldrich. Freeze-dried *Bothrops moojeni* crude venom was purchased from Centro de Extração de Toxina Animais (CETA), Morungaba – SP, Brazil and freeze-dried *Crotalus durissus collineatus* crude venom was purchased from Serpentarium Bioagents of Batatais – SP, Brazil. BmooMP-I was isolated from *B. moojeni* snake venom by cation-exchange chromatography on a CM-FF column (5 mL) followed by reverse-phase using a C-18 column as described by Salvador *et al* (50). CBCol was isolated from *C. d. collilineatus* snake venom by cation-exchange chromatography on a CM-Sepharose column (2 x 20 cm), followed by the dissociation of subunits according to Hendon & Fraenkel-Conrat (51).

### Crystallographic experiments

Crystals were obtained by the hanging-drop vapour diffusion method at 18°C (McPherson, 2009). Lysozyme crystals were grown in crystallization drops composed by 1 μL of protein solution (50 mg/mL) and 1 μL of precipitant solution, equilibrated against reservoir solution of 1.2 M MgCl and 0.1 M Tris HCl, pH 7.6. BmooMP-I crystals were obtained by mixing 0.5 μL protein solution (20 mg/mL) and 0,5 μL reservoir solution equilibrated against a 50 μL reservoir containing 30% (w/v) PEG 400, 0.05 M Tris HCl, pH 8.5, 0.05 M Lithium sulphate and 0.05 M Sodium sulphate (50). CBCol crystals were obtained from a crystallization drop composed of 1 μL of protein (10 mg/mL) and 1 μL of precipitant solution equilibrated against reservoir solution of 2.0 M ammonium sulphate and 0.1 M Tris HCl, pH 9.0 (52).

Crystals were mounted in nylon loops and flash cooled in liquid nitrogen. Diffraction data were collected at the MX2 beamline, Laboratório Nacional de Luz Síncrotron (LNLS, Campinas, Brazil). The BmooMP-I dataset was indexed, integrated, and scaled using HLK2000 (53), all other data with XDS (54). The BmooMP-I, MjTX-I, CBCol and lysozyme structures were solved by molecular replacement with PHASER (48), using the BmooMPalpha-I (PDB code 3GBO), MjTX-I (6CE2), crotoxin basic subunit (2QOG), lysozyme (6G8A) coordinates, respectively. Refinement was performed with phenix.refine (55) and the model was manually built with Coot (56) using the F_obs_-F_calc_ and 2F_obs_-F_calc_ maps. The final models are sanctioned by the validation analysis from Molprobity (57).

### Sequence assignment

#### Crystallography

SEQUENCE SLIDER for natural compounds was used to assign the sequence of lysozyme, BmooMP-I, MjTX-I and CBCol. A protein model with good agreement to the experimental map based on global indicators conforming to the norm was used as input. Rotamers for each one of the 20 possible amino acids were generated using coot function “auto-fit-best rotamer” in each residue in protein model and atoms within 5 Å distance were refined using coot “refine residues” (58). Side-chain atoms had their B-factors set to 30, and their real-space correlation coefficient (RSCC) was calculated against phenix.polder maps (59). Each residue with its omit map from phenix.polder was reviewed manually to assess the possibility of double occupancy and its chemical surrounding. Amino acids with top and non-discriminated RSCC values were included in the probability distribution to generate a theoretical database of sequences to be used against MS data, which will be called PatternLab theoretical database. Possibilities are generated in a peptide basis following expected proteolytic cleavage, which in the cases of this article were for Arg/Lys (trypsin). Images were generated using Coot (58) and Raster3D (60).

#### Mass Spectrometry

*Intact proteins.* 5 μL aliquots of lysozyme previously diluted in 50% acetonitrile (ACN) and 1% formic acid were analysed by direct infusion into the ESI-IT-TOF mass spectrometer (Shimadzu, Kyoto, Japan), under 0.2 mL.min^-1^ constant flow of 50% of solution A&B (solution A: water: DMSO: formic acid (949: 50: 1) and solution B: ACN: DMSO: water: formic acid (850: 50: 99: 1)). The MS interface was kept at 4.5 kV and 275 °C. Detector operated at 1.95 kV. MS spectra were acquired in positive mode, in the 350–1400m/z range and the multiply charged ions were manually deconvoluted for molecular mass determination.

#### Digested proteins

Tryptic digests of the samples were solubilized in 0.1% (v/v) formic acid (solution A) and subjected to nano-ESI-LCMS/MS analysis, using an Ultimate 3000 HPLC (Dionex), coupled to a Q Exactive Orbitrap™ mass spectrometer (Thermo Fisher Scientific). Peptides were loaded on a Trap Column with nanoViper Fitting (P/N 164649, C_18_, 5 mm x 30 μm, Thermo Fisher Scientific) and eluted at a flow rate of 300 nL/min using an isocratic gradient of 4% solution B (100% (v/v) acetonitrile containing 0.1% (v/v) formic acid) for 3 minutes. Thereafter, peptides were loaded on a C18 PicoChip column (Reprosil-Pur®, C18-AQ, 3 μm, 120 Å, 105 mm, New Objective, Woburn, MA, USA) using a segmented concentration gradient from 4–55% B for 30 min, 55–90% B for 1 min, 90% B over 5 min and then returning to 4% B over 20 min at a flow rate of 300 nL/min. Ion polarity was set to positive ionization mode using data-dependent acquisition (DDA) mode. Mass spectra were acquired with a scan range of *m/z* 200–2000, resolution of 70,000 and injection time of 100 ms. The fragmentation chamber was conditioned with collision energy between 29 to 35% with a resolution of 17,500, 50 ms of injection time, 4.0 m/z of isolation window and dynamic exclusion of 10 s. The spectrometric data were acquired through the *Thermo Xcalibur™* software v.4.0.27.19 (Thermo Fisher Scientific).

All MS raw data were compared against the uniprot (61) database with the *PatternLab* software v.4.0.0.84 (62). A database with most probable sequences from the crystallographic evaluation was also used (PatternLab theoretical database). The following search parameters were used: semi-tryptic cleavage products (two tryptic missed cleavage allowed), carbamidomethylation of cysteine as fixed modification and oxidation of methionine as variable modification. Parent mass tolerance error was set at 40 ppm and fragment mass error at 0.02 ppm. Proteins were identified considering a minimum of one fragment ion per peptide, five fragment ions per protein, two peptides per protein and a false discovery identification rate set to 1%, estimated by a simultaneous search against a reversed database.

MS raw data were evaluated using PTM, point mutation and *de novo* algorithms with PEAKS/SPIDER (63, 64). *De novo* analyses were performed with tolerances of 0.01 Da for precursor and fragment ions; methionine oxidation and carbamidomethylation of cysteine were considered variable modifications and fixed, respectively; maximum of 3 PTMs; *de novo* score (ALC%) threshold of 15 and peptide hit threshold (−10logP): 30.0.

#### Phylogenetic analysis

The most probable sequences rendered by crystallographic and MS analysis were used as input in ConSurf (26). For each sample, the 300 most similar sequences (within an interval of 100% and 35%) were used to calculate a percentage of occurrences.

SEQUENCE SLIDER is an open source initiative written in Python3 and distributed through GitHub repository (https://github.com/LBME/slider) and within ARCIMBOLDO (http://chango.ibmb.csic.es/) through the package installer for Python (pip) and Collaborative Computational Project No. 4 (CCP4) (65).

## RESULTS

### Method description

SLIDER integrates crystallography, mass spectrometry and phylogenetic data to assign sequence and its heterogeneity of complex samples (Figure 1). In crystallography, it builds the rotamer with the best fit in electron density map for each one of the 20 natural amino acids in each residue position using the software Coot (56). Their side chain atoms are used to estimate the agreement between their theoretical electron density with the observed one through calculation of their real-space correlation coefficient (RSCC) using phenix.polder (59), a methodology that creates an omit map excluding the bulk solvent around the omitted region. Alternatively, instead of searching for all 20 amino acids, restrictions may be extracted from Genetic information (homologues) or from MS data (illustrated in Figure 1 by arrow “Restricting seq hyp” connecting Genetic information or Mass spectrometry data ellipses to Structural data ellipse). By residue position, single or multiple amino acid possibilities may accommodate in the observed map, measured herein with RSCC (Structural data ellipse in Figure 1).

**Figure 1.**
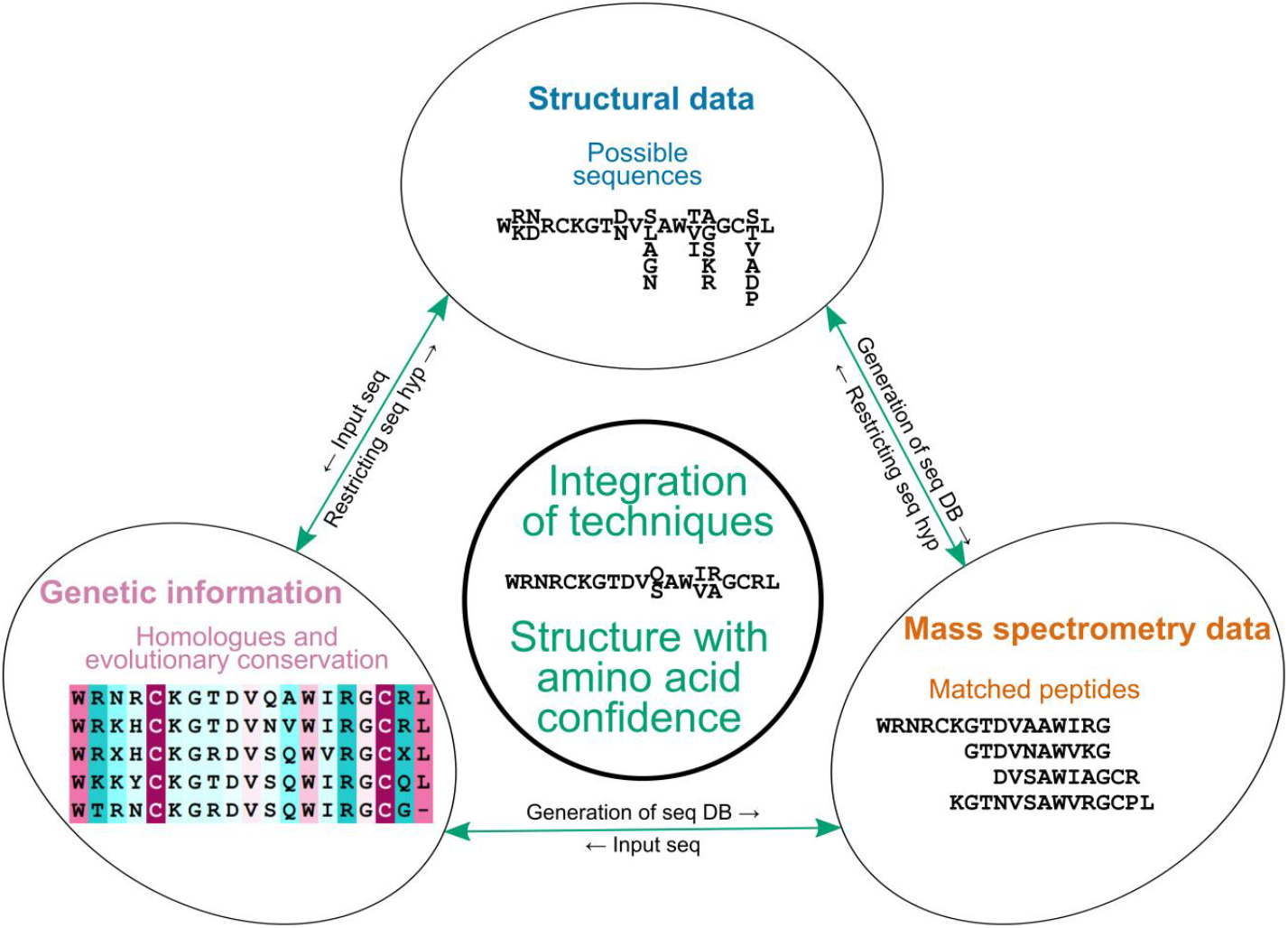
Scheme of how SEQUENCE SLIDER integrates structural, mass spectrometry and phylogenetic data for sequence assignment.

From the single and multiple possibilities of amino acids in each residue position, a theoretical database of sequences in FASTA format can be built containing all this variability (arrow “Generation of seq DB” connecting Structural data ellipse to Mass spectrometry ellipse in Figure 1). To reduce the exponential growth of the combination of amino acid sequences, possibilities are generated in a peptide basis following expected proteolytic cleavage. This database also incorporates the reversed order of each sequence as a baseline to estimate true and false positives by the available mass spectrometry software. Herein, we used PatternLab for Proteomics, the Brazilian integrated computational environment for analysing shotgun proteomic data (62). Alternatively, the MS/MS spectrum can be searched against *in-silico* digested sequences of proteins available in the public databases (NCBI (66) and uniprot databases (61)) (illustrated in Figure 1 by arrow “Generation of seq DB” connecting Genetic information ellipse to Mass spectrometry data ellipse). Additionally, possibility of PTM and homology peptide mutations can be evaluated with *de novo* sequencing, which is available in PEAKS/SPIDER (63, 64) (arrow “Restricting seq hyp” connecting Mass spectrometry ellipse to Structural data ellipse in Figure 1).

Estimation of sequence conservation and the amino acid frequency of each residue position based on homologue sequences (illustrated by Genetic information ellipse in Figure 1) is calculated in SLIDER by ConSurf, a bioinformatics tool to estimate evolutionary conservation based on phylogenetic relations between homologues (26, 27), giving as input sequence the most probable sequence using results from previous techniques (arrow “Input seq” connecting either Structural data or Mass spectrometry data ellipses to Genetic information ellipse in Figure 1). For some residues, information gathered may point to a single residue, for others, more than one amino acid may be possible, which can be built with partial occupancy or even as an unknown residue (described as UNK). Double occupancy of different amino acids is allowed in the deposition and is known as microheterogeneity (67). In summary, SLIDER balances different sources of information to either assign a single amino acid if unambiguously identified or multiple possibilities or unknown when the evidence is not conclusive, in an approach that drives sequence hypothesis from data and prevents bias propagation from information known *a priori.*

### Lysozyme

Lysozyme is one of the most studied structures, as it is readily obtained (e.g. from egg white) and crystallized, totalling over 5000 entries in the PDB (12). Still, these represent but a fraction of the thousands of sequences registered for different species in multiple isoforms in (NCBI (66) and uniprot databases (61)). As the structure and sequence of this enzyme have been extensively characterized, we chose it to calibrate the SLIDER methodology. Using the NCBI database against the MS spectrum of *Gallus gallus* lysozyme yields a high coverage, 117 residues out of 129 (Figure 2A). We collected diffraction data to a resolution of 1.9 Å and concluded model-building and refinement once regression between experimental and calculated data (R_work_/R_free_ of 18.5/21.7%) and stereochemistry (Molprobity score is 1.71, which refers as 87^th^ percentile) were conventionally acceptable.

**Figure 2.**
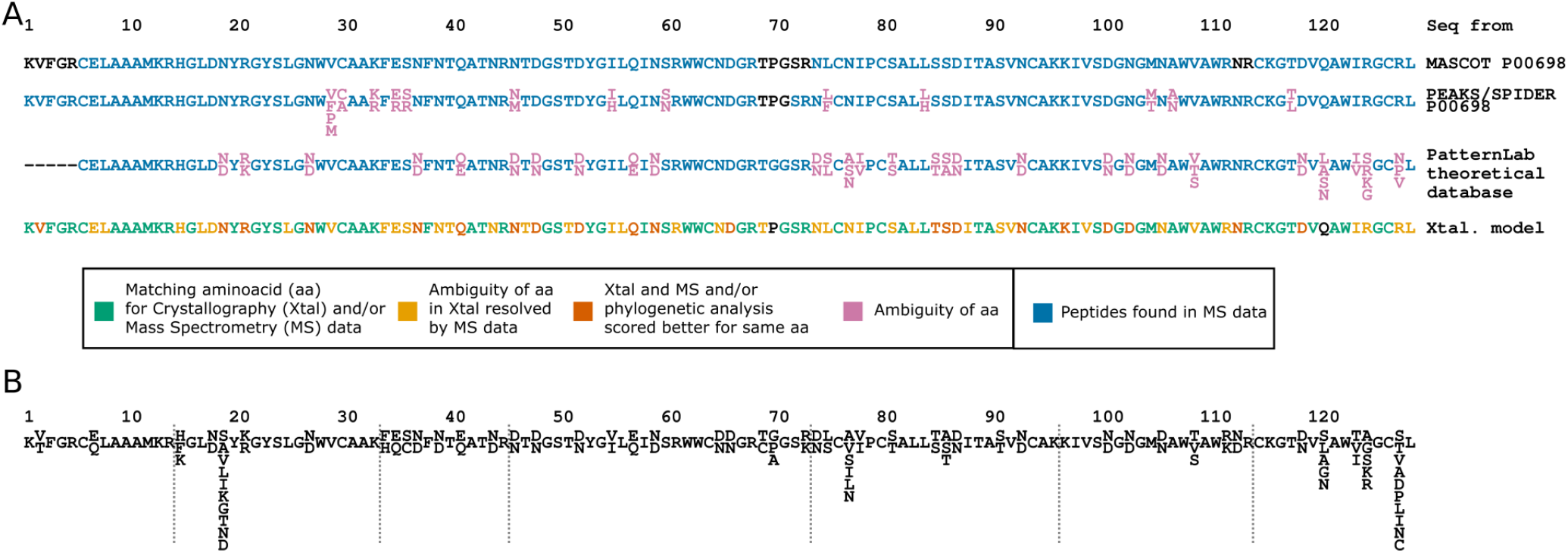
Sequence assignment of lysozyme. (A) Mass spectrometry data is used against an NCBI database of lysozyme sequences, against PEAKS/SPIDER evaluation and against PatternLab theoretical database and the sequence from the crystal model. (B) Theoretical database constructed by SEQUENCE SLIDER

For the SLIDER sequence evaluation using crystallographic data based on the RSCC calculations, seven residues from three groups that share similar scattering and shape were ambiguous: A) valine and threonine; B) asparagine, aspartic acid and leucine; C) glutamic acid and glutamine. On the other hand, analysis of the chemical environment may restrict possibilities as hydrophobic residues would not undergo hydrogen bonds with water molecules or polar atoms and would be most probably buried. Mostly exposed to the solvent, 28 residues (21.7%) had possibilities restricted to the hydrophilic group, while mainly buried, 13 residues (10.1%), were restricted to hydrophobic (Figure 2B). Hence, we generated 9648 sequence hypotheses involving the residues that were favoured by the electron density, as evaluated by the RSCC, and the previous physicochemical restrictions (Figure 2B).

In PEAKS/SPIDER application against the lysozyme MS data and uniprot database restricted to *Gallus gallus* sequences, we found 845 peptide-spectrum matches with a false discovery rate of 0.2%, which is the expected percentage of peptides where a reversed sequence was selected. Single sequence obtains good coverage, its sequence (P00698) had 3, 113, 12 and 1 residues with none, one, two and four amino acid possibilities, respectively (sequences shown in PEAKS/SPIDER description of Figure 2A). SLIDER sequences evaluated with PatternLab had more variation, 5, 96, 23, 3 and 2 residues with none, one, two, three and four amino acid possibilities, respectively (sequences shown in PatternLab theoretical database of Figure 2A). Comparing both evaluations, 83 residues (64.3%) were given same assignment and 46 (35.7%) were different.

Most residues in the lysozyme crystallographic model clearly favoured a single amino acid, discriminated by comparison of the top and second best RSCC scoring, a statistical indicator we called Δcontrast. These calculations were confirmed by manual evaluation of omit maps, by MS and/or by phylogenetic analysis (Figure 2A and Supplementary Table 1). Empirically, we established a Δcontrast of RSCC of 3.0% as an appropriate threshold to distinguish an amino acid from the rest. Ambiguity of 39 residues (30.2%) were resolved by MS spectrum (orange residues in Figure 2A and Supplementary Table 2) and ambiguity of the remaining 21 residues (16.3%) was excluded with phylogenetic analysis (Supplementary Table 3).

The Isomorphous template (PDB code 6G8A) used to phase our lysozyme crystallographic model has the presumably correct sequence. It was confirmed by measuring the intact mass of our sample using mass spectrometry as the molecular mass determined of 14306.31 ± 2.81 Da is in agreement with the theoretical (14303.88 Da). PEAKS/SPIDER found 113 residues correctly (87.6%), 13 correct but had other amino acid possibilities (10.0%) and 3 were not found (2.4%). PatternLab with theoretical SLIDER database found 95 residues correctly (73.6%), 27 correct but had other amino acid possibilities (20.9%), 2 wrong (1.6%) and 5 were not found (3.9%). Pro70 and Gln121 were not seen in PatternLab theoretical database, instead a Gly70 and Leu/Asn/Ser/Ala/Gly121 were observed (Supplementary Table 4). In fact, the residue Gln121 could not have been found by PatternLab as this possibility was not within generated sequences (Figure 2B), as its electron density omit map supports a Ser121 (Figure 3). Also, the omit map for residue 70 supports a Gly (Figure 3). Considering the PEAKS/SPIDER amino acids, residues 70–72 were not seen, but Gln121 was observed. Therefore, we used the lysozyme data to calibrate SEQUENCE SLIDER, to establish the minimum Δcontrast to distinguish a particular side chain, to integrate MS and genetic information to previous assignments.

**Figure 3.**
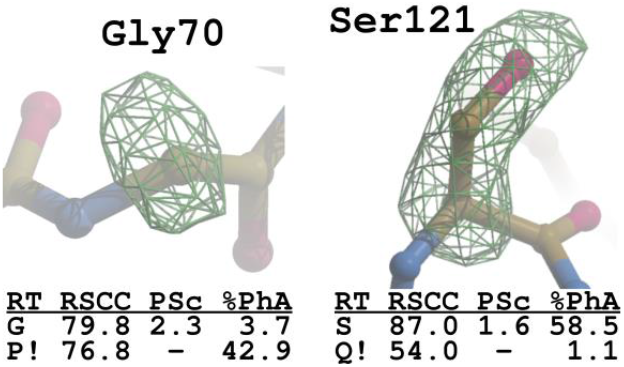
Electron density of omit map of different lysozyme residues in comparison to the PDB code 6G8A with mass spectrometry and phylogenetic analysis statistics. RT stands for residue type; RSCC for real-space correlation coefficient, PSc for Primary Score, %PhA frequency from phylogenetic analysis; and ! for SLIDER choice for given amino acid in a single letter code.

### Metalloproteinase BmooMP-I

The first novel crystallographic and mass spectrometry data that SLIDER in natural compounds was applied is the metalloproteinase BmooMP-I from *B. moojeni,* a protein able to express fibrinogenolytic and gelatinase activities that is important in the prey’s immobilization and digestion whose biological article was already published by us but did not include phylogenetic analysis (50).

The MS data analysis in a typical strategy using a single digestion procedure, trypsin, against NCBI database of Bothropic genus reached a coverage of 131 residues (65.5%) out of 200. The sequences with highest scores were five bothropic metalloproteinases with 78–57 residues matching in MS spectrum (Figure 4A). With 34.5% of the sequence missing, this was an ideal case for application of the SLIDER methodology to assign sequence heterogeneity.

**Figure 4.**
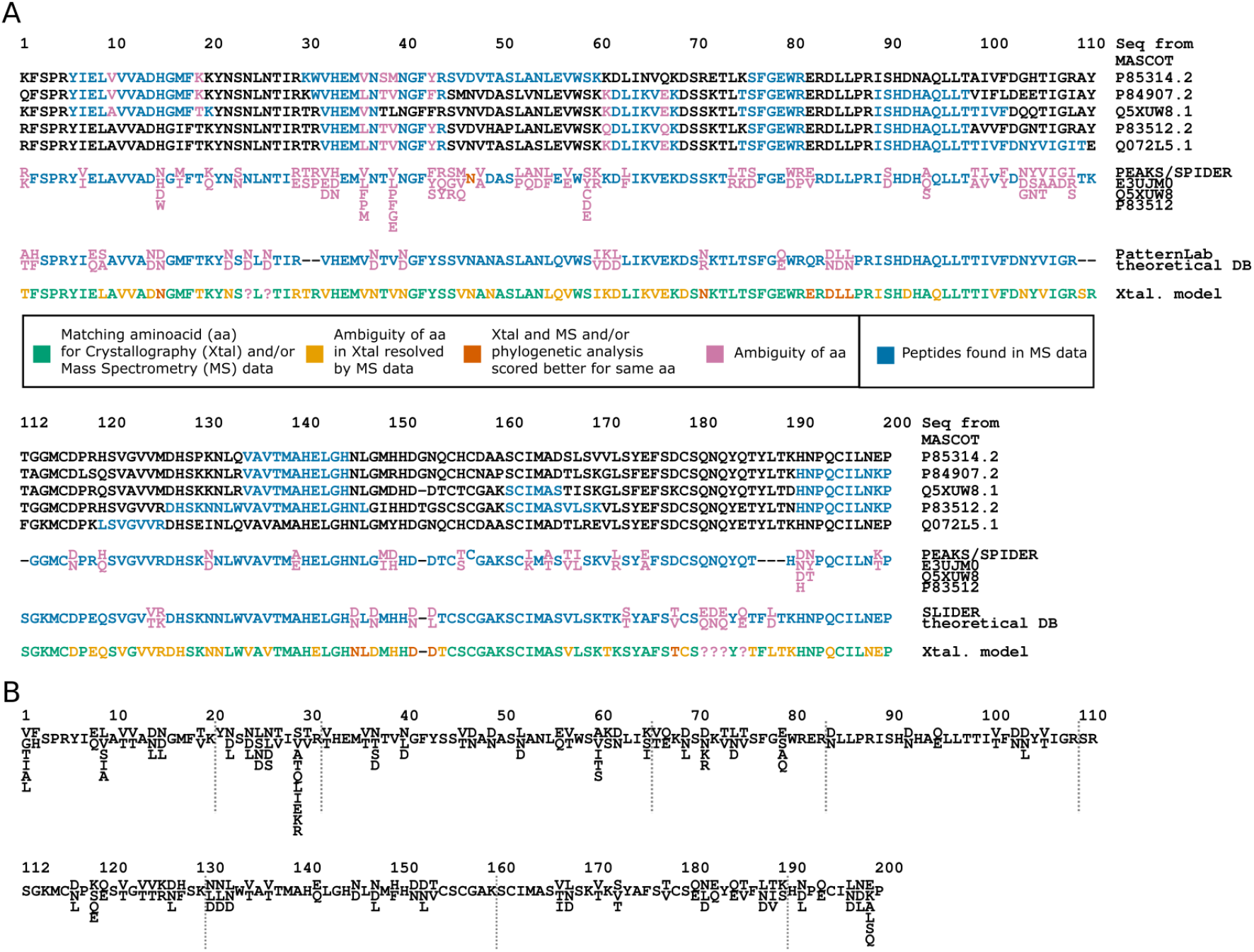
Sequence heterogeneity modelling of BmooMP-I. (A) Mass spectrometry data against an NCBI database with Bothrops genus sequences, against PEAKS/SPIDER evaluation and against PatternLab theoretical database and the sequence from the crystal model. (B) Theoretical database constructed by SEQUENCE SLIDER. Adapted from (50).

The BmooMP-I crystallographic model was deposited under PDB code 6X5X, at 1.92 Å resolution with excellent agreement to the data (R_work_/R_free_ of 16.0/18.1%) and to the stereochemistry (Molprobity score of 0.89, which refers as 100^th^ percentile). The side chain possibility was restricted to hydrophobic residues evaluating its surrounding for 19 residues (9.5%), mostly buried (Figure 4B). Hydrogen bonds were found for 33 residues (16.5%), mainly exposed (Figure 4B). Based on these observations, the amino acids and peptides breakage used for theoretical database generation are illustrated in Figure 4B and summed 419268 sequences.

The theoretical peptides with spectrum matching the MS data from PatternLab are shown in Figure 4A in PatternLab theoretical DB (database) description with certainties coloured in green and uncertain residues in magenta. Using the MS data and uniprot database restricted to taxonomy Serpentes with PEAKS/SPIDER, we found 1900 peptide-spectrum matches with a false discovery rate of 1.4%. In order to achieve good sequence coverage, we merged peptides from three sequences, all bothropic metalloproteases, and 6, 137, 42, 12, 2, 2 and 1 residues had none, one, two, three, four, five and six amino acid possibilities, respectively (sequences shown in PEAKS/SPIDER description of Figure 4A). On the other hand, PatternLab theoretical sequence strategy had 6, 164 and 32 residues having none, one and two amino acid possibilities, respectively (sequences shown in PatternLab theoretical DB description of Figure 4A). Comparing these two MS evaluations, 103 residues (51.5%) were given same assignment and 97 (48.5%) were different.

In the crystallographic model, out of 200 residues, 121 (60.5%) favoured a single, well discriminated assignement, confirmed by MS and/or phylogenetic analysis (coloured green in Figure 4A and shown in Supplementary Table 5). More than one possibility was found for 60 residues (30.0%), but their ambiguity was resolved by MS (coloured orange in Figure 4A and shown in Supplementary Table 6). Using the phylogenetic analysis from ConSurf and 300 most similar sequences, it was resolved the amino acid 15, 71 and 146 as asparagine, 82 as glutamic acid, 84, 152 and 153 as aspartic acid, 85, 86 and 147 as leucine, and 178 as threonine (Supplementary Table 7). From the group of residues possessing atomic similarity, the mass spectrometry data and phylogenetic analysis were not enough to distinguish Asn/Asp24 (residue 24 which could be asparagine or aspartic acid), Asn/Asp26, Gln/Glu181, Asn/Asp182, Glu/Gln183 and Gln/Glu185 (coloured magenta in Figure 4A, illustrated in Figure 5 and shown in Supplementary Table 8). As two amino acids were equally possible for these previous residues, SLIDER identified this heterogeneity by leaving them partially occupied in the model. Therefore, SLIDER was able to conclude a single amino acid possibility for 192 residues and two possibilities for the remaining 8.

**Figure 5.**
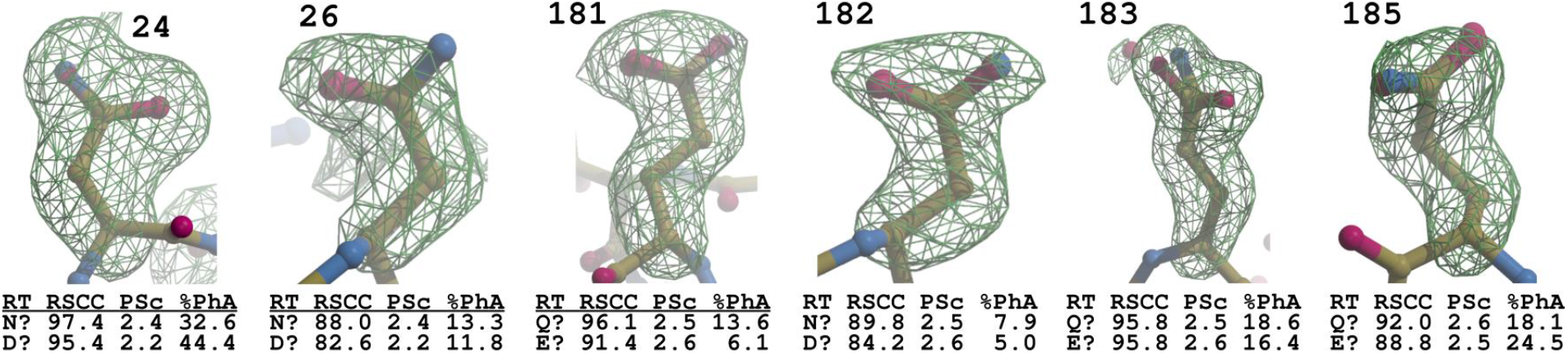
Electron density of omit map with possible residues given crystallographic, mass spectrometry and phylogenetic analysis statistics in metalloproteinase BmooMP-I. RT stands for residue type; RSCC for real-space correlation coefficient, PSc for Primary Score, %PhA frequency from phylogenetic analysis; and ! for SLIDER choice for given amino acid in a single letter code.

### PLA_2_-like protein MjTX-I

The MjTX-I is an unique PLA_2_-like protein due to an insertion in its C-terminus and few mutations that decrease its myotoxicity compared to other PLA_2_-like proteins (68). The NCBI database of protein sequences contains various bothropic PLA_2_s and using them against MjTX-I MS data, five bothropic PLA_2_-like proteins sequences have 114–92 matched residues out of 122, respectively (Figure 6A). The ten matched residues being different (coloured in magenta in Figure 6A) evidence presence of isoforms and SEQUENCE SLIDER should resolve these uncertainties.

**Figure 6.**
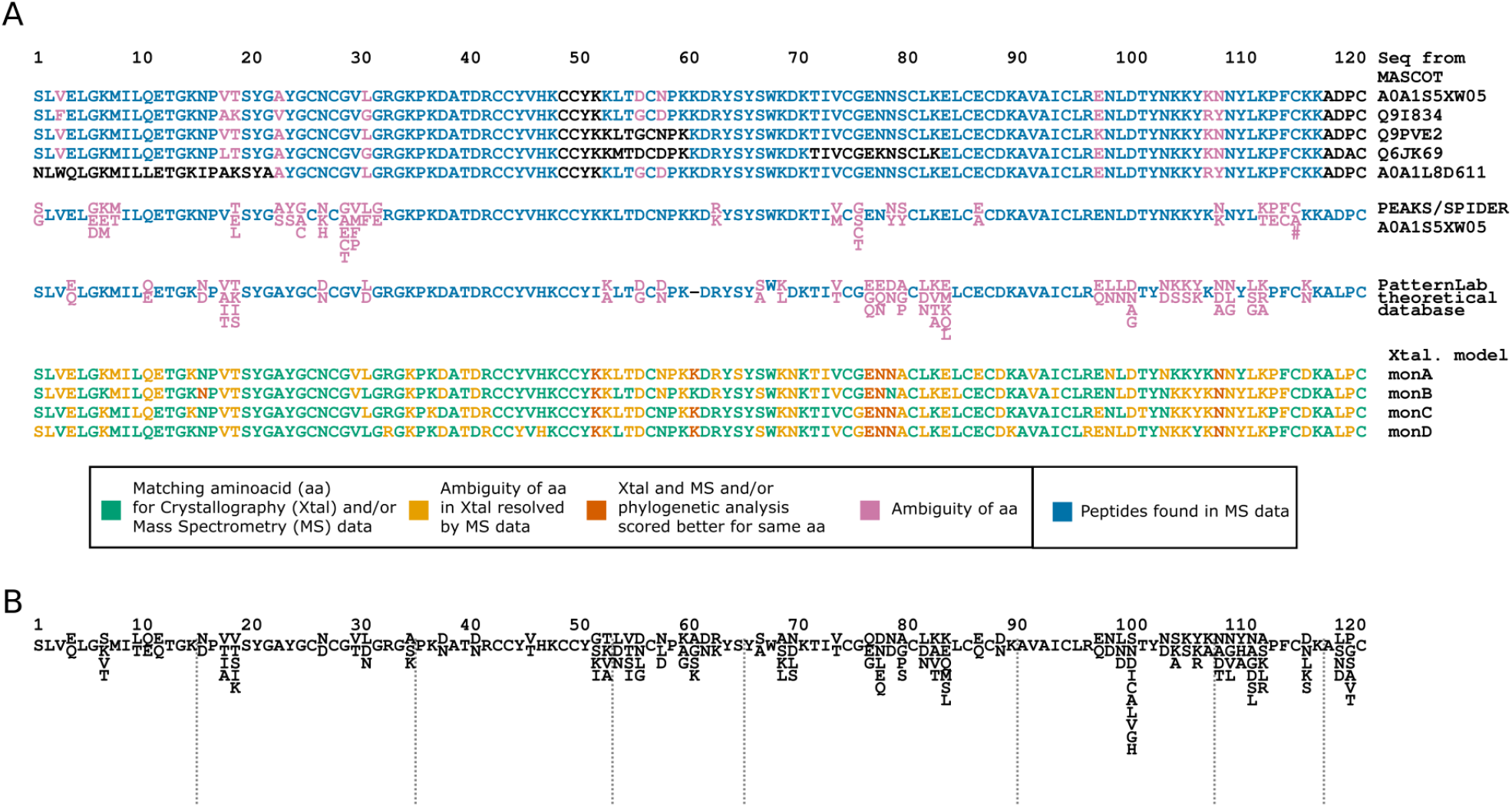
Sequence heterogeneity modelling of MjTX-I. (A) Mass spectrometry data against an NCBI database with Bothrops genus sequences, against PEAKS/SPIDER evaluation and against PatternLab theoretical database and the sequence from the crystal model. (B) Theoretical database constructed by SEQUENCE SLIDER.

The MjTX-I crystallographic structure was deposited under the PDB code 7LYE at 1.85 Å resolution with good agreement to the data (R_work_/R_free_ of 19.0/23.0%) and to the stereochemistry (Molprobity score of 1.72, which refers as 83^rd^ percentile), whose biological article was already published by us (69). Four chains were present in the asymmetric unit and required SLIDER to evaluate them separately to allow possibility to distinguish isoforms. For the MjTX-I calculations, side chains of 13 mostly exposed residues (10.7%) were surrounded by charged atoms and/or had the presence of hydrogen bond(s) and had their possibilities restricted to hydrophilic residues (Figure 6B). The side chains of buried 5 residues (4.1%) were surrounded by hydrophobic residues and had their possibilities to such chemical character (Figure 6B). The hypotheses of sequences extracted by SLIDER are shown in Figure 6B and summed 2236248.

In PEAKS/SPIDER application against the MjTX-I MS data and uniprot database restricted to Serpentes taxonomy, we found 1489 peptide-spectrum matches with a false discovery rate of 1.5%. A single sequence was enough to obtain good sequence coverage, BomoTx (A0A1S5XW05) had 1,98, 15, 6, 2 and 1 residues with none, one, two, three, four and five amino acid possibilities, respectively (sequences shown in PEAKS/SPIDER description of Figure 6A). Theoretical sequences had more options, 1, 89, 20, 8, 4 and 1 residues with none, one, two, three, four and five amino acid possibilities, respectively (sequences shown in PatternLab theoretical DB of Figure 6A). Comparing both evaluations, 71 residues (58.2%) were given same assignment and 51 (41.8%) were different. Moreover, in residue number 116, a deletion was discovered by PEAKS/SPIDER evaluation, showing MjTX-I isoforms contain at least one protein of 121 residues and another with 122, therefore SLIDER restricted database works well to restrict possibilities within crystal structure, but fail to capture overall isoforms complexity.

Most of the residues in the crystallographic model were resolved, which was confirmed by either MS or phylogenetic analysis (coloured green in Figure 6A under Xtal. model description and shown in Supplementary Table 9). The residues coloured in orange in Figure 6A (Supplementary Table 10) had no single discrimination based on RSCC but were resolved based on MS. The side chain choice of 23 residues, out of the 488 considering the tetramer, were based on the conjunction of RSCC, MS and phylogenetic statistics (vermilion residues in Figure 6A and Supplementary Table 11). We did not find clear evidence that the monomers had a different sequence, the only variation in fact was lack of electron density for the residues Lys7C–D, Lys52D, Asp56C, Lys60C, Lys61A/C–D, Ser67D, Lys69A–D, Glu77D, Lys83B, Glu84C, Lys105C, Lys106B–D, Asn109C, Leu112A/C–D, Lys113A–C, Pro121C. Also, we generated omit maps in C-terminal region to consider the deletion in residue 116 present in the PEAKS/SPIDER analysis, although electron density did not support this deletion.

By applying this novel methodology, we could identify that the sequence obtained was not yet seen in databases, its highest identity observed to other proteins in PDB is 92.6% with MtTX-II (PDB code 4DCF) with the difference in 9 residues of 121. Compared to the crystallographic model of MjTX-I complexed with suramin (PDB code 6CE2), sequence differences are found for Lys-Thr19, Asp-Ser67, Glu-Asn78, Pro-Ala80, Lys-Leu100, Gly-Asp101, an insertion in 107, Arg-Lys108, Val-Asn110, Gly-Ala119, Arg-Leu120 and Asp-Pro121 and undistinguished residues, Asn-Asp56, Gln-Glu84 and Asp-Asn109 (Figure 7). Compared to the non-redundant protein sequences in BLAST, BomoTx from *B. moojeni* (code A0A1S5XW05.1) shares 96.7% of identity with 4 residues different.

**Figure 7.**
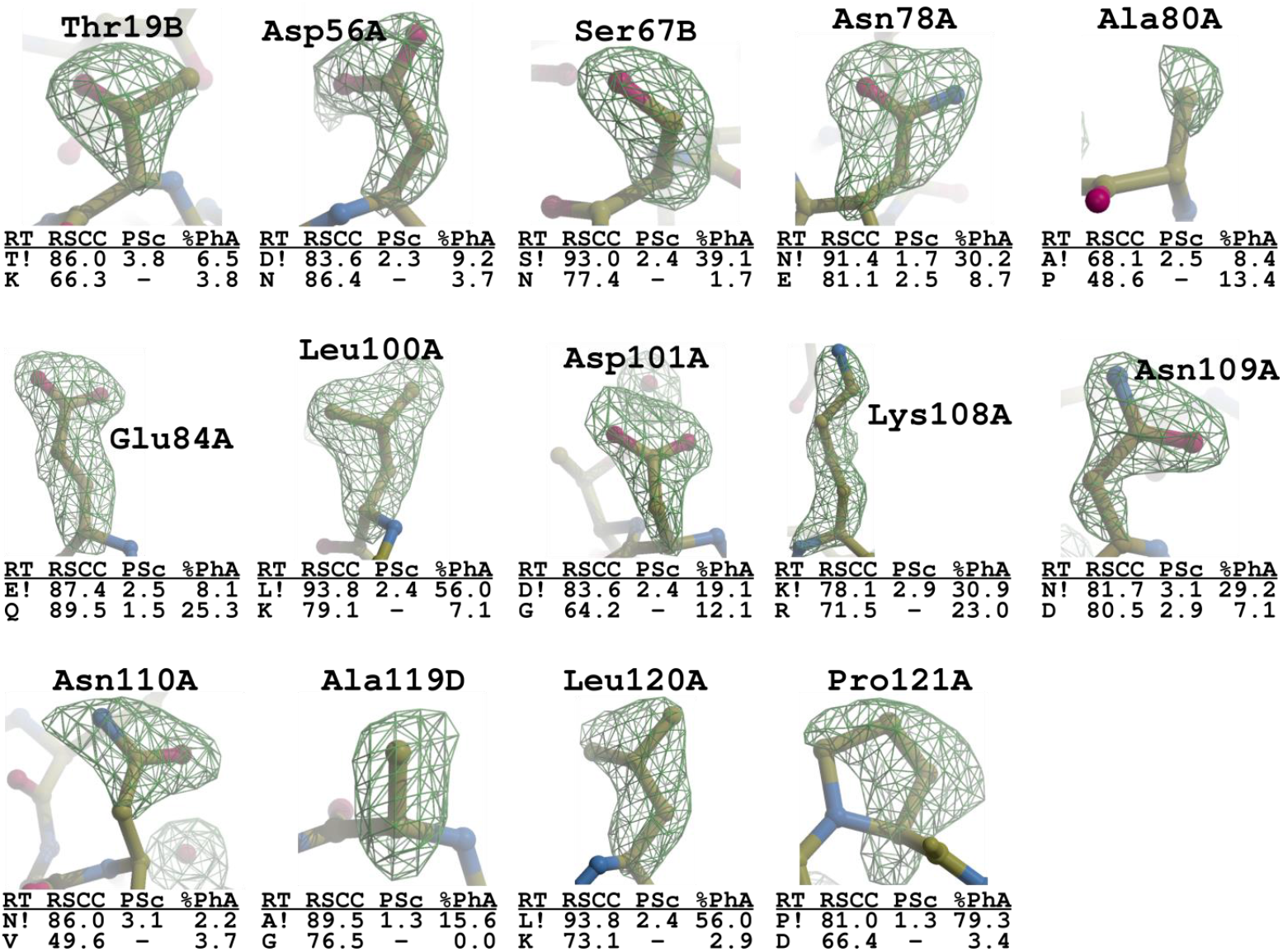
Electron density of omit map of different MjTX-I residues in respect to its previous crystallographic model (PDB code 6CE2) with mass spectrometry and phylogenetic analysis statistics. RT stands for residue type; RSCC for real-space correlation coefficient, PSc for Primary Score, %PhA frequency from phylogenetic analysis; and ! for SLIDER choice for given amino acid in a single letter code.

### Crotoxin CBCol

The crotoxin is one of the most studied snake venom toxins, due to its high abundance in Crotalus venom, to its toxicity and pharmaceutical potentiality (70). The database of protein sequences contains various isoforms of the crotoxin, most of them being from *Crotalus durissus terrificus.* Against the MS data from *Crotalus durissus collilineatus,* the NCBI database of snakes gave a maximum of 114 matched residues out of 122 for the CBc and CBa_2_ (both from *C. d. terrificus),* with 8 amino acids being different (magenta in Figure 8A) evidencing isoforms presence in the purified sample. Applying SEQUENCE SLIDER to the MS, crystallographic and phylogenetic data should improve even more this coverage and resolve uncertainty.

**Figure 8.**
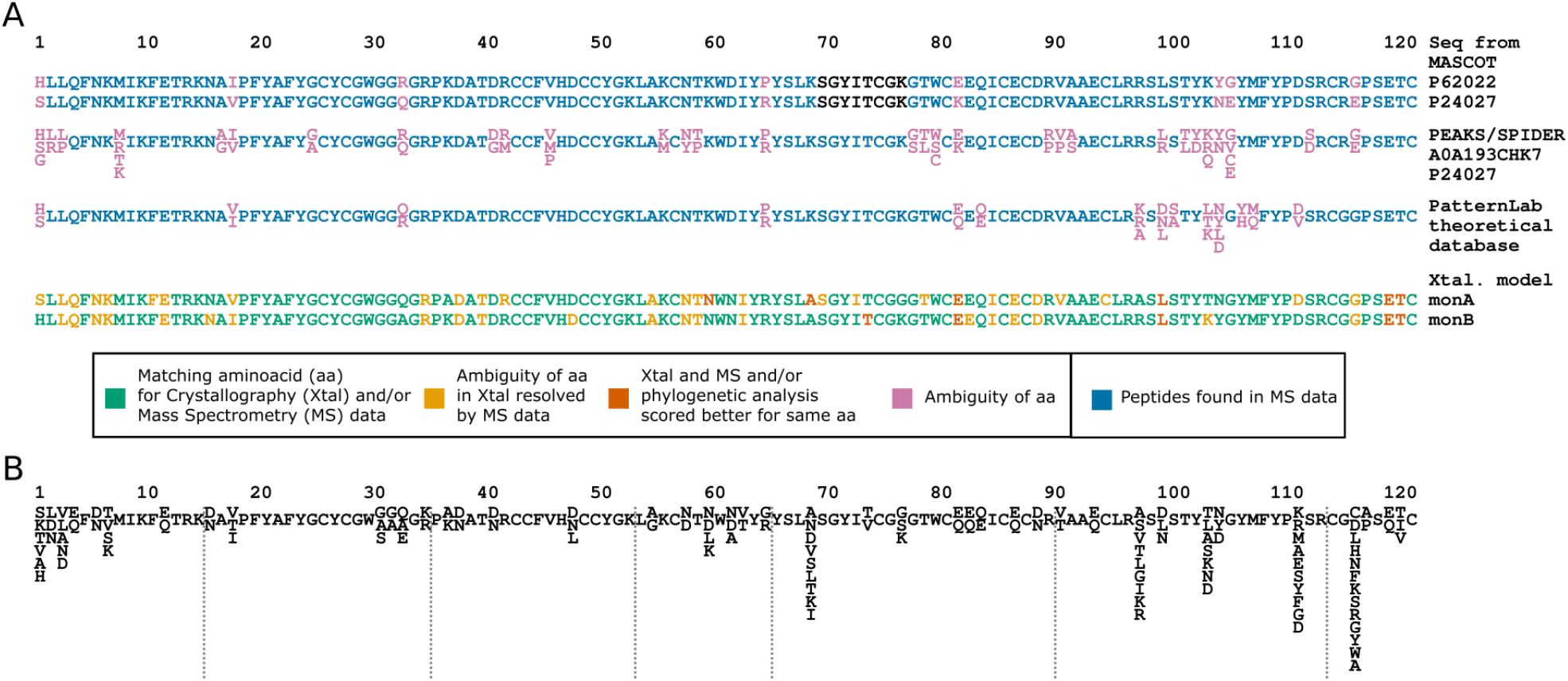
Sequence heterogeneity modelling of CBCol. (A) Mass spectrometry data is used against an NCBI database with Crotalus genus sequences, against PEAKS/SPIDER evaluation and against PatternLab theoretical database and the sequence from the crystal model. (B) Theoretical database constructed by SEQUENCE SLIDER.

We finished the crystallographic model of crotoxin crystallized from *C. d. collineatus* at 1.9 Å resolution with good agreement with the data (R_work_/R_free_ of 20.2/24.3%) and with the stereochemistry (Molprobity score of 1.6, which refers as 91^st^ percentile). Herein, two chains were present in the asymmetric unit and required SLIDER to evaluate them separately. Possibilities of 14 mostly exposed residues (11.5%) were reduced based on hydrophilic residues and the presence of hydrogen bond(s) (Figure 8B). Buried 4 residue had their possibilities restricted to hydrophobic amino acids. The hypotheses of sequences extracted by SLIDER are shown in Figure 8B and summed 27876.

In PEAKS/SPIDER application against the CBCol MS data and uniprot database restricted to Serpentes taxonomy, we found 1994 peptide-spectrum matches with a false discovery rate of 1.8%. Two sequences (CBa_2_ from *Crotalus durissus terrificus* and Cb3 from *Crotalus basiliscus)* had to be considered to obtain a better distribution of the amino acid possibilities that had 5, 92, 24, 4 and 2 residues with none, one, two, three and four amino acid possibilities, respectively (sequences shown in PEAKS/SPIDER description of Figure 8A). Theoretical sequences evaluated with PatternLab had less variation, 1, 107, 11, 3 and 1 residues with none, one, two, three and four amino acid possibilities, respectively (sequences shown in PatternLab theoretical database of Figure 8A). Comparing both evaluations, 89 residues (73.0%) were given same assignment and 33 (27.0%) were different. Good agreement is seen in two methodologies.

Most of the residues of the crystallographic model were resolved by a single possibility confirmed by either MS or phylogenetic analysis (coloured green in Figure 8A and Supplementary Table 12). Out of the dimer (244 residues), 47 residues (19.3%) had no single discrimination based on RSCC but were resolved based on MS (orange residues in Figure 8A and Supplementary Table 13). The side chain choice of 9 residues (3.7%) were based on best RSCC confirmed by phylogenetic analysis (vermilion residues in Figure 8A and Supplementary Table 14). Residue 102 had matches for Asp, Asn and Leu in MS spectrum, but only Leu was observed in remote homologous sequences with 47.5% frequency, therefore this last residue was chosen. Residue 7A/B and 112A had their side chain atoms removed due to lack of clear electron density.

By applying this novel methodology, we could identify that residues 1, 18, 33, 37, 77, 98, 104 and 105 are different when chain A is compared to B (Figure 9) as SLIDER was able to distinguish these two isoforms of crotoxin and better characterize the crystal sample. These differences would have probably been neglected in a conventional structure elucidation practice. The monomer A is similar to CBa_2_ isoform from *C. d. terrificus* (PDB code 2QOG) sharing 92.6% of identity and the monomer B is comparable to CBd (PDB code 6TMY) from this same specie with 97.5% of identity. In fact, crotoxin complex consists of the noncovalent association of a crotapotin (CA). acid component of one of its isoforms CA1-4, product of different PTM events; with a basic CB, one of its isoforms CBa_2_, CBb, CBc and CBd, consequence of expression of different mRNA (23). CA/CBa_2_ results in a less stable complex with a higher enzymatic activity and lower toxicity than the other associations. Our model has only basic components, CBa_2_ and CBd. These two monomers are similar, their RMSD considering all Cαs is 1.5 Å, which is improved to 0.4 Å rejecting 23 Cαs. The mostly varied region is comprehended by loop residues 58–62 that participates in the crystal packing with residue 1 in monomer CBa_2_ and with residue 77 in monomer CBd, supporting hypothesis that isoform preference in one of the position in the dimer is favoured by the different crystal packing. Moreover, in monomer CBd, Trp61 is attracted by His1 of helix 1, while in CBa_2_, Trp61 is pointed towards hydrophobic residues in antiparallel helices 2 and 3, therefore the point mutation in residue 1 seems the responsible for largest seen difference between two monomers.

**Figure 9.**
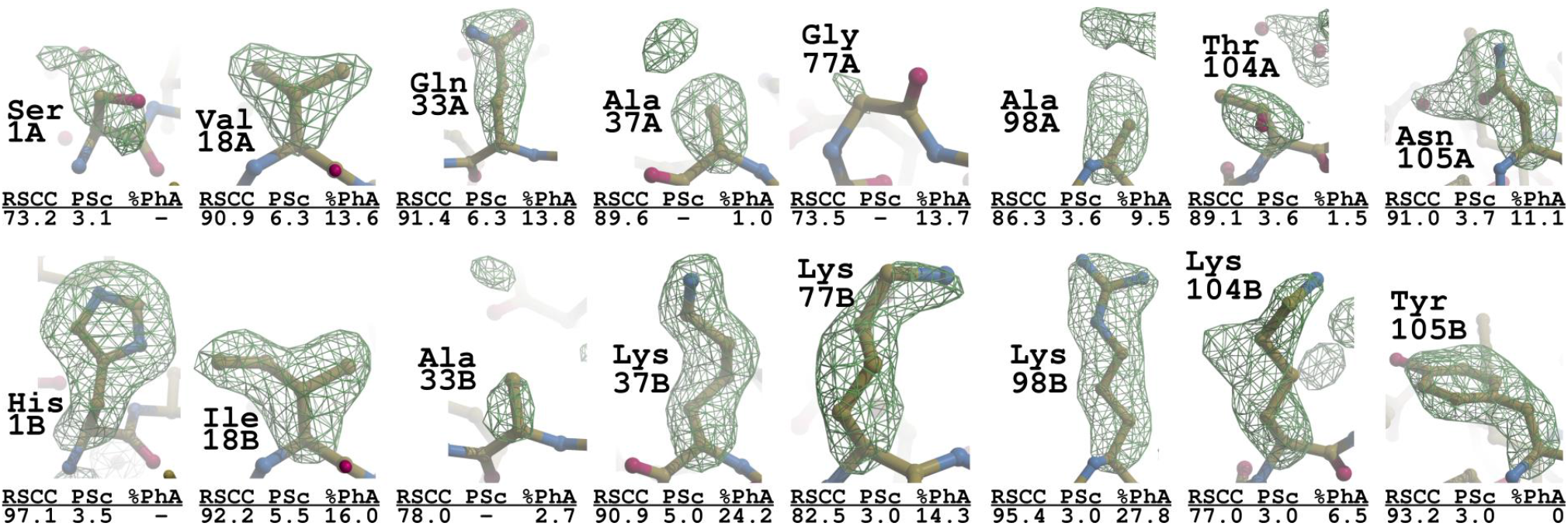
Electron density of omit map of different CbCol residues among its monomers with mass spectrometry and phylogenetic analysis statistics. RT stands for residue type; RSCC for real-space correlation coefficient, PSc for Primary Score, %PhA frequency from phylogenetic analysis; and ! for SLIDER choice for given amino acid in a single letter code.

## DISCUSSION AND CONCLUSION

The coexistence of isoforms with similar physicochemical properties may prevent complete purification of proteins from natural source. This is typical in toxinology, as toxins are usually obtained from extracted venom and they are characterized by one of the most rapid evolutionary divergence and variability in any category of proteins (15). In the absence of atomic resolution data, the elucidation of the structure from those samples requires conjunction of biophysical complementary methods to account for sequence heterogeneity.

Herein, we repurposed the amino acid evaluator SEQUENCE SLIDER (46) from phasing to integrate crystallography, MS and phylogenetic analysis aiming the characterization of these complex samples. We use the agreement between theoretical electron density with the observed one (59) to foresee which amino acids are possible in each residue position. From this probability distribution, we generate a theoretical sequence database that is used against MS data to identify amino acids from the peptides present in sample (62). Algorithms searching for point mutations in homologues and PTM can also be integrated (63, 64). Our approach integrates those instances successfully exploited by Diemer *et al.,* who obtained a preliminary sequence from crystallography and then corrected it by MS data (43) and Guo *et al., who* estimated residue conservation and frequency using the phylogenetic analysis (26), to distinguish aspartic acid from asparagine and glutamic acid from glutamine using homolog sequences in regions where no MS data revealed the correct amino acids. We extend the integrative framework to estimate in an automated program the probability of establishing a single assignment or acknowledge an existing indetermination with the available experimental data and prior knowledge.

In the absence of atomic resolution, the small electron scattering difference between O, C and N, renders three groups of residues indistinguishable in the crystallographic omit electron density maps: A) valine and threonine; B) asparagine, aspartic acid and leucine; C) glutamic acid and glutamine (Figure 10) (43, 44). Depending on the chemical environment of such residues, the ambiguity of groups A and B may be decreased by hydrophobic or hydrophilic residues (blue and orange description in Figure 10) (43). Such restrictions were essential to reduce the number of sequences in the theoretical database that, for the MjTX-I case per example, reached more than two million. Moreover, despite the stability of the protein structure and their multiple disulphide bonds, large and hydrophilic residues exposed to the solvent may have side chain atoms flexible with no clear electron density being difficult to distinguish from small residues, such as Ala and Ser (43). For the MjTX-I model, 28 out of 488 residues had their side chain omitted. On the other hand, in a high-resolution MS equipment, such as the orbitrap detector used herein, the resulting spectrum allows the distinction of these residues, the only residues whose charge and molecular mass are the same are leucine and isoleucine (Figure 10) (43, 44). Given the different ambiguities of crystallography and MS, the tandem of the two structural techniques, informed by phylogenetic analysis, should enhance identification of single sequences present in the purified simple, while realistically accounting for lingering indetermination (43).

**Figure 10.**
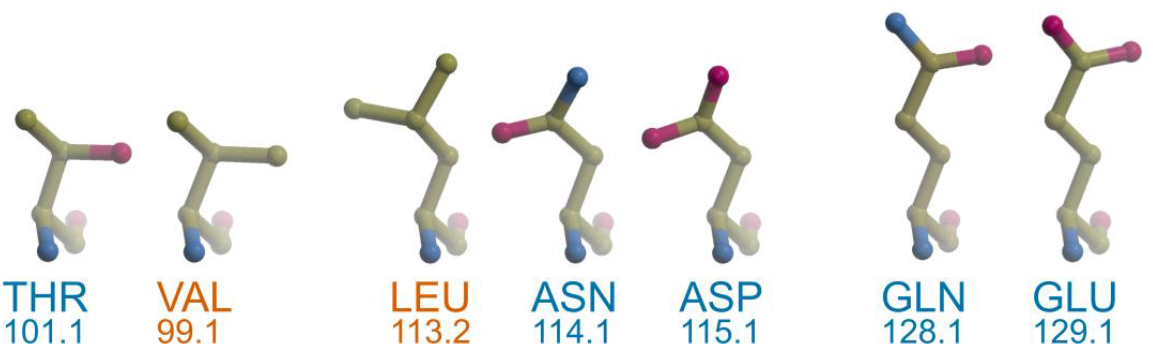
Three groups of residues sharing atomic structure with their molecular mass, coloured in blue and red hydrophilic and hydrophobic residues, respectively.

These assumptions were confirmed by the datasets collected herein, as most of the residues were distinguished either by crystallography itself or with its conjunction to genetically informed MS. We compared two different strategies, first evaluating a theoretical database of sequences against MS data with PatternLab (62), and second using the MS spectrum to sequence *de novo* and against a uniprot sequences considering possible point mutation and PTM with PEAKS/SPIDER (63). Considering the matched spectrum of peptides, PatternLab theoretical database had less possibilities than PEAKS/SPIDER for BmooMP-I and CbCol cases, while lysozyme and MjTX-I were the opposite. In this last data, the generality of PEAKS/SPIDER considering less restriction was able to find peptides in the C-terminus of MjTX-I supporting the presence of at least two different sized proteins, one containing 121 and another with 122. While our approach in PatternLab is dependent of crystallographic omit maps, PEAKS/SPIDER run may discover potential deletion or insertion and is run as an independent analysis. The agreement between the two strategies ranged from half to three quarters of each residue assignment depending on the case. The only possible appraisal in respect to the true sequence is for the lysozyme data, PEAKS/SPIDER, with its calculated lower false discovery rate of 0.2%, was slightly better than PatternLab that had 1.6% of wrongly assigned residues. On the other hand, PatternLab is available to the community for free and may be used for all structural biology groups (62). Moreover, the use of a theoretical sequence database approximates it to a *de novo* sequencing and does not require the presence of remote homologue sequences in databases.

The use of the conservation estimation based on remote homologues (26) is an additional level of information, although cautious is necessary to prevent introduction of bias from what is known from the database of sequences. In fact, the models of BmooMP-I, MjTX-I and CBCol were not identical to a previously known sequence and this evidence the importance to have a proper strategy to model crystal heterogeneity. The MjTX-I had all four chains identical and with four residues difference to another protein from the venom of *B. moojeni* (BooMooTX). Differently, CBCol was in fact a heterodimer, having 8 residues different when chains are compared, and confirms the hypothesis that two isoforms could be distinguished using crystallographic data together with MS by SLIDER. The preference of an isoform in the dimer over the other is supported by the different crystal packing. The database of metalloproteinase sequences is smaller compared to these other enzymes and BmooMP-I was also a novel sequence.

Validation is an essential step in structure deposition and Molprobity (57) is the current software that performs a series of evaluation to check if model is physically plausible and is in agreement with experimental data. For protein structures obtained from natural source, SLIDER sub-routines could complement such analysis with sequence validation as for some residue positions single assignment may be given and for others it may be uncertain. Large and flexible residues with high B-factors may be confused with small amino acids in crystallography. Similar scenario is in single particle electron cryomicroscopy, as maps are electrostatic potential and negatively charged side chains are not observed (71).

In natural, non-recombinant species, sequence variability should be acknowledged rather than brushed aside. We propose a tool to address this intrinsic characteristic, by integration of different experimental and bioinformatic analysis, thus effectively combining different sources of information. Furthermore, the model deposited should reflect this resulting probability in microheterogeneities. Moreover, SLIDER approach to test multiple hypothesis of amino acids is not exclusive to toxins nor it is for crystallography. It can be applied for single particle electron cryomicroscopy maps, to assign nucleotides in DNA/RNA and to elucidate naturally purified proteins whose sequences cannot be retrieved from genomes. This was the case for feline haemoglobins, whose study was important to understand differences in erythrocyte life span from various organisms (44), and DING proteins (43), that are medically important due to ability inhibit carcinogenic cell growth (72, 73) and Transcription of HIV-1 LTR Promoter (74).

## AVAILABILITY

SEQUENCE SLIDER is an open-source initiative available in the GitHub repository (https://github.com/LBME/slider), in ARCIMBOLDO (http://chango.ibmb.csic.es/) through the package installer for Python (pip) and CCP4.

Mass spectrometry data of BmooMP-I (doi: 10.25345/C55J77), MjTX-I (doi: 10.25345/C59B95), CBCol (doi: 10.25345/C5X22B) and lysozyme (doi: 10.25345/C51V5B) are available at Mass Spectrometry Interactive Virtual Environment (MassIVE) https://massive.ucsd.edu/

## ACCESSION NUMBERS

Atomic coordinates and structure factors for the reported crystal structures have been deposited with the Protein Data bank under accession number 6X5X (BmooMP-I) and 7LYE (MjTX-I/varespladib).

## SUPPLEMENTARY DATA

Supplementary Tables are available at NAR online.

## ACKNOWLEDGEMENT

We thank the use of the Brazilian Synchrotron Light Laboratory (LNLS, Campinas, SP, Brazil) for the use of its facilities, the Institute of Biotechnology (IBTEC/UNESP – Botucatu, SP, Brazil) for the use of its Mass Spectrometry facilities and Paulo C. Carvalho for his aid processing the Mass Spectrometry data using his software, PatternLab.

## FUNDING

This work was financially supported by São Paulo Research Foundation (FAPESP, 2015/17286-0 and 2020/10143-7). RJB. was supported by FAPESP [16/24191-8] and IU by PGC [PGC2018-101370-BI00]. DCP and MRMF are CNPq fellow researchers [301974/2019-5 and 302883/2017-7].

## CONFLICT OF INTEREST

None declared.

## Notes

### Competing Interest Statement

The authors have declared no competing interest.

## REFERENCES

1. Wlodawer, A., Minor, W., Dauter, Z. and Jaskolski, M. (2008) Protein crystallography for non-crystallographers, or how to get the best (but not more) from published macromolecular structures: Protein crystallography for non-crystallographers. FEBS Journal, 275, 1–21.

2. Nakane, T., Kotecha, A., Sente, A., McMullan, G., Masiulis, S., Brown, P.M.G.E., Grigoras, I.T., Malinauskaite, L., Malinauskas, T., Miehling, J., et al. (2020) Single-particle cryo-EM at atomic resolution Biophysics.

3. Yip, K.M., Fischer, N., Paknia, E., Chari, A. and Stark, H. (2020) Breaking the next Cryo-EM resolution barrier – Atomic resolution determination of proteins! Biophysics.

4. Bartesaghi, A., Matthies, D., Banerjee, S., Merk, A. and Subramaniam, S. (2014) Structure of β – galactosidase at 3.2-Å resolution obtained by cryo-electron microscopy. PNAS, 111, 11709–11714.

5. Wang, J. and Moore, P.B. (2017) On the interpretation of electron microscopic maps of biological macromolecules: Electric Potentials Involving Negative Charges. Protein Science, 26, 122–129.

6. Ke, Z., Oton, J., Qu, K., Cortese, M., Zila, V., McKeane, L., Nakane, T., Zivanov, J., Neufeldt, C.J., Cerikan, B., et al. (2020) Structures and distributions of SARS-CoV-2 spike proteins on intact virions. Nature, 588, 498–502.

7. Radermacher, M., Rao, V., Grassucci, R., Frank, J., Timerman, A.P., Fleischer, S. and Wagenknecht, T. (1994) Cryo-electron microscopy and three-dimensional reconstruction of the calcium release channel/ryanodine receptor from skeletal muscle. Journal of Cell Biology, 127, 411–423.

8. Liu, Z., Zhang, J., Li, P., Chen, S.R.W. and Wagenknecht, T. (2002) Three-dimensional Reconstruction of the Recombinant Type 2 Ryanodine Receptor and Localization of Its Divergent Region 1. Journal of Biological Chemistry, 277, 46712–46719.

9. Yan, Z., Bai, X., Yan, C., Wu, J., Li, Z., Xie, T., Peng, W., Yin, C., Li, X., Scheres, S.H.W., et al. (2015) Structure of the rabbit ryanodine receptor RyR1 at near-atomic resolution. Nature, 517, 50–55.

10. Sharma, M.R., Penczek, P., Grassucci, R., Xin, H.-B., Fleischer, S. and Wagenknecht, T. (1998) Cryoelectron Microscopy and Image Analysis of the Cardiac Ryanodine Receptor *. Journal of Biological Chemistry, 273, 18429–18434.

11. Yan, Z., Zhou, Q., Wang, L., Wu, J., Zhao, Y., Huang, G., Peng, W., Shen, H., Lei, J. and Yan, N. (2017) Structure of the Nav1.4-β1 Complex from Electric Eel. Cell, 170, 470–482.e11.

12. Strynadka, N.C.J. and James, M.N.G. (1996) Lysozyme: A model enzyme in protein crystallography. In Jollès, P. (ed), Lysozymes: Model Enzymes in Biochemistry and Biology, Experientia Supplementum. Birkhäuser Basel, Basel, Vol. 75, pp. 185–222.

13. Moss, J.M., Van Damme, M.-P.I., Murphy, W.H., Stanton, P.G., Thomas, P. and Preston, B.N. (1997) Purification, Characterization, and Biosynthesis of Bovine Cartilage Lysozyme Isoforms. Archives of Biochemistry and Biophysics, 339, 172–182.

14. Zavalova, L.L., Artamonova, I.I., Berezhnoy, S.N., Tagaev, A.A., Baskova, I.P., Andersen, J., Roepstorff, P. and Egorov, Ts.A. (2003) Multiple forms of medicinal leech destabilase-lysozyme. Biochemical and Biophysical Research Communications, 306, 318–323.

15. Calvete, J.J. (2009) Venomics: digging into the evolution of venomous systems and learning to twist nature to fight pathology. J Proteomics, 72, 121–126.

16. Guércio, R.A., Shevchenko, A., Shevchenko, A., López-Lozano, J.L., Paba, J., Sousa, M.V. and Ricart, C.A. (2006) Ontogenetic variations in the venom proteome of the Amazonian snake Bothrops atrox. Proteome Sci, 4, 11.

17. Daltry, J.C., Wüster, W. and Thorpe, R.S. (1996) Diet and snake venom evolution. Nature, 379, 537–540.

18. Williams, V. and White, J. (1992) Variation in the composition of the venom from a single specimen of Pseudonaja textilis (common brown snake) over one year. Toxicon, 30, 202–206.

19. Massey, D.J., Calvete, J.J., Sánchez, E.E., Sanz, L., Richards, K., Curtis, R. and Boesen, K. (2012) Venom variability and envenoming severity outcomes of the Crotalus scutulatus scutulatus (Mojave rattlesnake) from Southern Arizona. Journal of Proteomics, 75, 2576–2587.

20. Menezes, M.C., Furtado, M.F., Travaglia-Cardoso, S.R., Camargo, A.C.M. and Serrano, S.M.T. (2006) Sex-based individual variation of snake venom proteome among eighteen Bothrops jararaca siblings. Toxicon, 47, 304–312.

21. Amorim, F., Costa, T., Baiwir, D., De Pauw, E., Quinton, L. and Sampaio, S. (2018) Proteopeptidomic, Functional and Immunoreactivity Characterization of Bothrops moojeni Snake Venom: Influence of Snake Gender on Venom Composition. Toxins, 10, 177.

22. Chu, C.-C., Li, S.-H. and Chen, Y.-H. (1995) Resolution of isotoxins in the β-bungarotoxin family. Journal of Chromatography A, 694, 492–497.

23. Faure, G., Guillaume, J.L., Camoin, L., Saliou, B. and Bon, C. (1991) Multiplicity of acidic subunit isoforms of crotoxin, the phospholipase A2 neurotoxin from Crotalus durissus terrificus venom, results from posttranslational modifications. Biochemistry, 30, 8074–8083.

24. Shih-Hsiung, W., Fu-Hsiung, C. and Mu-Chin, T. (1983) Separation of the subunits of crotoxin by high-performance liquid chromatography. Journal of Chromatography A, 259, 375–377.

25. Saul, F.A., Prijatelj-Znidarsic, P., Vulliez-le Normand, B., Villette, B., Raynal, B., Pungercar, J., Krizaj, I. and Faure, G. (2010) Comparative structural studies of two natural isoforms of ammodytoxin, phospholipases A2 from Vipera ammodytes ammodytes which differ in neurotoxicity and anticoagulant activity. J. Struct. Biol., 169, 360–369.

26. Ashkenazy, H., Abadi, S., Martz, E., Chay, O., Mayrose, I., Pupko, T. and Ben-Tal, N. (2016) ConSurf 2016: an improved methodology to estimate and visualize evolutionary conservation in macromolecules. Nucleic Acids Res, 44, W344–W350.

27. Landau, M., Mayrose, I., Rosenberg, Y., Glaser, F., Martz, E., Pupko, T. and Ben-Tal, N. (2005) ConSurf 2005: the projection of evolutionary conservation scores of residues on protein structures. Nucleic Acids Research, 33, W299–W302.

28. Marks, D.S., Hopf, T.A. and Sander, C. (2012) Protein structure prediction from sequence variation. Nat Biotechnol, 30, 1072–1080.

29. Figliuzzi, M., Barrat-Charlaix, P. and Weigt, M. (2018) How Pairwise Coevolutionary Models Capture the Collective Residue Variability in Proteins? Molecular Biology and Evolution, 35, 1018–1027.

30. Cocco, S., Feinauer, C., Figliuzzi, M., Monasson, R. and Weigt, M. (2018) Inverse statistical physics of protein sequences: a key issues review. Rep. Prog. Phys., 81, 032601.

31. de Juan, D., Pazos, F. and Valencia, A. (2013) Emerging methods in protein co-evolution. Nat Rev Genet, 14, 249–261.

32. Senior, A.W., Evans, R., Jumper, J., Kirkpatrick, J., Sifre, L., Green, T., Qin, C., Žídek, A., Nelson, A.W.R., Bridgland, A., et al. (2020) Improved protein structure prediction using potentials from deep learning. Nature, 577, 706–710.

33. Stiffler, M.A., Poelwijk, F.J., Brock, K.P., Stein, R.R., Riesselman, A., Teyra, J., Sidhu, S.S., Marks, D.S., Gauthier, N.P. and Sander, C. (2020) Protein Structure from Experimental Evolution. Cell Systems, 10, 15–24.e5.

34. Simkovic, F., Thomas, J.M.H., Keegan, R.M., Winn, M.D., Mayans, O. and Rigden, D.J. (2016) Residue contacts predicted by evolutionary covariance extend the application of ab initio molecular replacement to larger and more challenging protein folds. IUCrJ, 3, 259–270.

35. Saraswathy, N. and Ramalingam, P. (2011) Protein sequencing techniques. In Concepts and Techniques in Genomics and Proteomics. Elsevier, pp. 193–201.

36. Domon, B. (2006) Mass Spectrometry and Protein Analysis. Science, 312, 212–217.

37. Wilson, D. and Daly, N.L. (2018) Venomics: A Mini-Review. High Throughput, 7.

38. Calvete, J.J., Juárez, P. and Sanz, L. (2007) Snake venomics. Strategy and applications. J Mass Spectrom, 42, 1405–1414.

39. Liska, A.J. and Shevchenko, A. (2003) Expanding the organismal scope of proteomics: Cross-species protein identification by mass spectrometry and its implications. Proteomics, 3, 19–28.

40. Liska, A.J. and Shevchenko, A. (2003) Combining mass spectrometry with database interrogation strategies in proteomics. TrAC Trends in Analytical Chemistry, 22, 291–298.

41. Standing, K. (2003) Peptide and protein de novo sequencing by mass spectrometry. Current Opinion in Structural Biology, 13, 595–601.

42. Cohen, S.L. and Chait, B.T. (2001) Mass Spectrometry as a Tool for Protein Crystallography. Annu. Rev. Biophys. Biomol. Struct., 30, 67–85.

43. Diemer, H., Elias, M., Renault, F., Rochu, D., Contreras-Martel, C., Schaeffer, C., Dorsselaer, A.V. and Chabriere, E. (2008) Tandem use of X-ray crystallography and mass spectrometry to obtain ab initio the complete and exact amino acids sequence of HPBP, a human 38-kDa apolipoprotein. Proteins: Structure, Function, and Bioinformatics, 71, 1708–1720.

44. Guo, J., Uppal, S., Easthon, L.M., Mueser, T.C. and Griffith, W.P. (2012) Complete sequence determination of hemoglobin from endangered feline species using a combined ESI-MS and X-ray crystallography approach. International Journal of Mass Spectrometry, 312, 70–77.

45. Chojnowski, G., Simpkin, A.J., Leonardo, D.A., Seifert-Davila, W., Vivas-Ruiz, D.E., Keegan, R.M. and Rigden, D.J. (2021) Identification of unknown proteins in X-ray crystallography and cryo-EM Molecular Biology.

46. Borges, R.J., Meindl, K., Triviño, J., Sammito, M., Medina, A., Millán, C., Alcorlo, M., Hermoso, J.A., Fontes, M.R. de M. and Usón, I. (2020) *SEQUENCE SLIDER*: expanding polyalanine fragments for phasing with multiple side-chain hypotheses. Acta Crystallogr D Struct Biol, 76.

47. Sammito, M., Millán, C., Frieske, D., Rodríguez-Freire, E., Borges, R.J. and Usón, I. (2015) ARCIMBOLDO_LITE: single-workstation implementation and use. Acta Crystallogr. D Biol. Crystallogr., 71, 1921–1930.

48. McCoy, A.J., Grosse-Kunstleve, R.W., Adams, P.D., Winn, M.D., Storoni, L.C. and Read, R.J. (2007) Phaser crystallographic software. Journal of Applied Crystallography, 40, 658–674.

49. Thorn, A. and Sheldrick, G.M. (2013) Extending molecular-replacement solutions with SHELXE. Acta Crystallogr. D Biol. Crystallogr., 69, 2251–2256.

50. Salvador, G.H.M., Borges, R.J., Eulálio, M.M.C., dos Santos, L.D. and Fontes, M.R.M. (2020) Biochemical, pharmacological and structural characterization of BmooMP-I, a new P–I metalloproteinase from Bothrops moojeni venom. Biochimie, 179, 54–64.

51. Hendon, R.A. and Fraenkel-Conrat, H. (1971) Biological Roles of the Two Components of Crotoxin. Proceedings of the National Academy of Sciences, 68, 1560–1563.

52. Salvador, G.H.M., Fernandes, C.A.H., Corrêa, L.C., Santos-Filho, N.A., Soares, A.M. and Fontes, M.R.M. (2009) Crystallization and preliminary X-ray diffraction analysis of crotoxin B from *Crotalus durissus collilineatus* venom. Acta Crystallogr F Struct Biol Cryst Commun, 65, 1011–1013.

53. Otwinowski, Z. and Minor, W. (1997) Processing of X-ray diffraction data collected in oscillation mode. In. Elsevier, Vol. 276, pp. 307–326.

54. Kabsch, W. (2010) XDS. Acta Crystallogr. D Biol. Crystallogr., 66, 125–132.

55. Afonine, P.V., Grosse-Kunstleve, R.W., Echols, N., Headd, J.J., Moriarty, N.W., Mustyakimov, M., Terwilliger, T.C., Urzhumtsev, A., Zwart, P.H. and Adams, P.D. (2012) Towards automated crystallographic structure refinement with phenix.refine. Acta Crystallogr. D Biol. Crystallogr., 68, 352–367.

56. Emsley, P., Lohkamp, B., Scott, W.G. and Cowtan, K. (2010) Features and development of Coot. Acta Crystallographica Section D Biological Crystallography, 66, 486–501.

57. Davis, I.W., Leaver-Fay, A., Chen, V.B., Block, J.N., Kapral, G.J., Wang, X., Murray, L.W., Arendall, W.B., 3rd, Snoeyink, J., Richardson, J.S., et al. (2007) MolProbity: all-atom contacts and structure validation for proteins and nucleic acids. Nucleic Acids Res., 35, W375–383.

58. Emsley, P. and Cowtan, K. (2004) *Coot*: model-building tools for molecular graphics. Acta Crystallogr D Biol Crystallogr, 60, 2126–2132.

59. Liebschner, D., Afonine, P.V., Moriarty, N.W., Poon, B.K., Sobolev, O.V., Terwilliger, T.C. and Adams, P.D. (2017) Polder maps: improving OMIT maps by excluding bulk solvent. Acta Crystallographica Section D Structural Biology, 73, 148–157.

60. Merritt, E.A. and Bacon, D.J. (1997) [26] Raster3D: Photorealistic molecular graphics. In Methods in Enzymology. Elsevier, Vol. 277, pp. 505–524.

61. The UniProt Consortium (2019) UniProt: a worldwide hub of protein knowledge. Nucleic Acids Research, 47, D506–D515.

62. Carvalho, P.C., Lima, D.B., Leprevost, F.V., Santos, M.D.M., Fischer, J.S.G., Aquino, P.F., Moresco, J.J., Yates, J.R. and Barbosa, V.C. (2016) Integrated analysis of shotgun proteomic data with PatternLab for proteomics 4.0. Nat Protoc, 11, 102–117.

63. Han, X., He, L., Xin, L., Shan, B. and Ma, B. (2011) PeaksPTM: Mass Spectrometry-Based Identification of Peptides with Unspecified Modifications. J. Proteome Res., 10, 2930–2936.

64. Han, Y., Ma, B. and Zhang, K. (2005) SPIDER: SOFTWARE FOR PROTEIN IDENTIFICATION FROM SEQUENCE TAGS WITH *DE NOVO* SEQUENCING ERROR. J. Bioinform. Comput. Biol., 03, 697–716.

65. Winn, M.D., Ballard, C.C., Cowtan, K.D., Dodson, E.J., Emsley, P., Evans, P.R., Keegan, R.M., Krissinel, E.B., Leslie, A.G.W., McCoy, A., et al. (2011) Overview of the CCP4 suite and current developments. Acta Crystallographica Section D Biological Crystallography, 67, 235–242.

66. NCBI Resource Coordinators, Agarwala, R., Barrett, T., Beck, J., Benson, D.A., Bollin, C., Bolton, E., Bourexis, D., Brister, J.R., Bryant, S.H., et al. (2018) Database resources of the National Center for Biotechnology Information. Nucleic Acids Research, 46, D8–D13.

67. Colvin, J.R., Smith, D.B. and Cook, W.H. (1954) The Microheterogeneity of Proteins. Chem. Rev., 54, 687–711.

68. Salvador, G.H.M., Fernandes, C.A.H., Magro, A.J., Marchi-Salvador, D.P., Cavalcante, W.L.G., Fernandez, R.M., Gallacci, M., Soares, A.M., Oliveira, C.L.P. and Fontes, M.R.M. (2013) Structural and phylogenetic studies with MjTX-I reveal a multi-oligomeric toxin--a novel feature in Lys49-PLA2s protein class. PLoS ONE, 8, e60610.

69. Salvador, G.H.M., Borges, R.J., Lomonte, B., Lewin, M.R. and Fontes, M.R.M. (2021) The synthetic varespladib molecule is a multi-functional inhibitor for PLA2 and PLA2-like ophidic toxins. Biochimica et Biophysica Acta (BBA) – General Subjects, 1865, 129913.

70. Sampaio, S.C., Hyslop, S., Fontes, M.R.M., Prado-Franceschi, J., Zambelli, V.O., Magro, A.J., Brigatte, P., Gutierrez, V.P. and Cury, Y. (2010) Crotoxin: Novel activities for a classic β-neurotoxin. Toxicon, 55, 1045–1060.

71. Marques, M.A., Purdy, M.D. and Yeager, M. (2019) CryoEM maps are full of potential. Current Opinion in Structural Biology, 58, 214–223.

72. Adams, L., Davey, S. and Scott, K. (2002) The DING protein: an autocrine growth-stimulatory protein related to the human synovial stimulatory protein. Biochimica et Biophysica Acta (BBA) – Molecular Basis of Disease, 1586, 254–264.

73. Bookland, M.J., Darbinian, N., Weaver, M., Amini, S. and Khalili, K. (2012) Growth inhibition of malignant glioblastoma by DING protein. J Neurooncol, 107, 247–256.

74. Sachdeva, R., Darbinian, N., Khalili, K., Amini, S., Gonzalez, D., Djeghader, A., Chabriére, E., Suh, A., Scott, K. and Simm, M. (2013) DING Proteins from Phylogenetically Different Species Share High Degrees of Sequence and Structure Homology and Block Transcription of HIV-1 LTR Promoter. PLoS ONE, 8, e69623.

