## Supplementary Table for "SEQUENCE SLIDER: integration of structural and genetic data to characterize isoforms from natural source"

SUPPLEMENTARY DATA

PrimaryScore from PatternLab is a metric to compare experimental and theoretical spectrum.

Supplementary Table 1: Residues of lysozyme resolved by crystallography and MS and/or phylogenetic analysis

| | | Crystallography | | | MS | ConSurf | |
|---|---|---|---|---|---|---|---|
| Res# | ResType | RSCC | ΔCont | RSCC* | PrimaryScore | n | % |
| 1 | K | 92.2 | 9.3 | 94 | | 173 | 95.6 |
| 3 | F | 94 | 4 | 92.2 | | 93 | 36.2 |
| 4 | G | 90.9 | 24.3 | 94.7 | | 37 | 13.7 |
| 5 | R | 92.9 | 3.3 | 91.5 | | 211 | 76.7 |
| 6 | C | 94.5 | 21.7 | 95.2 | 1.73 | 275 | 98.9 |
| 9 | A | 90.3 | 5.7 | 93.4 | 1.73 | 242 | 86.7 |
| 10 | A | 92.3 | 12.4 | 92.6 | 1.73 | 5 | 1.8 |
| 11 | A | 85.2 | 3.2 | 93 | 1.73 | 24 | 8.6 |
| 12 | M | 95.9 | 3.3 | 92 | 1.73 | 22 | 7.9 |
| 13 | K | 89.2 | 5.8 | 88.4 | 1.73 | 87 | 31.3 |
| 14 | R | 89.3 | 4.2 | 88.8 | 1.12 | 74 | 26.4 |
| 16 | G | 77.6 | 20.1 | 91.9 | 2.02 | 190 | 69.6 |
| 20 | Y | 93.7 | 7.7 | 94.3 | 2.02 | 91 | 55.8 |
| 22 | G | 83 | 11.5 | 91.2 | 2.71 | 152 | 54.1 |
| 23 | Y | 95.3 | 7 | 94.2 | 2.71 | 79 | 27.9 |
| 24 | S | 92.8 | 7.4 | 94 | 2.71 | 133 | 46.8 |
| 26 | G | 94 | 9.7 | 93.7 | 2.71 | 37 | 12.9 |
| 28 | W | 95.1 | 7.2 | 93 | 2.71 | 273 | 94.8 |
| 30 | C | 97.1 | 12 | 94.1 | 2.71 | 295 | 99.7 |
| 31 | A | 90.1 | 11.2 | 94.3 | 2.71 | 1 | 0.3 |
| 32 | A | 93 | 9.1 | 95.4 | 2.71 | 201 | 67.9 |
| 33 | K | 91.2 | 7.4 | 95.1 | 2.71 | 43 | 14.5 |
| 38 | F | 95 | 5.4 | 94.3 | 2.44 | 142 | 48.0 |
| 42 | A | 94.5 | 8.4 | 96.8 | 2.44 | 206 | 69.4 |
| 43 | T | 95.1 | 5.2 | 95.5 | 2.44 | 52 | 17.5 |
| 45 | R | 92.4 | 4.9 | 93.9 | 2.65 | 53 | 18.0 |
| 49 | G | 80.2 | 29 | 90.5 | 3.27 | 213 | 72.2 |
| 50 | S | 94.4 | 8.1 | 94.1 | 3.27 | 252 | 85.1 |
| 53 | Y | 95.5 | 5.5 | 93.4 | 3.27 | 198 | 66.7 |
| 54 | G | 92 | 15.6 | 94.9 | 3.27 | 294 | 98.7 |
| 58 | I | 85.5 | 3.7 | 91.6 | 3.27 | 279 | 93.6 |
| 60 | S | 94.1 | 5.7 | 92.9 | 3.27 | 162 | 54.5 |
| 62 | W | 92 | 6.4 | 90.3 | 1.35 | 48 | 16.3 |
| 63 | W | 93 | 8.7 | 93.3 | 1.35 | 292 | 99.0 |
| 64 | C | 97.2 | 12.7 | 92.6 | 1.35 | 297 | 99.7 |
| 67 | G | 83 | 14.1 | 94.4 | 1.35 | 130 | 44.5 |
| 68 | R | 94.1 | 9.4 | 90.9 | 1.35 | 39 | 13.3 |
| 71 | G | 67.3 | 16.2 | 85.8 | 2.15 | 27 | 19.3 |
| 72 | S | 79.9 | 3.3 | 79 | 2.15 | 87 | 31.0 |
| 73 | R | 76.3 | **2** | 67.8 | 2.15 | 13 | 4.6 |
| 76 | C | 95.4 | 13.4 | 91.8 | 3.85 | 296 | 99.3 |
| 79 | P | 91.4 | 12.9 | 91.9 | 3.85 | 45 | 15.1 |
| 80 | C | 96.9 | 13.8 | 94.3 | 3.85 | 296 | 99.3 |

| | | | | | | | |
|---|---|---|---|---|---|---|---|
| 82 | A | 90.5 | 12.2 | 94.3 | 3.85 | 64 | 22.1 |
| 84 | L | 91.3 | 5.3 | 94.1 | 3.85 | 199 | 70.6 |
| 89 | T | 90 | 4.2 | 93.5 | 3.85 | 100 | 36.0 |
| 90 | A | 87.1 | 13.7 | 95.9 | 3.85 | 24 | 8.6 |
| 94 | C | 96.3 | 16.5 | 91.4 | 3.85 | 278 | 99.6 |
| 95 | A | 90.4 | 10.1 | 94.7 | 3.85 | 216 | 77.7 |
| 96 | K | 91.7 | 5 | 93.5 | 3.85 | 196 | 70.5 |
| 98 | I | 94.2 | 7.5 | 94.8 | 3.39 | 201 | 72.3 |
| 100 | S | 91.1 | 7.8 | 90.3 | 3.39 | 42 | 15.2 |
| 102 | G | 78.2 | 18.9 | 89.8 | 3.39 | 1 | 0.4 |
| 104 | G | 85.4 | 9.2 | 94.1 | 3.39 | 259 | 93.8 |
| 105 | M | 95.8 | 3.3 | 94.6 | 3.39 | 70 | 25.4 |
| 107 | A | 89.2 | 16.1 | 91.8 | 3.39 | 199 | 72.1 |
| 108 | W | 94.7 | 12.6 | 91.7 | 3.39 | 273 | 99.3 |
| 110 | A | 93.8 | 11.7 | 93.8 | 3.39 | 95 | 35.1 |
| 111 | W | 95.9 | 12.7 | 91.9 | 3.39 | 205 | 74.5 |
| 114 | R | 91.4 | 4.6 | 90.2 | 1.07 | 22 | 8.1 |
| 115 | C | 95.9 | 10.6 | 91.3 | 1.07 | 265 | 98.9 |
| 116 | K | 94.5 | 11 | 95.2 | 1.07 | 62 | 23.4 |
| 118 | T | 95.5 | 5.1 | 93.8 | 1.67 | 12 | 5.3 |
| 120 | V | 94.6 | 3.6 | 93.3 | 1.67 | 18 | 11.4 |
| 122 | A | 86.4 | 8.2 | 89.4 | 1.67 | 5 | 3.0 |
| 123 | W | 93.4 | 13.8 | 91.2 | 1.67 | 79 | 47.3 |
| 126 | G | 78.7 | 26.1 | 88.4 | 1.67 | 130 | 81.2 |
| 127 | C | 94 | 12.7 | 92.5 | 1.20 | 159 | 99.4 |

Res# stands for residue number; ResType for residues type; RSCC for real-space correlation coefficient; ΔCont for minimum difference within other RSCC; * for main chain atoms; MS for mass spectrometry. RSCC in bold denotes cysteine partially occupied having ΔCont less than 3.0.

Supplementary Table 2: Residues of lysozyme having assignment ambiguity from crystallographic data being resolved by mass spectrometry

| | | Crystallography | | | MS | ConSurf | |
|---|---|---|---|---|---|---|---|
| Res# | ResType | RSCC | ΔCont | Observation | PrimaryScore | n | % |
| 7 | E! | 93.5 | 0.6 | | 1.73 | 167 | 59.9 |
| 7 | Q | 93 | 5.8 | | | 12 | 4.3 |
| 7 | MainCh | 94.6 | | | | | |
| 8 | L! | 92.5 | 1 | Hydrophobic | 1.73 | 215 | 77.1 |
| 8 | N | 91.5 | 0.5 | | | 0 | 0.0 |
| 8 | D | 91 | 7.5 | | | 0 | 0.0 |
| 8 | MainCh | 93.7 | | | | | |
| 15 | H! | 93.5 | 2.1 | | 2.02 | 60 | 22.1 |
| 15 | F | 91.4 | 0.5 | | | 20 | 7.4 |
| 15 | K | 90.9 | 2.1 | | | 6 | 2.2 |
| 15 | MainCh | 93.4 | | | | | |
| 17 | D | 91.3 | 0.2 | | | 0 | 0.0 |
| 17 | N | 91.1 | 2 | | | 0 | 0.0 |

| | | | | | | | |
|---|---|---|---|---|---|---|---|
| 17 | L! | 89.1 | 8.1 | Hydrophobic | 2.02 | 110 | 39.3 |
| 17 | MainCh | 88.4 | | | | | |
| 18 | N | 91.3 | 0.7 | 2 H-bonds | | 12 | 7.1 |
| 18 | D! | 90.6 | 1 | | 2.02 | 120 | 71.4 |
| 18 | L | 89.6 | 7.6 | | | 0 | 0.0 |
| 18 | MainCh | 93.5 | | | | | |
| 25 | L! | 91.5 | 1.9 | Hydrophobic | 2.71 | 198 | 69.2 |
| 25 | N | 89.6 | 0.6 | | | 0 | 0.0 |
| 25 | D | 89 | 7.1 | | | 0 | 0.0 |
| 25 | MainCh | 92.3 | | | | | |
| 29 | V! | 95.5 | 1.1 | Hydrophobic | 2.78 | 160 | 54.1 |
| 29 | T | 94.5 | 4.1 | | | 18 | 6.1 |
| 29 | MainCh | 91.4 | | | | | |
| 34 | F! | 95.7 | 2.3 | | 2.44 | 39 | 13.2 |
| 34 | H | 93.5 | 3.6 | | | 97 | 32.8 |
| 34 | MainCh | 94.8 | | | | | |
| 35 | E! | 94.2 | 1.2 | | 2.44 | 248 | 83.8 |
| 35 | Q | 93 | 4.2 | | | 3 | 1.0 |
| 35 | MainCh | 94.7 | | | | | |
| 36 | S! | 92.7 | 1.2 | | 2.44 | 290 | 98.0 |
| 36 | C | 91.5 | 10.1 | | | 0 | 0.0 |
| 36 | MainCh | 93.5 | | | | | |
| 39 | N! | 94.5 | 1.5 | 4 H-bonds | 2.44 | 208 | 70.3 |
| 39 | D | 93 | 1.2 | | | 48 | 16.2 |
| 39 | L | 91.8 | 7.1 | | | 0 | 0.0 |
| 39 | MainCh | 94.5 | | | | | |
| 40 | T! | 91.1 | 0.7 | 3 H-bonds | 2.44 | 249 | 84.1 |
| 40 | V | 90.4 | 3.6 | | | 2 | 0.7 |
| 40 | MainCh | 95.1 | | | | | |
| 44 | N! | 92.9 | 2.8 | 4 H-bonds | 2.44 | 106 | 35.9 |
| 44 | L | 90.1 | 0.1 | | | 0 | 0.0 |
| 44 | D | 90 | 6.8 | | | 20 | 6.8 |
| 44 | MainCh | 96 | | | | | |
| 47 | V | 86.2 | 4.6 | 3 H-bonds | | 10 | 3.6 |
| 47 | T! | 81.7 | 1.2 | | 3.27 | 31 | 11.0 |
| 47 | MainCh | 85.9 | | | | | |
| 51 | T! | 95.5 | 1.5 | 2 H-bonds | 3.27 | 118 | 39.9 |
| 51 | V | 94 | 5.4 | | | 8 | 2.7 |
| 51 | MainCh | 94.3 | | | | | |

| | | | | | | | |
|---|---|---|---|---|---|---|---|
| 55 | V | 90.4 | 1.3 | Hydrophobic | | 2 | 0.7 |
| 55 | T | 89.1 | 4.8 | | | 0 | 0.0 |
| 55 | S | 84.3 | 0 | | | 0 | 0.0 |
| 55 | I! | 84.3 | 4.5 | | 3.28 | 179 | 60.1 |
| 55 | MainCh | 93.2 | | | | | |
| 56 | N | 91 | 0.8 | Hydrophobic | | 0 | 0.0 |
| 56 | D | 90.2 | 2.3 | | | 0 | 0.0 |
| 56 | L! | 88 | 4.3 | | 3.28 | 10 | 3.4 |
| 56 | MainCh | 91.6 | | | | | |
| 61 | S | 84.6 | 1.6 | | | 6 | 2.0 |
| 61 | R! | 83 | 2.8 | | 3.28 | 111 | 37.2 |
| 61 | MainCh | 87.7 | | | | | |
| 65 | D | 95.1 | 0.8 | 2 H-bonds | | 41 | 13.8 |
| 65 | N! | 94.3 | 1.9 | | 1.35 | 75 | 25.3 |
| 65 | L | 92.5 | 2.8 | | | 0 | 0.0 |
| 65 | MainCh | 94.8 | | | | | |
| 69 | S | 93.2 | 0.2 | 4 H-bonds | | 27 | 13.9 |
| 69 | T! | 93 | 2 | | 2.15 | 66 | 34.0 |
| 69 | C | 91 | 7.2 | | | 0 | 0.0 |
| 69 | MainCh | 93.8 | | | | | |
| 74 | D | 93.9 | 2.9 | 3 H-bonds | 2.69 | 11 | 3.7 |
| 74 | N! | 90.9 | 0.8 | | 3.85 | 202 | 67.8 |
| 74 | MainCh | 74.5 | | | | | |
| 75 | L! | 92.1 | 0.5 | Exposed | 3.85 | 40 | 13.4 |
| 75 | S | 91.6 | 3.8 | | 2.15 | 2 | 0.7 |
| 75 | MainCh | 90.6 | | | | | |
| 77 | A | 82.5 | 3 | | 2.15 | 1 | 0.3 |
| 77 | V | 79.5 | 0.4 | | | 1 | 0.3 |
| 77 | S | 79.1 | 1.1 | | 2.00 | 8 | 2.7 |
| 77 | I | 78 | 1 | | | 0 | 0.0 |
| 77 | L | 76.9 | 1.1 | | | 0 | 0.0 |
| 77 | N! | 75.8 | 0.3 | | 3.85 | 75 | 25.2 |
| 77 | MainCh | 93.3 | | | | | |
| 78 | T | 90.3 | 1.2 | Hydrophobic | | 12 | 4.0 |
| 78 | V | 89.1 | 0.2 | | 2.00 | 82 | 27.5 |
| 78 | I! | 88.9 | 6.4 | | 3.85 | 116 | 38.9 |
| 78 | MainCh | 91.5 | | | | | |
| 81 | S! | 91.3 | 2.5 | | 3.85 | 117 | 39.3 |
| 81 | T | 88.8 | 2.7 | | 2.15 | 15 | 5.0 |
| 81 | MainCh | 92.8 | | | | | |

| | | | | | | | |
|---|---|---|---|---|---|---|---|
| 83 | L! | 90.5 | 2.8 | Hydrophobic | 3.85 | 222 | 77.9 |
| 83 | N | 87.7 | 1.3 | | | 0 | 0.0 |
| 83 | D | 86.4 | 5.5 | | | 0 | 0.0 |
| 83 | MainCh | 93.5 | | | | | |
| 88 | I! | 92.2 | 2.6 | | 3.85 | 181 | 65.1 |
| 88 | V | 89.6 | 0.8 | | | 11 | 4.0 |
| 88 | MainCh | 95.5 | | | | | |
| 91 | S! | 87.6 | 1.3 | 2 H-bonds | 3.85 | 41 | 14.7 |
| 91 | T | 86.3 | 2.7 | | 1.87 | 27 | 9.7 |
| 91 | MainCh | 91.8 | | | | | |
| 92 | I | 87.9 | 0.4 | | | 126 | 45.2 |
| 92 | T | 87.5 | 0.1 | | | 4 | 1.4 |
| 92 | V! | 87.4 | 7.3 | | 3.85 | 112 | 40.1 |
| 92 | MainCh | 92.3 | | | | | |
| 99 | V! | 92.3 | 0.6 | Hydrophobic | 3.39 | 118 | 42.6 |
| 99 | T | 91.7 | 5.2 | | | 3 | 1.1 |
| 99 | MainCh | 95.4 | | | | | |
| 106 | D | 93.9 | 0.5 | 3 H-bonds | 1.97 | 21 | 7.6 |
| 106 | N! | 93.4 | 1.5 | | 3.39 | 56 | 20.3 |
| 106 | L | 91.9 | 0.8 | | | 1 | 0.4 |
| 106 | E | 91.1 | 9.1 | | | 7 | 2.5 |
| 106 | MainCh | 93.5 | | | | | |
| 109 | T | 85.8 | 0.8 | Exposed | 1.15 | 36 | 13.3 |
| 109 | V! | 85 | 0.9 | | 3.39 | 73 | 26.9 |
| 109 | S | 84.1 | 3.6 | | | 19 | 7.0 |
| 109 | MainCh | 92.2 | | | | | |
| 112 | R! | 86.9 | 0.9 | | 3.39 | 68 | 25.5 |
| 112 | K | 86.1 | 1 | | | 82 | 30.7 |
| 112 | S | 85 | 2.4 | | | 14 | 5.2 |
| 112 | MainCh | 91.7 | | | | | |
| 117 | G! | 64.2 | 0.3 | | 1.67 | 139 | 54.5 |
| 117 | K | 63.9 | 8.6 | | | 3 | 1.2 |
| 117 | MainCh | 89.6 | | | | | |
| 124 | T | 92.6 | 2.1 | Exposed | | 18 | 11.0 |
| 124 | V | 90.5 | 2.1 | | 1.34 | 56 | 34.4 |
| 124 | I! | 88.4 | 4.7 | | 1.67 | 32 | 19.6 |
| 124 | MainCh | 93.7 | | | | | |
| 125 | A | 83.4 | 4.3 | | | 20 | 12.5 |
| 125 | G | 79.2 | 2.2 | | 0.82 | 1 | 0.6 |

| | | | | | | | |
|---|---|---|---|---|---|---|---|
| 125 | S | 76.9 | 4.8 | | 0.90 | 9 | 5.6 |
| 125 | K | 72.2 | 1.7 | | 1.34 | 27 | 16.9 |
| 125 | R! | 70.5 | 0.5 | | 1.67 | 25 | 15.6 |
| 125 | MainCh | 91.4 | | | | | |
| 128 | S | 88.2 | 2.6 | | | 0 | 0.0 |
| 128 | T | 85.6 | 1.4 | | | 2 | 1.6 |
| 128 | V | 84.2 | 1.3 | | 0.82 | 0 | 0.0 |
| 128 | A | 82.8 | 5 | | | 0 | 0.0 |
| 128 | D | 77.8 | 2.7 | | 0.90 | 20 | 16.1 |
| 128 | P | 75.1 | 0.5 | | 1.07 | 0 | 0.0 |
| 128 | L | 74.6 | 2.5 | | | 0 | 0.0 |
| 128 | I | 72.1 | 2.1 | | | 0 | 0.0 |
| 128 | N | 70 | 1.1 | | 1.64 | 15 | 12.1 |
| 128 | C | 68.9 | last | | | 0 | 0.0 |
| 128 | R! | | | | 1.20 | 13 | 10.5 |
| 128 | MainCh | 79.4 | | | | | |
| 129 | N | 81.1 | 0.4 | Hydrophobic | | 0 | 0.0 |
| 129 | D | 80.7 | 0.4 | | | 0 | 0.0 |
| 129 | L! | 80.2 | 2.8 | | 1.07 | 70 | 97.2 |
| 129 | MainCh | 71.1 | | | | | |

Res# stands for residue number; ResType for residues type; RSCC for real-space correlation coefficient; ΔCont for minimum difference within other RSCC; MainCh for main chain atoms; MS for mass spectrometry; H-bond for hydrogen bond; ! for resolved residue.

Supplementary Table 3: Residues of lysozyme having ambiguity assignment from both crystallographic and mass spectrometry data but being resolved by phylogenetic analysis

| | | Crystallography | | | MS | ConSurf | |
|---|---|---|---|---|---|---|---|
| Res# | ResType | RSCC | ΔCont | Observation | PrimaryScore | n | % |
| 2 | V! | 92.5 | 2.3 | Exposed | | 104 | 47.5 |
| 2 | T | 90.2 | 5.5 | | | 20 | 9.1 |
| 2 | MainCh | 94.6 | | | | | |
| 19 | S | 85 | 3.6 | | | 1 | 0.6 |
| 19 | A | 81.4 | 2.4 | | | 0 | 0.0 |
| 19 | V | 79 | 1.4 | | | 0 | 0.0 |
| 19 | L | 77.6 | 0.7 | | | 1 | 0.6 |
| 19 | I | 76.9 | 0.3 | | | 0 | 0.0 |
| 19 | K | 76.6 | 1.7 | | | 4 | 2.5 |
| 19 | G | 74.9 | 0.1 | | | 124 | 76.5 |
| 19 | T | 74.8 | 1.3 | | | 1 | 0.6 |
| 19 | N! | 73.5 | 2.1 | | 1.88 | 14 | 8.6 |
| 19 | D | 71.4 | 1 | | 2.02 | 2 | 1.2 |
| 19 | MainCh | 91.8 | | | | | |
| 21 | K | 83.1 | 1.7 | | 2.40 | 17 | 6.1 |
| 21 | A | 81.4 | 0.5 | | | 8 | 2.9 |
| 21 | R! | 80.9 | 3.1 | | 2.71 | 58 | 20.8 |
| 21 | MainCh | 95.7 | | | | | |

| | | | | | | | |
|---|---|---|---|---|---|---|---|
| 27 | N! | 95.6 | 0.6 | 4 H-bonds | 2.78 | 122 | 42.4 |
| 27 | D | 95 | 3.4 | | 2.55 | 72 | 25.0 |
| 27 | MainCh | 93.8 | | | | | |
| 37 | N! | 93.1 | 0.6 | 1 H-bonds | 2.44 | 55 | 18.6 |
| 37 | D | 92.5 | 1.3 | | 2.13 | 14 | 4.7 |
| 37 | L | 91.2 | 4.4 | | | 2 | 0.7 |
| 37 | MainCh | 94.7 | | | | | |
| 41 | E | 95.4 | 1.3 | | 1.95 | 5 | 1.7 |
| 41 | Q! | 94.1 | 3.2 | | 2.44 | 35 | 11.8 |
| 41 | MainCh | 93.5 | | | | | |
| 46 | D | 94.6 | 1.3 | 3 H-bonds | 3.27 | 0 | 0.0 |
| 46 | N! | 93.3 | 2.5 | | 3.11 | 106 | 41.9 |
| 46 | MainCh | 88.8 | | | | | |
| 48 | D! | 96.2 | 0.3 | 5 H-bonds | 3.27 | 178 | 62.0 |
| 48 | N | 96 | 9.6 | | 2.63 | 53 | 18.5 |
| 48 | MainCh | 93.8 | | | | | |
| 52 | D! | 93.8 | 0.9 | 5 H-bonds | 3.28 | 212 | 71.1 |
| 52 | N | 92.9 | 6.1 | | 3.01 | 32 | 10.7 |
| 52 | MainCh | 96.6 | | | | | |
| 57 | E | 91.1 | 0.2 | | 1.44 | 1 | 0.3 |
| 57 | Q! | 90.9 | 5 | | 1.32 | 293 | 98.3 |
| 57 | MainCh | 93.9 | | | | | |
| 59 | N! | 85.1 | 0.1 | 4 H-bonds | 3.28 | 207 | 69.5 |
| 59 | M | 84.9 | 2.3 | | | 0 | 0.0 |
| 59 | D | 82.6 | 0.4 | | 3.12 | 2 | 0.7 |
| 59 | MainCh | 90.7 | | | | | |
| 66 | D! | 95.6 | 1.4 | 8 H-bonds | 1.00 | 110 | 37.2 |
| 66 | N | 94.3 | 8.2 | | 0.60 | 45 | 15.2 |
| 66 | MainCh | 96.2 | | | | | |
| 85 | T! | 89.6 | 2 | | 2.98 | 45 | 16.4 |
| 85 | S | 87.6 | 3 | | 3.85 | 35 | 12.7 |
| 85 | MainCh | 91.1 | | | | | |
| 86 | A | 84.1 | 3.7 | 2 H-bonds | 2.00 | 6 | 2.2 |
| 86 | V | 80.4 | 0.6 | | | 1 | 0.4 |
| 86 | S! | 79.8 | 3.9 | | 3.85 | 30 | 10.8 |
| 86 | T | 75.8 | 1.4 | | 1.88 | 2 | 0.7 |
| 86 | MainCh | 91.3 | | | | | |
| 87 | D! | 91.7 | 0.1 | 2 H-bonds | 3.85 | 179 | 64.4 |

| | | | | | | | |
|---|---|---|---|---|---|---|---|
| 87 | N | 91.6 | 1.4 | | 2.94 | 76 | 27.3 |
| 87 | Q | 90.3 | 4.4 | | | 0 | 0.0 |
| 87 | MainCh | 94.2 | | | | | |
| 93 | N! | 91.8 | 2.9 | 1 H-bonds | 3.85 | 24 | 8.6 |
| 93 | D | 88.9 | 1 | | 3.56 | 4 | 1.4 |
| 93 | MainCh | 93 | | | | | |
| 97 | A | 83.4 | 2.8 | | | 1 | 0.4 |
| 97 | K! | 80.7 | 0.2 | | 2.92 | 107 | 38.5 |
| 97 | MainCh | 94.1 | | | | | |
| 101 | N | 92.9 | 1 | 1 H-bonds | 3.39 | 4 | 2.9 |
| 101 | D! | 91.9 | 1 | | 3.09 | 60 | 43.8 |
| 101 | L | 90.9 | 6.8 | | | 0 | 0.0 |
| 101 | MainCh | 91.2 | | | | | |
| 103 | N | 88 | 1.3 | 1 H-bonds | 2.91 | 24 | 8.7 |
| 103 | L | 86.7 | 0.9 | | | 2 | 0.7 |
| 103 | D! | 85.9 | 6.6 | | 3.39 | 9 | 3.3 |
| 103 | MainCh | 94.4 | | | | | |
| 113 | N! | 96.2 | 2.4 | 2 H-bonds | 1.07 | 61 | 24.0 |
| 113 | D | 93.8 | 1.2 | | | 7 | 2.8 |
| 113 | MainCh | 91.3 | | | | | |
| 119 | D! | 91.5 | 1.6 | 2 H-bonds | 1.67 | 102 | 50.7 |
| 119 | N | 90 | 0.9 | | 1.07 | 29 | 14.4 |
| 119 | L | 89.1 | 7 | | | 0 | 0.0 |
| 119 | MainCh | 92.3 | | | | | |

Res# stands for residue number; ResType for residues type; RSCC for real-space correlation coefficient; ΔCont for contrast difference within other RSCC; MainCh for main chain atoms; MS for mass spectrometry; H-bond for hydrogen bond; ! for resolved residue.

Supplementary Table 4: Residues of lysozyme having ambiguity assignment from both crystallographic and mass spectrometry data but being resolved by intact mass MS measurement.

| | | Crystallography | | | MS | ConSurf | |
|---|---|---|---|---|---|---|---|
| Res# | ResType | RSCC | ΔCont | Observation | PrimaryScore | n | % |
| 70 | G | 79.8 | 3 | | 2.15 | 7 | 3.7 |
| 70 | P! | 76.8 | 3 | | | 81 | 42.9 |
| 70 | A | 73.8 | 8.7 | | | 3 | 1.6 |
| 70 | MainCh | 87 | | | | | |
| 121 | S | 87.3 | 6.5 | | 1.59 | 103 | 57.9 |
| 121 | L | 80.8 | 0.0 | | 0.50 | 1 | 0.6 |
| 121 | A | 80.8 | 0.9 | | 1.67 | 11 | 6.2 |
| 121 | G | 79.9 | 1.7 | | 0.90 | 2 | 1.1 |
| 121 | N | 78.2 | 0.4 | | 1.34 | 7 | 3.9 |
| 121 | V | 77.8 | 2.3 | | | 1 | 0.6 |
| 121 | C | 75.5 | 2.5 | | | 0 | 0.0 |

| | | | | | | | |
|---|---|---|---|---|---|---|---|
| 121 | K | 73.0 | 1.0 | | | 4 | 2.2 |
| 121 | D | 72.0 | 3.9 | | | 12 | 6.7 |
| 121 | T | 68.1 | 0.7 | | | 17 | 9.6 |
| 121 | I | 67.4 | 2.0 | | | 1 | 0.6 |
| 121 | Y | 65.4 | 1.1 | | | 2 | 1.1 |
| 121 | E | 64.3 | 1.8 | | | 5 | 2.8 |
| 121 | H | 62.5 | 1.2 | | | 0 | 0.0 |
| 121 | R | 61.3 | 2.4 | | | 5 | 2.8 |
| 121 | P | 58.9 | 4.9 | | | 3 | 1.7 |
| 121 | Q! | 54.0 | 1.3 | | | 2 | 1.1 |
| 121 | MainCh | 91.8 | | | | | |

Supplementary Table 5: Residues of metalloproteinase resolved by crystallography and MS and/or phylogenetic analysis

| | | Crystallography | | | MS | ConSurf | |
|---|---|---|---|---|---|---|---|
| Res# | ResType | RSCC | ΔCont | RSCC* | PrimaryScore | n | % |
| 2 | F | 92.2 | 4.0 | 94.3 | 1.04 | 10 | 76.9 |
| 3 | S | 91.8 | 5.5 | 94.3 | 1.33 | 26 | 44.8 |
| 4 | P | 93.7 | 12.5 | 97.1 | 1.33 | 30 | 15.9 |
| 5 | R | 96.9 | 8.2 | 97.9 | 1.33 | 87 | 33.2 |
| 6 | Y | 97.3 | 5 | 98.1 | 3.03 | 226 | 85.3 |
| 7 | I | 96.8 | 3.8 | 97.7 | 3.03 | 75 | 28.3 |
| 8 | E | 97.7 | 4.6 | 98.3 | 3.03 | 247 | 93.6 |
| 10 | A | 95.2 | 12.1 | 97.7 | 3.03 | 15 | 5.7 |
| 13 | A | 95.3 | 9.9 | 97.8 | 3.03 | 161 | 60.8 |
| 16 | G | 92 | 21.3 | 97.8 | 3.03 | 24 | 9.0 |
| 17 | M | 96.5 | 4.5 | 97.9 | 3.03 | 67 | 24.7 |
| 18 | F | 95.6 | 4.6 | 96 | 3.03 | 140 | 52.2 |
| 20 | K | 97 | 9.5 | 96.8 | 3.03 | 149 | 55.6 |
| 21 | Y | 97.2 | 7.2 | 96.5 | 2.26 | 115 | 42.6 |
| 23 | S | 95.7 | 8.8 | 97.2 | 2.36 | 34 | 12.6 |
| 25 | L | 93.7 | 3.3 | 96.2 | 2.36 | 102 | 39.1 |
| 27 | T | 96.9 | 3.3 | 97.3 | 2.36 | 50 | 18.5 |
| 28 | I | 95.2 | 4 | 97.3 | 2.36 | 81 | 30.0 |
| 31 | R | 96.9 | 9.6 | 96.9 | 2.08 | 213 | 78.3 |
| 33 | H | 93.6 | 4.4 | 97.2 | 2.52 | 9 | 3.3 |
| 34 | E | 92.4 | 5.8 | 97.5 | 2.52 | 206 | 75.7 |
| 35 | M | 98.6 | 4.7 | 98.7 | 2.52 | 39 | 14.3 |
| 38 | T | 97.5 | 5.6 | 98.1 | 2.52 | 24 | 8.8 |
| 41 | G | 95.6 | 21.5 | 98 | 2.52 | 10 | 3.5 |
| 42 | F | 96.9 | 6.5 | 98.4 | 2.52 | 108 | 36.7 |
| 43 | Y | 98.4 | 6.4 | 98.5 | 1.36 | 257 | 86.8 |
| 44 | S | 91.5 | 8.1 | 97.4 | 1.36 | 4 | 1.4 |
| 45 | S | 96.6 | 11.6 | 97.8 | 1.36 | 33 | 11.1 |
| 48 | A | 97 | 11.1 | 98.2 | 1.36 | 2 | 0.7 |
| 50 | A | 94.4 | 13.9 | 95.4 | 1.36 | 4 | 1.3 |
| 51 | S | 97.1 | 7.5 | 97 | 1.36 | 9 | 3.0 |
| 52 | L | 97 | 4.6 | 97.4 | 1.36 | 291 | 98.0 |
| 53 | A | 96 | 8.4 | 98.2 | 1.36 | 5 | 1.7 |

| | | | | | | | |
|---|---|---|---|---|---|---|---|
| 54 | N | 96.9 | 3 | 97.5 | 1.36 | 4 | 1.3 |
| 58 | W | 98.3 | 12.2 | 97.8 | 1.36 | 294 | 99.0 |
| 59 | S | 95.8 | 7.8 | 96.5 | 1.36 | 146 | 49.2 |
| 63 | L | 97.1 | 3 | 97.6 | 1.82 | 43 | 14.5 |
| 64 | I | 96.7 | 6.9 | 97.9 | 1.82 | 171 | 57.6 |
| 68 | K | 95.3 | 10 | 98.1 | 1.82 | 16 | 5.4 |
| 70 | S | 97.6 | 7.9 | 98.2 | 1.65 | 31 | 10.4 |
| 72 | K | 93.6 | 4.6 | 97.3 | 1.65 | 3 | 1.0 |
| 73 | T | 98.2 | 4.7 | 97.6 | 2.26 | 256 | 85.6 |
| 74 | L | 96.9 | 3.2 | 98.1 | 2.26 | 290 | 97.0 |
| 75 | T | 96.7 | 5.7 | 97.2 | 2.26 | 11 | 3.7 |
| 76 | S | 96.7 | 8.8 | 97.3 | 2.26 | 79 | 26.4 |
| 77 | F | 98 | 8 | 97.8 | 2.26 | 294 | 98.3 |
| 78 | G | 91.1 | 29.5 | 97 | 2.26 | 47 | 15.7 |
| 79 | S | 85.5 | 4.3 | 95.7 | | 27 | 9.0 |
| 80 | W | 98.5 | 13.6 | 97.6 | 2.26 | 284 | 95.0 |
| 81 | R | 97.2 | 10.2 | 97.8 | 2.26 | 276 | 92.3 |
| 83 | R | 93.2 | 6 | 98.1 | 1.80 | 42 | 14.1 |
| 87 | P | 96.2 | 17 | 96.4 | 1.80 | 107 | 35.9 |
| 88 | R | 92.9 | 5.8 | 98 | 1.80 | 211 | 70.8 |
| 90 | S | 95.6 | 9.6 | 98.6 | 4.59 | 37 | 12.4 |
| 91 | H | 97.9 | 3.8 | 98.6 | 4.59 | 234 | 78.3 |
| 93 | H | 98.3 | 3.3 | 98.8 | 4.59 | 10 | 3.3 |
| 94 | A | 97.2 | 7.9 | 98.4 | 4.59 | 275 | 92.0 |
| 96 | L | 97.2 | 3.7 | 97.9 | 4.59 | 265 | 89.2 |
| 97 | L | 92.9 | 3.7 | 98 | 4.59 | 125 | 42.1 |
| 98 | T | 97.1 | 4.6 | 98.4 | 4.59 | 220 | 74.1 |
| 99 | T | 97.3 | 4.7 | 98.1 | 4.59 | 3 | 1.0 |
| 100 | I | 94.2 | 3.9 | 96.8 | 4.59 | 98 | 33.0 |
| 102 | F | 98.1 | 4.8 | 97.5 | 4.59 | 264 | 88.9 |
| 103 | D | 97.7 | 3.1 | 97.8 | 4.59 | 67 | 22.6 |
| 105 | Y | 95.4 | 4.5 | 97.4 | 4.59 | 1 | 0.3 |
| 107 | I | 96 | 4.3 | 98 | 4.59 | 118 | 39.6 |
| 108 | G | 92.8 | 13.9 | 98.2 | 4.59 | 296 | 99.3 |
| 109 | R | 96.7 | 7 | 98.2 | 4.59 | 10 | 3.4 |
| 111 | R | 94.5 | 7 | 97.3 | 3.04 | 6 | 2.0 |
| 112 | S | 93.2 | 2.9 | 96.1 | 1.19 | 2 | 0.7 |
| 113 | G | 94.5 | 10.6 | 98.2 | 1.19 | 79 | 26.5 |
| 114 | K | 96.6 | 5.2 | 98.8 | 1.19 | 1 | 0.3 |
| 115 | M | 98.2 | 4.4 | 97.3 | 1.19 | 220 | 73.8 |
| 116 | C | 97.9 | 10.3 | 97.9 | 1.19 | 298 | 100.0 |
| 118 | P | 95.5 | 18.3 | 97.4 | 1.19 | 68 | 22.9 |
| 121 | S | 98 | 9.1 | 98 | 2.37 | 293 | 98.7 |
| 123 | G | 93.6 | 9.2 | 98.3 | 2.37 | 227 | 76.7 |
| 129 | S | 96.2 | 6 | 97.5 | 2.37 | 194 | 65.5 |
| 130 | K | 86.9 | 3.3 | 96.9 | 2.37 | 25 | 8.5 |
| 133 | L | 96.6 | 4.9 | 97.9 | 2.97 | 102 | 34.6 |
| 134 | W | 97.6 | 8 | 98.7 | 2.97 | 1 | 0.3 |
| 136 | A | 97 | 15.3 | 97.8 | 2.97 | 266 | 89.9 |
| 139 | M | 98.7 | 5.3 | 98.2 | 2.97 | 159 | 53.7 |

| | | | | | | | |
|---|---|---|---|---|---|---|---|
| 138 | T | 97.5 | 6.4 | 98.2 | 2.97 | 245 | 82.8 |
| 140 | A | 97 | 12.7 | 98 | 2.97 | 282 | 95.3 |
| 141 | H | 97.1 | 4.8 | 98.2 | 2.97 | 296 | 100.0 |
| 143 | L | 96.6 | 3.7 | 97.4 | 2.97 | 121 | 40.9 |
| 144 | G | 96.5 | 13.3 | 98.8 | 2.97 | 295 | 99.7 |
| 145 | H | 98.3 | 5.7 | 98.2 | 2.97 | 296 | 100.0 |
| 149 | M | 98.2 | 4.9 | 98 | 2.97 | 222 | 75.3 |
| 151 | H | 96.8 | 5.9 | 98.5 | 2.97 | 292 | 99.0 |
| 155 | C | 90.7 | **2.1** | 96.4 | 2.97 | 289 | 98.6 |
| 156 | S | 96.4 | 8.5 | 97.3 | 2.97 | 32 | 10.9 |
| 157 | C | 91.9 | **1.6** | 97.6 | 2.97 | 239 | 87.9 |
| 158 | G | 96.7 | 15.1 | 98.6 | 2.97 | 53 | 18.3 |
| 159 | A | 97.7 | 16.5 | 98.4 | 2.97 | 97 | 33.9 |
| 160 | K | 95.5 | 7.5 | 98.5 | 2.97 | 18 | 6.3 |
| 161 | S | 97.9 | 9.5 | 97.7 | 2.21 | 58 | 20.5 |
| 162 | C | 93.2 | **2.7** | 97.6 | 2.21 | 281 | 98.9 |
| 163 | I | 97.4 | 8.9 | 97.6 | 2.21 | 198 | 69.7 |
| 164 | M | 98.8 | 5.6 | 98 | 2.21 | 280 | 98.6 |
| 165 | A | 96.8 | 9.5 | 98.1 | 2.21 | 129 | 45.6 |
| 166 | S | 95.8 | 8 | 96.2 | 2.21 | 21 | 7.4 |
| 168 | L | 97.9 | 3.4 | 96.9 | 2.21 | 65 | 24.0 |
| 169 | S | 95.9 | 10.5 | 97.3 | 2.21 | 85 | 30.4 |
| 170 | K | 94.9 | 7.2 | 97.2 | 2.21 | 6 | 2.2 |
| 172 | K | 92.9 | 6.5 | 97.3 | 2.54 | 2 | 0.7 |
| 174 | Y | 97.4 | 6.1 | 97.3 | 2.63 | 25 | 9.0 |
| 175 | A | 95.7 | 13 | 98.3 | 2.63 | 4 | 1.4 |
| 176 | F | 97.7 | 5.9 | 97.8 | 2.63 | 265 | 94.6 |
| 177 | S | 97.2 | 7.7 | 98.2 | 2.63 | 262 | 93.6 |
| 180 | S | 96.9 | 9 | 97.6 | 2.63 | 256 | 91.8 |
| 184 | Y | 97.4 | 6.8 | 97.9 | 2.63 | 56 | 20.2 |
| 187 | F | 96.9 | 6.5 | 97 | 2.63 | 93 | 33.8 |
| 191 | H | 94.9 | 5.4 | 98.6 | 1.50 | 10 | 3.7 |
| 192 | N | 97.7 | 3.1 | 98.2 | 1.50 | 30 | 11.1 |
| 193 | P | 95.7 | 12.3 | 97.6 | 1.50 | 108 | 39.9 |
| 195 | C | 91.1 | 3.5 | 97.4 | 1.50 | 271 | 100.0 |
| 196 | I | 98 | 6 | 98.4 | 1.50 | 62 | 23.0 |
| 197 | L | 95.9 | 4.2 | 97.4 | 1.50 | 99 | 37.9 |
| 200 | P | 91.6 | 16.9 | 92.8 | 0.82 | 246 | 100.0 |

Res# stands for residue number; ResType for residues type; RSCC for real-space correlation coefficient; ΔCont for contrast difference within other RSCC; * for main chain atoms; MS for mass spectrometry. RSCC in bold denotes cysteine partially occupied having ΔCont less than 3.0.

Supplementary Table 6: Residues of metalloproteinase having assignment ambiguity from crystallographic data being resolved by mass spectrometry

| | | Crystallography | | | MS | ConSurf | |
|---|---|---|---|---|---|---|---|
| Res# | ResType | RSCC | ΔCont | Observation | PrimaryScore | n | % |
| 1 | V | 78 | 1.2 | No H-bond | | | |
| 1 | G | 76.8 | 2.6 | | | | |
| 1 | T! | 74.2 | 0.9 | | 1.04 | | |
| 1 | I | 73.3 | 1.3 | | | | |

| | | | | | | | |
|---|---|---|---|---|---|---|---|
| 1 | A | 72 | 1.4 | | 1.33 | | |
| 1 | L | 70.6 | 0.1 | | | | |
| 1 | MainCh | 88.3 | | | | | |
| 9 | L! | 87.7 | 0.9 | | 3.03 | 211 | 79.9 |
| 9 | V | 86.8 | 0.8 | | | 8 | 3.0 |
| 9 | S | 86 | 2.1 | | 1.33 | 0 | 0.0 |
| 9 | I | 83.9 | 0.2 | | | 2 | 0.8 |
| 9 | A | 83.7 | 0.2 | | 1.04 | 0 | 0.0 |
| 9 | MainCh | 97.4 | | | | | |
| 11 | V! | 97.7 | 1.6 | | 3.03 | 64 | 24.2 |
| 11 | T | 96.1 | 3.2 | | | 0 | 0.0 |
| 11 | MainCh | 97.4 | | | | | |
| 12 | V! | 95.5 | 1.8 | | 3.03 | 256 | 96.6 |
| 12 | T | 93.7 | 2.4 | | | 0 | 0.0 |
| 12 | MainCh | 97 | | | | | |
| 14 | D! | 97.6 | 1.1 | 3 H-bonds | 3.03 | 263 | 99.2 |
| 14 | N | 96.5 | 4.7 | | 1.33 | 1 | 0.4 |
| 14 | L | 91.8 | 10.2 | | | 0 | 0.0 |
| 14 | MainCh | 97.7 | | | | | |
| 19 | V | 94.4 | 0.7 | | | 10 | 3.7 |
| 19 | T! | 93.7 | 6.5 | 2 H-bonds | 3.03 | 22 | 8.2 |
| 19 | MainCh | 95.7 | | | | | |
| 22 | N! | 92.6 | 2.7 | | 2.36 | 76 | 34.7 |
| 22 | S | 89.9 | 3.8 | 1 H-bond and exposed residue | | 17 | 7.8 |
| 22 | L | 86.1 | 2.1 | | | 0 | 0.0 |
| 22 | D | 84 | 3.3 | | 2.08 | 15 | 6.8 |
| 22 | MainCh | 96.5 | | | | | |
| 29 | S | 90.5 | 2.2 | | | 1 | 0.4 |
| 29 | V | 88.3 | 0.2 | | | 3 | 1.1 |
| 29 | A | 88.1 | 5.9 | | | 0 | 0.0 |
| 29 | T | 82.2 | 2.2 | | | 2 | 0.7 |
| 29 | Q | 80 | 1.3 | | | 22 | 8.1 |
| 29 | L | 78.7 | 0.4 | | | 9 | 3.3 |
| 29 | I | 78.3 | 0.4 | | | 9 | 3.3 |
| 29 | E | 77.9 | 1.1 | | 2.08 | 6 | 2.2 |
| 29 | K | 76.8 | 1.6 | Double occupancy | | 101 | 37.0 |
| 29 | R! | 74.5 | 2.1 | 1 H-bond and exposed residue | 2.36 | 103 | 37.7 |
| 29 | MainCh | 98 | | | | | |
| 30 | T! | 95.7 | 1.7 | 2 H-bonds | 2.08 | 45 | 16.5 |
| 30 | V | 94 | 3.4 | | | 4 | 1.5 |
| 30 | MainCh | 97.3 | | | | | |
| 32 | V! | 98.1 | 1.8 | Hydrophobic and buried region | 2.52 | 70 | 25.7 |
| 32 | T | 96.3 | 3.2 | | | 7 | 2.6 |
| 32 | MainCh | 97.7 | | | | | |
| 36 | V! | 97.2 | 1.2 | Hydrophobic and buried region | 2.52 | 126 | 46.2 |
| 36 | T | 96 | 4.5 | | | 4 | 1.5 |
| 36 | MainCh | 98.5 | | | | | |

| | | | | | | | |
|---|---|---|---|---|---|---|---|
| 37 | N! | 86 | 0.4 | | 2.52 | 255 | 93.8 |
| 37 | T | 85.6 | 0.2 | | | 0 | 0.0 |
| 37 | S | 85.4 | 0.2 | 1 H-bond and exposed residue | | 6 | 2.2 |
| 37 | V | 85.2 | 6.3 | | | 0 | 0.0 |
| 37 | L | 78.9 | 7.7 | | | 4 | 1.5 |
| 37 | D | 71.2 | 4.3 | | 1.36 | 1 | 0.4 |
| 37 | MainCh | 97.2 | | | | | |
| 39 | V! | 96.7 | 2.5 | Hydrophobic and buried region | 2.52 | 174 | 63.3 |
| 39 | T | 94.2 | 3.4 | | | 0 | 0.0 |
| 39 | MainCh | 98.8 | | | | | |
| 40 | N! | 94.7 | 1.7 | 3 H-bonds and exposed residue | 2.52 | 117 | 42.2 |
| 40 | L | 93 | 0.1 | | | 1 | 0.4 |
| 40 | D | 92.9 | 5.4 | | 1.36 | 146 | 52.7 |
| 40 | MainCh | 98.1 | | | | | |
| 46 | V! | 96.4 | 0.8 | Hydrophobic and buried region | 1.36 | 12 | 4.1 |
| 46 | T | 95.6 | 4.7 | | | 0 | 0.0 |
| 46 | MainCh | 97.7 | | | | | |
| 47 | D | 96.8 | 1.5 | 2 H-bonds and exposed residue | | 23 | 7.7 |
| 47 | N! | 95.3 | 3.9 | | 1.36 | 213 | 71.7 |
| 47 | L | 91.4 | 7.3 | | | 1 | 0.3 |
| 47 | MainCh | 97.7 | | | | | |
| 49 | D | 95.6 | 1.9 | 4 H-bonds and exposed residue | | 4 | 1.3 |
| 49 | N! | 93.7 | 0.4 | | 1.36 | 1 | 0.3 |
| 49 | L | 93.3 | 4.2 | | | 2 | 0.7 |
| 49 | MainCh | 98 | | | | | |
| 55 | L! | 96.2 | 2.5 | Hydrophobic and buried region | 1.36 | 228 | 76.8 |
| 55 | N | 93.7 | 1.1 | | | 0 | 0.0 |
| 55 | D | 92.6 | 7.8 | | | 0 | 0.0 |
| 55 | MainCh | 96.9 | | | | | |
| 56 | E | 96.3 | 2.1 | | | 268 | 90.2 |
| 56 | Q! | 94.2 | 2.2 | | 1.36 | 5 | 1.7 |
| 56 | MainCh | 97 | | | | | |
| 57 | V! | 97.3 | 2.7 | Hydrophobic and buried region | 1.36 | 135 | 45.6 |
| 57 | T | 94.6 | 1.4 | | | 1 | 0.3 |
| 57 | MainCh | 97.5 | | | | | |
| 60 | V | 85.5 | 0.9 | | 1.32 | 1 | 0.3 |
| 60 | I! | 84.6 | 0.7 | | 1.36 | 1 | 0.3 |
| 60 | T | 83.9 | 0.9 | | | 4 | 1.4 |
| 60 | S | 83 | 1.9 | | | 16 | 5.5 |
| 60 | A | 81.1 | 5.2 | | | 0 | 0.0 |
| 60 | MainCh | 96.7 | | | | | |
| 61 | S | 91.2 | 1.2 | Exposed residue | | 21 | 7.1 |
| 61 | K! | 90 | 4.1 | | 1.36 | 37 | 12.5 |
| 61 | MainCh | 97.6 | | | | | |
| 62 | D! | 97.7 | 1.6 | 4 H-bonds and exposed residue | 1.82 | 273 | 91.9 |

| | | | | | | | |
|---|---|---|---|---|---|---|---|
| 62 | N | 96.1 | 6.3 | | | 22 | 7.4 |
| 62 | L | 89.8 | 5.5 | | | 0 | 0.0 |
| 62 | MainCh | 98.2 | | | | | |
| 65 | S | 87.6 | 1 | | | 32 | 10.8 |
| 65 | I | 86.6 | 0.2 | | | 16 | 5.4 |
| 65 | K! | 86.4 | 0.3 | | 1.82 | 23 | 7.7 |
| 65 | MainCh | 97 | | | | | |
| 66 | V! | 98.4 | 1.9 | Hydrophobic and buried region | 1.82 | 191 | 64.3 |
| 66 | T | 96.5 | 3.6 | | | 3 | 1.0 |
| 66 | MainCh | 97.4 | | | | | |
| 67 | Q | 95.9 | 2.9 | | | 21 | 7.1 |
| 67 | E! | 93 | 5.7 | | 1.82 | 29 | 9.8 |
| 67 | MainCh | 97.5 | | | | | |
| 69 | D! | 96.1 | 1.6 | 3 H-bonds and exposed residue | 1.65 | 82 | 27.6 |
| 69 | N | 94.5 | 5.3 | | | 82 | 27.6 |
| 69 | L | 89.2 | 3.7 | | | 0 | 0.0 |
| 69 | MainCh | 96.9 | | | | | |
| 89 | I! | 96.6 | 1.7 | | 4.59 | 20 | 6.7 |
| 89 | T | 94.9 | 0.8 | | | 11 | 3.7 |
| 89 | V | 94.1 | 6.4 | | | 7 | 2.3 |
| | MainCh | 98.4 | | | | | |
| 92 | D! | 98.3 | 2.5 | Coordinating Ca2+ | 4.59 | 297 | 99.3 |
| 92 | N | 95.8 | 5 | | | 0 | 0.0 |
| 92 | L | 90.8 | 7.9 | | | 0 | 0.0 |
| 92 | MainCh | 97.8 | | | | | |
| 95 | Q! | 97.3 | 2.2 | | 4.59 | 263 | 88.0 |
| 95 | E | 95.1 | 5.2 | | | 0 | 0.0 |
| 95 | MainCh | 97.7 | | | | | |
| 101 | V! | 95.9 | 1.3 | Exposed residue | 4.59 | 8 | 2.7 |
| 101 | T | 94.6 | 3.5 | | | 21 | 7.0 |
| 101 | MainCh | 96.4 | | | | | |
| 104 | D | 95.4 | 0.3 | 1 H-bond and exposed residue | | 7 | 2.3 |
| 104 | N! | 95.1 | 1.7 | | 4.59 | 6 | 2.0 |
| 104 | L | 93.4 | 8 | | | 0 | 0.0 |
| 104 | MainCh | 96.6 | | | | | |
| 106 | V! | 95.6 | 2.5 | | 4.59 | 20 | 6.7 |
| 106 | T | 93.1 | 4.3 | | | 250 | 83.6 |
| 106 | MainCh | 97.8 | | | | | |
| 110 | S! | 96.2 | 2.2 | | 3.04 | 14 | 4.7 |
| 110 | T | 94 | 6.5 | | | 4 | 1.3 |
| 110 | MainCh | 96.4 | | | | | |
| 117 | D! | 97.2 | 0.9 | 3 H-bonds and exposed residue | 1.19 | 20 | 6.7 |
| 117 | N | 96.3 | 3.8 | | | 12 | 4.0 |
| 117 | L | 92.5 | 6.9 | | | 11 | 3.7 |
| 117 | MainCh | 97.9 | | | | | |

| | | | | | | | |
|---|---|---|---|---|---|---|---|
| 119 | S | 89.3 | 0.6 | | | 8 | 3.5 |
| 119 | K | 88.7 | 2.5 | | | 31 | 13.7 |
| 119 | Q | 80 | 1.2 | | | 8 | 3.5 |
| 119 | E! | 78.8 | 1.4 | | 1.19 | 73 | 32.3 |
| 119 | MainCh | 96.7 | | | | | |
| 120 | Q! | 94.1 | 1.7 | | 2.37 | 63 | 21.2 |
| 120 | E | 92.4 | 0.2 | | 1.22 | 6 | 2.0 |
| 120 | MainCh | 97.5 | | | | | |
| 122 | V! | 97 | 1.5 | Hydrophobic and buried region | 2.37 | 65 | 21.9 |
| 122 | T | 95.5 | 5.3 | | | 12 | 4.0 |
| 122 | MainCh | 97.5 | | | | | |
| 124 | V! | 97.1 | 2 | Hydrophobic and buried region | 2.37 | 194 | 65.5 |
| 124 | T | 95.1 | 5.2 | | | 0 | 0.0 |
| 124 | MainCh | 98.3 | | | | | |
| 125 | V! | 97.9 | 1.3 | Hydrophobic and buried region | 2.37 | 91 | 31.1 |
| 125 | T | 96.6 | 5.6 | | 1.19 | 4 | 1.4 |
| 125 | MainCh | 98.7 | | | | | |
| 126 | K | 91.7 | 3.3 | | 1.19 | 8 | 2.7 |
| 126 | R! | 88.4 | 0.6 | | 2.37 | 8 | 2.7 |
| 126 | MainCh | 97.7 | | | | | |
| 127 | D! | 97.6 | 1 | 4 H-bonds and exposed residue | 2.37 | 278 | 93.9 |
| 127 | N | 96.6 | 5.3 | | | 4 | 1.4 |
| 127 | L | 90.4 | 4.9 | | | 0 | 0.0 |
| 127 | MainCh | 97.8 | | | | | |
| 128 | H! | 98 | 2.8 | | 2.37 | 252 | 84.8 |
| 128 | F | 95.2 | 5.9 | | | 5 | 1.7 |
| 128 | MainCh | 96.9 | | | | | |
| 131 | L | 92.3 | 0.6 | 2 H-bonds and exposed residue | | 26 | 8.8 |
| 131 | D | 91.7 | 0.7 | | | 11 | 3.7 |
| 131 | N! | 91 | 5.6 | | 2.97 | 112 | 37.8 |
| 131 | MainCh | 93.7 | | | | | |
| 132 | N! | 96.5 | 1.7 | 3 H-bonds and exposed residue | 2.97 | 25 | 8.4 |
| 132 | L | 94.8 | 0.2 | | | 9 | 3.0 |
| 132 | D | 94.6 | 5.7 | | | 2 | 0.7 |
| 132 | MainCh | 97.2 | | | | | |
| 135 | V! | 97.1 | 1.9 | Hydrophobic and buried region | 2.97 | 164 | 55.4 |
| 135 | T | 95.2 | 2.7 | | | 4 | 1.4 |
| 135 | MainCh | 98 | | | | | |
| 137 | V! | 97.5 | 1.7 | Hydrophobic and buried region | 2.97 | 59 | 19.9 |
| 137 | T | 95.8 | 1.8 | | | 12 | 4.1 |
| 137 | MainCh | 98 | | | | | |
| 142 | E! | 97 | 2.9 | | 2.97 | 282 | 95.3 |
| 142 | Q | 94.1 | 6.9 | | | 13 | 4.4 |
| 142 | MainCh | 97.4 | | | | | |

| | | | | | | | |
|---|---|---|---|---|---|---|---|
| 148 | N | 93.4 | 0.5 | 1 H-bond and exposed residue | | 0 | 0.0 |
| 148 | D! | 92.9 | 2.5 | | 2.97 | 1 | 0.3 |
| 148 | L | 90.4 | 5.8 | | | 0 | 0.0 |
| 148 | MainCh | 96.7 | | | | | |
| 150 | H! | 95.2 | 2.7 | | 2.97 | 17 | 5.8 |
| 150 | F | 92.5 | 1.9 | | | 4 | 1.4 |
| 150 | MainCh | 97.4 | | | | | |
| 154 | T! | 96.7 | 2.4 | 1 H-bond and exposed residue | 2.97 | 11 | 3.8 |
| 154 | V | 94.3 | 3.4 | | | 5 | 1.7 |
| 154 | MainCh | 96.5 | | | | | |
| 167 | V! | 92.9 | 0 | Exposed residue | 2.21 | 15 | 5.4 |
| 167 | T | 92.9 | 0.1 | | | 28 | 10.0 |
| 167 | I | 92.8 | 6.7 | | | 5 | 1.8 |
| 167 | MainCh | 97.1 | | | | | |
| 171 | V | 95.9 | 0.8 | | | 33 | 12.0 |
| 171 | T! | 95.1 | 6.4 | | 2.54 | 14 | 5.1 |
| 171 | MainCh | 97.8 | | | | | |
| 173 | V | 90.1 | 1.6 | 1 H-bond and exposed residue | | 2 | 0.7 |
| 173 | S | 88.5 | 1.2 | | 2.63 | 36 | 12.9 |
| 173 | T! | 87.3 | 3.6 | | 0.82 | 0 | 0.0 |
| 173 | MainCh | 96.1 | | | | | |
| 179 | T | 88 | 4.6 | Double occupancy | | 0 | 0.0 |
| 179 | S | 83.4 | 0.9 | | | 0 | 0.0 |
| 179 | C! | 82.5 | 5.6 | | 2.63 | 276 | 98.6 |
| 179 | MainCh | 96.9 | | | | | |
| 186 | T! | 95.3 | 2.5 | 2 H-bonds and exposed residue | 2.63 | 33 | 12.0 |
| 186 | V | 92.8 | 3.6 | | | 10 | 3.6 |
| 186 | MainCh | 97 | | | | | |
| 188 | L! | 95.1 | 1 | Hydrophobic and buried region | 2.63 | 178 | 65.0 |
| 188 | N | 94.1 | 0.7 | | | 0 | 0.0 |
| 188 | D | 93.4 | 3.2 | | 0.82 | 0 | 0.0 |
| 188 | MainCh | 95.7 | | | | | |
| 189 | T! | 96.1 | 2.9 | 1 H-bond | 2.63 | 21 | 7.7 |
| 189 | I | 93.2 | 0.4 | | | 20 | 7.4 |
| 189 | V | 92.8 | 2.2 | | | 1 | 0.4 |
| 189 | MainCh | 96.4 | | | | | |
| 190 | S | 92.7 | 1.4 | | | 40 | 14.7 |
| 190 | K! | 91.3 | 4.2 | | 2.63 | 82 | 30.1 |
| 190 | MainCh | 98.3 | | | | | |
| 194 | Q! | 96.8 | 0.6 | | 1.50 | 48 | 17.7 |
| 194 | E | 96.2 | 10.8 | | | 6 | 2.2 |
| 194 | MainCh | 97.7 | | | | | |
| 198 | N! | 95.1 | 2.8 | Coordinating Ca2+ | 0.82 | 213 | 82.6 |
| 198 | D | 92.3 | 6.2 | | | 33 | 12.8 |

| | | | | | | | |
|---|---|---|---|---|---|---|---|
| 198 | L | 86.1 | 2.9 | | | 0 | 0.0 |
| 198 | MainCh | 95.9 | | | | | |
| 199 | K | 84.1 | 0.3 | | | 51 | 20.2 |
| 199 | E! | 83.8 | 0.9 | | 0.82 | 23 | 9.1 |
| 199 | A | 82.9 | 4.2 | | | 26 | 10.3 |
| 199 | L | 78.7 | 0.3 | | | 19 | 7.5 |
| 199 | S | 78.4 | 0 | | | 4 | 1.6 |
| 199 | Q | 78.4 | 0.4 | | | 2 | 0.8 |
| 199 | MainCh | 96.1 | | | | | |

Res# stands for residue number; ResType for residues type; RSCC for real-space correlation coefficient; ΔCont for contrast difference within other RSCC; MainCh for main chain atoms; MS for mass spectrometry; H-bond for hydrogen bond; ! for resolved residue.

Supplementary Table 7: Residues of metalloproteinase having ambiguity assignment from both crystallographic and mass spectrometry data but being resolved by phylogenetic analysis

| | | Crystallography | | | MS | ConSurf | |
|---|---|---|---|---|---|---|---|
| Res# | ResType | RSCC | ΔCont | Observation | PrimaryScore | n | % |
| 15 | N! | 96 | 0.2 | 3 H-bonds | 2.99 | 124 | 46.8 |
| 15 | D | 95.8 | 2.8 | | 3.03 | 3 | 1.1 |
| 15 | L | 93 | 7.8 | | | 0 | 0.0 |
| 15 | MainCh | 97.3 | | | | | |
| 71 | N! | 89.5 | 1.8 | Density looks like | 1.05 | 59 | 19.8 |
| 71 | D | 87.7 | 3 | is covalently bound | | 42 | 14.1 |
| 71 | K | 81.4 | 1.8 | | | 7 | 2.3 |
| 71 | R | 77.5 | 0.6 | | 1.65 | 7 | 2.3 |
| 71 | MainCh | 97.1 | | | | | |
| 82 | E! | 96.3 | 3 | | 1.80 | 66 | 22.1 |
| 82 | Q | 93.3 | 3.4 | | 1.05 | 48 | 16.1 |
| 82 | MainCh | 97.4 | | | | | |
| 84 | N | 91.9 | 0.8 | 4 H-bonds and exposed | 1.15 | 21 | 8.7 |
| 84 | D! | 91.1 | 4.3 | | 1.80 | 70 | 28.9 |
| 84 | L | 78.8 | 2 | | | 0 | 0.0 |
| 84 | MainCh | 96.5 | | | | | |
| 85 | L! | 96.5 | 5 | Hydrophobic and buried | 1.15 | 284 | 95.0 |
| 85 | N | 91.5 | 1 | | | 0 | 0.0 |
| 85 | D | 90.5 | 6.8 | | 1.80 | 0 | 0.0 |
| 85 | MainCh | 97.5 | | | | | |
| 86 | L! | 96.7 | 5.3 | | 1.80 | 209 | 70.1 |
| 86 | N | 91.4 | 1.3 | | 1.15 | 1 | 0.3 |
| 86 | D | 90.1 | 6.3 | | | 0 | 0.0 |
| 86 | MainCh | 97.5 | | | | | |
| 146 | N! | 98 | 0.6 | 4 H-bonds and buried | 2.26 | 275 | 92.9 |
| 146 | D | 97.4 | 4 | | 2.97 | 1 | 0.3 |
| 146 | L | 93.4 | 0.1 | | | 0 | 0.0 |
| 146 | MainCh | 97.2 | | | | | |
| 147 | L! | 96.4 | 3.5 | Hydrophobic and buried | 2.97 | 184 | 62.2 |

| | | | | | | | |
|---|---|---|---|---|---|---|---|
| 147 | N | 92.9 | 0.8 | | 2.26 | 0 | 0.0 |
| 147 | D | 92.1 | 5.2 | | | 0 | 0.0 |
| 147 | MainCh | 97.8 | | | | | |
| 152 | D! | 97.3 | 2.9 | 3 H-bonds and buried | 2.97 | 288 | 98.0 |
| 152 | N | 94.4 | 6.1 | | 2.26 | 2 | 0.7 |
| 152 | L | 88.3 | 4.7 | | | 0 | 0.0 |
| 152 | MainCh | 98 | | | | | |
| 153 | D! | 95.6 | 0.2 | 3 H-bonds and exposed | 2.97 | 52 | 17.7 |
| 153 | N | 95.4 | 0.2 | | | 27 | 9.2 |
| 153 | L | 95.2 | 4.3 | | 2.26 | 10 | 3.4 |
| 153 | MainCh | 96.2 | | | | | |
| 178 | V | 87.2 | 1.1 | | 2.54 | 0 | 0.0 |
| 178 | T! | 86.1 | 1.9 | | 2.63 | 21 | 7.5 |
| 178 | MainCh | 96.4 | | | | | |

Res# stands for residue number; ResType for residues type; RSCC for real-space correlation coefficient; ΔCont for contrast difference within other RSCC; MainCh for main chain atoms; MS for mass spectrometry; H-bond for hydrogen bond; ! for resolved residue.

Supplementary Table 8: Residues of metalloproteinase having ambiguity assignment from crystallographic, mass spectrometry and phylogenetic analysis data

| | | Crystallography | | | MS | ConSurf | |
|---|---|---|---|---|---|---|---|
| Res# | ResType | RSCC | ΔCont | Observation | PrimaryScore | n | % |
| 24 | N? | 97.4 | 2 | 2 H-bonds | 2.36 | 88 | 32.6 |
| 24 | D? | 95.4 | 4.2 | | 2.21 | 120 | 44.4 |
| 24 | L | 91.2 | 7.3 | | | 1 | 0.4 |
| 24 | MainCh | 96.8 | | | | | |
| 26 | N? | 88 | 4.7 | | 2.36 | 35 | 13.3 |
| 26 | L | 83.3 | 0.7 | 1 H-bond and exposed residue | | 1 | 0.4 |
| 26 | D? | 82.6 | 1.8 | | 2.24 | 31 | 11.8 |
| 26 | S | 80.8 | 0.3 | | | 6 | 2.3 |
| 26 | MainCh | 96.9 | | | | | |
| 181 | Q? | 96.1 | 4.7 | | 2.54 | 38 | 13.6 |
| 181 | E? | 91.4 | 1.2 | | 2.63 | 17 | 6.1 |
| 181 | MainCh | 97.7 | | | | | |
| 182 | N? | 89.8 | 1.2 | 2 H-bonds and exposed residue | 2.54 | 22 | 7.9 |
| 182 | L | 88.6 | 2 | | | 1 | 0.4 |
| 182 | D? | 84.2 | 2.8 | | 2.63 | 14 | 5.0 |
| 182 | MainCh | 97.2 | | | | | |
| 183 | Q? | 95.8 | 0 | | 2.54 | 52 | 18.6 |
| 183 | E? | 95.8 | 7.2 | | 2.63 | 46 | 16.4 |
| 183 | MainCh | 97.4 | | | | | |
| 185 | Q? | 92 | 3.2 | | 2.63 | 50 | 18.1 |
| 185 | E? | 88.8 | 1.4 | | 2.54 | 68 | 24.5 |
| 185 | MainCh | 96.8 | | | | | |

Res# stands for residue number; ResType for residues type; RSCC for real-space correlation coefficient; ΔCont for contrast difference within other RSCC; MainCh for main chain atoms; MS for mass spectrometry; H-bond for hydrogen bond; ! for resolved residue.

Supplementary Table 9: Residues of MjTX-I resolved by crystallography and MS and/or phylogenetic analysis

| | | Cryst mon ABCD | | Cryst mon A | | Cryst mon B | | Cryst mon C | | Cryst mon D | | MS | ConSurf | |
|---|---|---|---|---|---|---|---|---|---|---|---|---|---|---|
| Res# | ResType | RSCC | ΔCont | RSCC | ΔCont | RSCC | ΔCont | RSCC | ΔCont | RSCC | ΔCont | PrimaryScore | #Struct | % |
| 1 | S | 90.5 | 10.6 | 91.5 | 6.4 | 94.9 | 4.4 | 93.3 | 11.4 | | | 3.49 | 40 | 100.0 |
| 2 | L | 91.1 | 13.1 | 93.6 | 3.8 | | | 85.2 | 11.7 | | | 3.49 | 114 | 78.1 |
| 3 | V | 86.9 | 9.6 | | | | | 87.9 | 4.6 | 79 | 10.2 | 3.49 | 25 | 14.5 |
| 5 | L | 92.5 | 5 | 95.2 | 5.4 | 92.9 | 3.6 | 89.7 | 4.1 | 89.6 | 5.5 | 3.49 | 95 | 47.0 |
| 6 | G | 85.7 | 21.7 | 94 | 29.4 | 92.4 | 18.8 | 69.4 | 4.9 | 81 | 32.1 | 3.49 | 62 | 29.4 |
| 7 | K | | | | | 91.6 | 6.0 | | | | | | 63 | 29.6 |
| 8 | M | 95.3 | 5 | | | | | 95.4 | 5.7 | 87.8 | 4.6 | 3.49 | 171 | 79.5 |
| 9 | I | 93.6 | 3.6 | | | | | 94.8 | 4.4 | 91.2 | 7 | 3.49 | 161 | 74.2 |
| 10 | L | 90.3 | 7 | 91.7 | 3.1 | 92 | 4.4 | | | 84.2 | 5.1 | 3.49 | 11 | 5.0 |
| 11 | Q | 91.8 | 3.6 | | | | | | | 88.5 | 9.3 | 3.49 | 24 | 11.0 |
| 12 | E | 94.4 | 3.4 | | | | | 93.2 | 5.1 | 93.8 | 5.9 | 3.49 | 9 | 4.0 |
| 13 | T | 94.8 | 5.5 | 96.7 | 5.6 | 93.8 | 6 | 95 | 5.7 | 94.2 | 5.7 | 3.49 | 176 | 72.7 |
| 14 | G | 86.5 | 31.3 | 89.7 | 21.5 | 90.8 | 28.8 | 78 | 18 | 83.3 | 34.8 | 3.49 | 198 | 77.6 |
| 15 | K | | | | | 88 | 4.1 | | | 88.4 | 3.8 | 3.49 | 102 | 36.8 |
| 16 | N | 89.5 | 5.2 | 91.6 | 4.3 | | | 91 | 5.5 | 88.5 | 5.3 | 2.45 | 72 | 25.6 |
| 17 | P | 91.8 | 14 | 92.4 | 12.8 | 95.9 | 19.9 | 92.1 | 11.7 | 87.4 | 1.1 | 3.84 | 141 | 49.3 |
| 20 | S | 91 | 10.1 | 86.8 | 3.5 | 92.6 | 10.3 | 91.2 | 5.1 | 94.4 | 8.6 | 3.84 | 75 | 25.3 |
| 21 | Y | 95.6 | 8.7 | 96.1 | 9.3 | 95.8 | 7.3 | 94.8 | 5.4 | 96 | 10.7 | 3.84 | 248 | 83.5 |
| 22 | G | 77.4 | 25.4 | 81.3 | 24.9 | 86.9 | 24.3 | 70.5 | 11.2 | 51.2 | 3 | 3.84 | 41 | 13.9 |
| 23 | A | 89.8 | 10.7 | 91.4 | 17.3 | 93.7 | 8.4 | 83.6 | 12.9 | 87.5 | 23.1 | 3.84 | 13 | 4.4 |
| 24 | Y | 95.9 | 6 | 95.6 | 6.8 | 95.6 | 5.2 | 96.1 | 5.4 | 95.4 | 4.3 | 3.84 | 297 | 99.7 |
| 25 | G | 85.3 | 10.7 | 92.6 | 18.1 | 91.6 | 11.3 | 83.9 | 14.4 | 77.5 | 21.1 | 3.84 | 297 | 99.7 |
| 26 | C | 96.7 | 14.6 | 97.6 | 16 | 97.4 | 11.3 | 95.4 | 15.5 | 95.9 | 15.7 | 3.84 | 295 | 98.7 |
| 27 | N | 95.8 | 13.5 | 97.3 | 11.4 | 95.6 | 9.4 | 94.2 | 12.9 | 95.6 | 13.3 | 3.73 | 10 | 3.3 |
| 28 | C | 96.5 | 7 | 96.9 | 6.4 | 96.2 | 5.7 | 95.7 | 8.1 | 96.7 | 6 | 3.84 | 300 | 100.0 |
| 29 | G | 82.5 | 23 | 87 | 28.4 | 87.8 | 11.4 | 73.4 | 16.3 | 74.9 | 23.3 | 3.84 | 294 | 98.0 |
| 30 | V | 92.6 | 4.8 | | | | | 93.5 | 4.2 | 85.8 | 4.4 | 3.84 | 17 | 5.7 |
| 31 | L | 86.5 | 4.6 | | | 93.6 | 3.7 | | | 79.3 | 4.7 | 3.84 | 5 | 1.7 |
| 32 | G | 82.1 | 35.6 | 93 | 27.3 | 89.2 | 18.2 | 69 | 20.2 | 60.1 | 41.3 | 3.84 | 291 | 97.0 |
| 33 | R | 91.4 | 6.1 | 93.7 | 11 | 93 | 3.4 | 91.5 | 5.7 | | | 3.84 | 61 | 20.3 |
| 34 | G | 88.5 | 23.5 | 90.4 | 22.6 | 91.9 | 17.5 | 80.3 | 23.3 | 89 | 28.1 | 3.42 | 278 | 92.7 |

| | | | | | | | | | | | | | | |
|---|---|---|---|---|---|---|---|---|---|---|---|---|---|---|
| 36 | P | 91.7 | 13.2 | 93.5 | 14.7 | 89.8 | 8.9 | 92.6 | 4.2 | 91.2 | 20.6 | 3.42 | 290 | 96.7 |
| 37 | K | 89.7 | 8 | 94.2 | 7.2 | 92.4 | 6.1 | | | 93.3 | 8.4 | 3.42 | 77 | 25.7 |
| 38 | D | | | | | | | 96 | 4.3 | | | 1.38 | 299 | 99.7 |
| 39 | A | 88.7 | 13.1 | 88.3 | 7.7 | 94 | 18.5 | 83.3 | 12 | 89.8 | 8.6 | 1.38 | 120 | 40.0 |
| 40 | T | | | | | 93.8 | 3.3 | | | | | 1.38 | 140 | 46.7 |
| 41 | D | | | | | | | | | 93.8 | 3.6 | 1.38 | 298 | 99.3 |
| 42 | R | | | 90.8 | 4.1 | 88.6 | 6.8 | | | | | 1.38 | 169 | 56.3 |
| 43 | C | 95.9 | 14.4 | 97.7 | 16.3 | 92.3 | 9.1 | 94.8 | 13 | 97.6 | 13.7 | 1.65 | 300 | 100.0 |
| 44 | C | 96.8 | 12.1 | 97 | 9.1 | 97 | 14.3 | 96 | 10 | 97.3 | 13.3 | 1.65 | 300 | 100.0 |
| 45 | Y | 92.4 | 8.1 | 94.1 | 4.4 | 92.9 | 10.9 | 89.7 | 4.7 | | | 1.65 | 12 | 4.0 |
| 46 | V | 92.6 | 7.8 | 93.4 | 4.8 | 92.2 | 4.1 | 93.6 | 5.3 | 94.3 | 7.3 | 1.65 | 81 | 27.0 |
| 47 | H | 95.5 | 3.2 | 97.1 | 3.8 | 95.8 | 3.3 | 94.5 | 3 | | | 1.65 | 298 | 99.3 |
| 48 | K | 95.1 | 14.5 | 94.7 | 13.8 | 94.9 | 11.4 | 96.3 | 15.3 | 94.5 | 11.9 | 1.65 | 7 | 2.3 |
| 49 | C | 94.7 | 11.8 | 96 | 11.6 | 94.5 | 9 | 96.3 | 12.6 | 93.6 | 5.8 | 0.86 | 175 | 58.3 |
| 50 | C | 95 | 31.9 | 97.2 | 36 | 96.2 | 14.9 | 94.2 | 20.5 | 92.8 | 10.4 | 0.86 | 300 | 100.0 |
| 51 | Y | 94.3 | 13.8 | 95.2 | 13.5 | 95.5 | 15.1 | 95.1 | 11.5 | 88.9 | 6.1 | 0.86 | 286 | 95.3 |
| 54 | L | 88.7 | 4.6 | | | 88.6 | 6.5 | | | 87.6 | 3.6 | 2.25 | 118 | 39.3 |
| 55 | T | | | | | 90.6 | 3.6 | | | | | 2.25 | 30 | 10.2 |
| 57 | C | 91.8 | 8.9 | 94 | 10.7 | 93.2 | 8.6 | 85.9 | 6.6 | 90.2 | 4.7 | | 295 | 98.3 |
| 58 | N | 85.4 | 5.8 | | | | | 83.8 | 4.8 | 88.9 | 7.8 | 2.25 | 70 | 23.6 |
| 59 | P | 90.1 | 10.2 | | | 94.1 | 11.7 | 89.5 | 8.9 | 90.6 | 16.2 | 2.25 | 191 | 64.1 |
| 60 | K | | | | | | | | | 70.2[α] | 4.4 | 2.25 | 127 | 42.5 |
| 61 | K | | | | | 90.9 | 7.1 | | | | | | 8 | 2.7 |
| 62 | D | 92 | 5.6 | 93.9 | 4.4 | 94.1 | 3.9 | 92.3 | 3.3 | 88.6 | 4.1 | 2.42 | 61 | 20.5 |
| 63 | R | 83.9 | 3.9 | | | | | 90.1 | 3.7 | 81.2 | 4.6 | 2.42 | 36 | 12.1 |
| 64 | Y | 94.6 | 6.4 | 95.3 | 5.6 | 95.6 | 3.3 | 93.5 | 8.1 | 89.5 | 6.1 | 2.42 | 281 | 94.3 |
| 65 | S | 85.7 | 9.6 | | | 87.1 | 7 | 86.4 | 6.7 | 76.6 | 7.8 | 2.42 | 89 | 29.9 |
| 66 | Y | 94.3 | 6.8 | 94.4 | 6.4 | 95.5 | 7.4 | 92.6 | 5.3 | 93.8 | 8.4 | 2.42 | 143 | 48.0 |
| 67 | S | 84.7 | 3.5 | | | 93 | 4.4 | 85.6 | 7.2 | | | 2.42 | 116 | 39.1 |
| 68 | W | 94.6 | 13.4 | 95.1 | 10 | 96.4 | 8.9 | 92.6 | 9.7 | 92.6 | 12 | 2.42 | 17 | 5.7 |
| 71 | K | 91.4 | 11.1 | 93.3 | 10.4 | 94 | 9.6 | 89.6 | 5.8 | 92.3 | 8.9 | 2.42 | 38 | 12.9 |
| 72 | T | 92.4 | 6.1 | | | | | 91 | 3.7 | 90.9 | 7 | 2.51 | 62 | 21.0 |
| 73 | I | 92.9 | 6.9 | | | 93.2 | 6.7 | 93 | 5.3 | 88.9 | 11.6 | 2.51 | 146 | 49.0 |
| 74 | V | 89.1 | 4 | 92.6 | 4.9 | | | 86.8 | 5.9 | | | 2.51 | 55 | 18.5 |
| 75 | C | 93.6 | 17.5 | 98.1 | 16 | 97.4 | 25.4 | 83 | 2.8 | 93 | 11.7 | 2.51 | 296 | 99.7 |
| 76 | G | 75.6 | 19.2 | 85.7 | 28.8 | 80.8 | 11 | | | | | 2.51 | 80 | 26.8 |

| | | | | | | | | | | | | | | |
|---|---|---|---|---|---|---|---|---|---|---|---|---|---|---|
| 79 | N | 86.3 | 4.6 | | | 89.6 | 4.2 | | | | | 1.77 | 65 | 21.8 |
| 81 | C | 92 | 14.1 | 93.3 | 13.3 | 94.7 | 13.2 | 93 | 16.7 | 84 | 9.2 | 2.51 | 292 | 98.0 |
| 82 | L | 90 | 4.1 | 92.1 | 3.5 | | | | | | | 2.51 | 12 | 4.1 |
| 83 | K | 83.5 | 6.1 | 93.4 | 9.1 | | | 88.9 | 8.6 | 88.3 | 8.4 | 2.51 | 81 | 27.3 |
| 85 | L | 91.7 | 4.7 | 95.6 | 4.1 | | | 90.8 | 5.3 | 88[α] | 3.9 | 2.52 | 87 | 29.3 |
| 86 | C | 93.3 | 15.5 | 97.2 | 16.6 | 95.6 | 15.9 | 87.4 | 10 | 91.1 | 13.1 | 2.52 | 297 | 100.0 |
| 87 | E | 94 | 6 | | | 95.2 | 5.4 | 91.3 | 5.4 | 90.4 | 6.9 | 2.52 | 151 | 50.8 |
| 88 | C | 96.3 | 17.2 | 97.7 | 12.5 | 97.3 | 17.1 | 96.8 | 12.3 | 92.8 | 16.9 | 2.52 | 296 | 99.7 |
| 90 | K | 92.5 | 6.2 | 93.7 | 9.4 | 93.3 | 3.7 | 93.7 | 8.6 | | | 2.52 | 107 | 36.0 |
| 91 | A | 89.1 | 16.2 | 94.9 | 10.6 | 82.2 | 5.2 | 92.7 | 12.1 | 86.5 | 12.2 | 2.52 | 78 | 26.4 |
| 92 | V | | | | | | | 92.3 | 3.1 | 92.8 | 3.9 | 2.52 | 32 | 10.8 |
| 93 | A | 93.4 | 15.4 | 95.2 | 5.3 | 95 | 13 | 90.8 | 13.2 | 87.1 | 14.9 | 2.52 | 248 | 84.1 |
| 94 | I | 93.2 | 5 | 95.8 | 3.7 | | | 94.5 | 4.4 | 89.3 | 4.7 | 2.52 | 53 | 18.0 |
| 95 | C | 95.6 | 13.4 | 96.1 | 14.4 | 96.1 | 8.2 | 97 | 14 | 97.8 | 21 | 2.52 | 295 | 100.0 |
| 96 | L | 94 | 6.4 | 93.9 | 4.6 | 92.9 | 10.6 | 94.5 | 6.9 | 95.4 | 4.7 | 2.52 | 82 | 27.9 |
| 97 | R | 89.9 | 4.9 | 94.9 | 6.9 | 91.7 | 5.6 | 86.7 | 6.2 | | | 2.52 | 64 | 22.1 |
| 98 | E | | | 95.2 | 3.1 | 87.4 | 3.7 | | | | | 2.37 | 27 | 9.9 |
| 99 | N | | | | | 91.9 | 3.5 | 95.4 | 3.1 | | | 2.37 | 97 | 38.8 |
| 100 | L | | | 93.8 | 3.6 | | | 89.3 | 3.1 | | | 2.37 | 79 | 56.0 |
| 102 | T | 91.1 | 4.9 | 92.6 | 5.1 | 89.8 | 8.8 | | | 92.8 | 3.5 | 2.37 | 108 | 45.4 |
| 103 | Y | 94.9 | 6 | 96.1 | 6.5 | 93.7 | 3.6 | 95.8 | 6.2 | 94.2 | 4.5 | 2.37 | 216 | 91.1 |
| 104 | N | | | | | 93.6 | 4.2 | 94.8 | 3.9 | | | 2.37 | 135 | 57.2 |
| 105 | K | | | 93 | 8.8 | | | | | | | 2.37 | 85 | 36.3 |
| 106 | K | | | 88.5 | 8 | | | | | | | 2.85 | 91 | 39.7 |
| 107 | Y | 87.4 | 4.9 | 91.7 | 6.4 | | | | | 94.4 | 7 | 2.85 | 150 | 67.0 |
| 108 | K | | | 78.1 | 5.3 | | | | | | | 2.85 | 63 | 30.9 |
| 111 | Y | | | 85.4 | 4 | | | | | | | 3.09 | 5 | 4.0 |
| 114 | P | | | | | | | 73.5 | 8.6 | 76.3 | 7.8 | 3.09 | 17 | 14.4 |
| 115 | F | | | 89.1 | 6.8 | | | | | 89.4 | 5.6 | 3.09 | 10 | 8.8 |
| 117 | D | | | | | | | 87.0 | 3.3 | | | | 3 | 3.9 |
| 116 | C | 95.3 | 14.1 | 97.1 | 16.6 | 96.4 | 15 | 94.8 | 12.8 | 95.4 | 16.3 | 3.09 | 109 | 99.1 |
| 118 | K | 94.8 | 9.4 | 96.1 | 10.2 | 96.3 | 8.7 | 91.3 | 6 | 95.4 | 9 | 1.3 | 11 | 20.8 |
| 119 | A | 80.8 | 7.7 | 81.6 | 10.6 | 85.6 | 4.8 | 69.6 | 4.7 | 89.5 | 9 | 1.3 | 7 | 15.6 |
| 122 | C | 89.7 | 16.1 | 94 | 21.9 | 84.7 | 12.3 | 93.8 | 14.9 | 91.2 | 16.9 | 1.3 | 16 | 94.1 |

Res# stands for residue number; ResType for residues type; Cryst for crystallography, mon for monomer, RSCC for real-space correlation coefficient; ΔCont for contrast difference within other RSCC; MS for mass spectrometry. RSCC numbers with $^{\alpha}$ refers to RSCC of main-chain being between 70 and 80, and $^{\beta}$ refers to values below 70.

Supplementary Table 10: Residues of MjTX-I having assignment ambiguity from crystallographic data being resolved by mass spectrometry

| | | Cryst mon ABCD | | Cryst mon A | | Cryst mon B | | Cryst mon C | | Cryst mon D | | MS | ConSurf | |
|---|---|---|---|---|---|---|---|---|---|---|---|---|---|---|
| Res# | ResType | RSCC | ΔCon | RSCC | ΔCon | RSCC | ΔCon | RSCC | ΔCon | RSCC | ΔCon | PrimScore | #Struct | % |
| 1 | S! | | | | | | | | | 75.9 | 2.4 | 3.49 | 40 | 100 |
| 1 | MainCh | | | | | | | | | 90.8 | | | | |
| 2 | L! | | | | | 92.5 | 2.2 | | | 89.6 | 1.5 | 3.49 | 114 | 78.1 |
| 2 | MainCh | | | | | 95.7 | | | | 90.9 | | | | |
| 3 | V! | | | 85.1 | 1.5 | 94.4 | 0.1 | | | | | 3.49 | 25 | 14.5 |
| 3 | MainCh | | | 94.8 | | 95.5 | | | | | | | | |
| 4 | E! | 92.7 | 2 | 92.9 | 6.8 | 95.4 | 0.6 | 95.3 | 2 | 83.9 | 0.7 | 3.49 | 47 | 25 |
| 4 | Q | 90.7 | 5.6 | 93.2 | 0.3 | 94.8 | 5.3 | 93.3 | 4.5 | 81 | 0.5 | 1.25 | 90 | 47.9 |
| 4 | MainCh | 93.8 | | 95 | | 95.4 | | 92.4 | | 92 | | | | |
| 7 | S | 81.9 | 1.3 | 82.1 | 1 | | | 81.5 | 4.4 | 76 | 2.9 | | 27 | 12.7 |
| 7 | K! | 73.9 | 1.4 | 89.8 | 1.8 | | | 63.4* | 1.2 | n/a* | n/a | 3.49 | 63 | 29.6 |
| 7 | MainCh | 92.1 | | 92.4 | | | | 89.2 | | 89 | | | | |
| 8 | M! | | | 96.2 | 2.8 | 97 | 0.1 | | | | | 3.49 | 171 | 79.5 |
| 8 | MainCh | | | 94.2 | | 97.5 | | 91.3 | | | | | | |
| 9 | I! | | | 94 | 0.5 | 94.3 | 2.7 | | | | | 3.49 | 161 | 74.2 |
| 9 | MainCh | | | 95.8 | | 94.9 | | 93 | | | | | | |
| 10 | L! | | | | | | | 83.2 | 2.3 | | | 3.49 | 11 | 5 |
| 10 | MainCh | | | 95.5 | | 93.8 | | 92.5 | | | | | | |
| 11 | Q! | | | 94.3 | 0 | 92.5 | 2.6 | 89.3 | 1.6 | | | 3.49 | 24 | 11 |
| 11 | E | | | 94.3 | 4.8 | 88.9 | 3.6 | 87.7 | 3.3 | | | 1.27 | 10 | 4.6 |

| | | | | | | | | | | | | | | |
|---|---|---|---|---|---|---|---|---|---|---|---|---|---|---|
| 11 | MainCh | | | 95.7 | | 94.1 | | 94 | | | | | | |
| 12 | E! | | | 94.1 | 0.8 | 96 | 2.5 | | | | | 3.49 | 9 | 4 |
| 12 | Q | | | 93.3 | 3.9 | 93.5 | 3 | | | | | | 0 | 0 |
| 12 | MainCh | | | 93.5 | | 94 | | 93.4 | | | | | | |
| 15 | K! | 84.9 | 0.7 | 83.1 | 1.3 | | | 89.1 | 2.5 | | | 3.49 | 102 | 36.8 |
| 15 | MainCh | 93.5 | | 93.6 | | | | 92.8 | | | | | | |
| 18 | A | 78 | 0.9 | 83.1 | 1.2 | 89.2 | 5.7 | 67.9 | 2.4 | 78.4 | 2 | 2.6 | 10 | 5.3 |
| 18 | V! | 77.1 | 2.9 | 86.1 | 0.1 | 83.5 | 4.2 | 71.6 | 0 | 76.4 | 3.3 | 3.84 | 29 | 15.4 |
| 18 | T | 74.2 | 0.6 | 86 | 1.1 | 76.8 | 3.5 | 71.6 | 0 | 73.1 | 2.2 | 2.19 | 8 | 4.3 |
| 18 | I | 72.9 | 0.5 | 81.9 | 4.2 | 73.3 | 3.5 | 74.5 | 2.9 | 70.3 | 0.9 | 2.44 | 35 | 18.6 |
| 18 | MainCh | 90.8 | | 92.5 | | 94.5 | | 89.1 | | 88 | | | | |
| 19 | V | 80.3 | 0.2 | 77.9 | 0.5 | 87.1 | 1.1 | 85.8 | 0.9 | 77.5 | 3.1 | | 11 | 3.8 |
| 19 | T! | 80.1 | 0.6 | 80.7 | 2.8 | 86 | 0.8 | 84.9 | 1.7 | 79.8 | 2.3 | 3.84 | 19 | 6.5 |
| 19 | S | 79.5 | 0.6 | 83.7 | 3 | 81.7 | 2 | 80.9 | 2.5 | 83.7 | 3.9 | 1.24 | 11 | 3.8 |
| 19 | I | 78.9 | 4.4 | 86.2 | 2.5 | 84 | 2.3 | 83.2 | 2.3 | 74.4 | 1.3 | | 10 | 3.4 |
| 19 | K | 61.8 | 1.8 | 65.4 | 0.6 | 66.3 | 6.7 | 66.7 | 0.8 | 70.7 | 5.6 | 2.6 | 11 | 3.8 |
| 19 | MainCh | 82.7 | | 90.9 | | 77.1 | | 82.3 | | 87.2 | | | | |
| 30 | V! | | | 95.6 | 1.4 | 91.8 | 6.1 | | | | | 3.84 | 17 | 5.7 |
| 30 | T | | | 94.2 | 5.9 | 91.9 | 0.1 | | | | | 1.51 | 4 | 1.3 |
| 30 | MainCh | | | 94.4 | | 95.7 | | | | | | | | |
| 31 | L! | | | 88 | 1.2 | | | 84.2 | 1.9 | | | 3.84 | 5 | 1.7 |
| 31 | N | | | 86.8 | 0.4 | | | 82.3 | 5.9 | | | | 1 | 0.3 |
| 31 | D | | | 86.4 | 5.8 | | | 70.8 | 1 | | | 1.51 | 2 | 0.7 |
| 31 | MainCh | | | 95.7 | | | | 91 | | | | | | |
| 33 | R! | | | | | | | | | 87.3 | 0.8 | 3.84 | 61 | 20.3 |
| 33 | MainCh | | | | | | | | | 92.2 | | | | |
| 35 | A | 83.5 | 2.4 | 91 | 12.5 | 82.6 | 2.1 | 79.9 | 0.4 | 79.6 | 1 | | 4 | 1.3 |
| 35 | S | 81.1 | 7.6 | 75.3 | 3.3 | 86.9 | 3.8 | 79.5 | 5.8 | 82.6 | 3 | | 15 | 5 |

| | | | | | | | | | | | | | | |
|---|---|---|---|---|---|---|---|---|---|---|---|---|---|---|
| 35 | K! | 65.1 | 0.5 | 76.7 | 1.4 | 83.1 | 0.5 | 73.7 | 1.2 | n/a | n/a | 3.42 | 28 | 9.3 |
| 35 | MainCh | 92.2 | | 94.5 | | 90.8 | | 94.7 | | 90.4 | | | | |
| 37 | K! | | | | | | | 81.6 | 0.6 | | | 3.42 | 77 | 25.7 |
| 37 | MainCh | | | | | | | 86.6 | | | | | | |
| **38** | D! | 95.6 | 2.2 | 96.4 | 1.4 | 95 | 1.7 | | | 97.3 | 1.4 | 1.38 | 299 | 99.7 |
| **38** | N | 93.4 | 11.4 | 95 | 9.4 | 93.3 | 7.4 | | | 95.9 | 9.1 | | 0 | 0 |
| **38** | L | 82 | 1.3 | 85.6 | 2.1 | 81.5 | 0.4 | | | 86.8 | 6.6 | | 0 | 0 |
| **38** | MainCh | 95.2 | | 94.4 | | 97.3 | | | | 97.9 | | | | |
| **40** | T! | 94.7 | 2.8 | 95.9 | 2.8 | | | 93.7 | 2.7 | 94.5 | 0.7 | 1.38 | 140 | 46.7 |
| **40** | V | 91.9 | 6.5 | 93.1 | 7.8 | | | 91 | 5.1 | 93.8 | 9.5 | | 26 | 8.7 |
| **40** | MainCh | 94.2 | | 95.6 | | | | 93.8 | | 93.7 | | | | |
| **41** | D! | 94.8 | 1.1 | 96.3 | 0.5 | 94.8 | 1.1 | 93.9 | 2.9 | | | 1.38 | 298 | 99.3 |
| **41** | N | 93.7 | 3.7 | 95.8 | 4 | 93.7 | 1.5 | 91 | 3 | | | | 0 | 0 |
| **41** | L | 90 | 0.1 | 91.8 | 0.2 | 92.2 | 2.2 | 88 | 0.6 | | | | 0 | 0 |
| **41** | MainCh | 93.9 | | 96.3 | | 94.3 | | 92.2 | | | | | | |
| 42 | R! | 86.2 | 2.7 | | | | | 88 | 1.5 | 84.4 | 0.9 | 1.38 | 169 | 56.3 |
| 42 | MainCh | 94 | | | | | | 91.8 | | 95.2 | | | | |
| 45 | Y! | | | | | | | | | 91.1 | 2.6 | 1.65 | 12 | 4 |
| 45 | MainCh | | | | | | | | | 93 | | | | |
| 47 | H! | | | | | | | | | 94 | 2.9 | 1.65 | 298 | 99.3 |
| 47 | MainCh | | | | | | | | | 91.9 | | | | |
| 53 | T | 74.6 | 0.6 | 71.1 | 2.2 | 83.3 | 2.7 | 67.2 | 0.3 | 79.6 | 1.2 | | 5 | 1.7 |
| 53 | K! | 74 | 0.8 | 79.8 | 3 | 76.1 | 1.5 | 76.7 | 1.1 | 76.5 | 1.1 | 2.25 | 79 | 26.3 |
| 53 | V | 73.2 | 0.2 | 68.9 | 0.4 | 80.6 | 1.5 | 64.7 | 1.2 | 82 | 2.4 | | 4 | 1.3 |
| 53 | A | 73 | 1.5 | 82.4 | 2.6 | 74.6 | 1.3 | 69.4 | 0.1 | 73.5 | 6 | 0.86 | 19 | 6.3 |
| 53 | MainCh | 86.7 | | 91.6 | | 86.7 | | 85.1 | | 84.7 | | | | |
| 54 | L! | | | 88.6 | 0.9 | | | 87.6 | 2.2 | | | 2.25 | 118 | 39.3 |

| | | | | | | | | | | | | | | |
|---|---|---|---|---|---|---|---|---|---|---|---|---|---|---|
| 54 | N | | | 89 | 0.4 | | | 83.5 | 0.3 | | | | 0 | 0 |
| 54 | D | | | 87.7 | 5.9 | | | 83.2 | 1.2 | | | | 0 | 0 |
| 54 | MainCh | | | 93.6 | | | | 89.5 | | | | | | |
| 55 | V | 79.9 | 1.3 | 87.1 | 1.1 | | | 81.3 | 0.1 | 66.4 | 0.7 | | 7 | 2.4 |
| 55 | T! | 78.6 | 0.4 | 88.6 | 1.5 | | | 78.7 | 12.3 | 58.5 | 0.7 | 2.25 | 30 | 10.2 |
| 55 | S | 78.2 | 2.2 | 86 | 4.6 | | | 81.7 | 0.4 | 64.7 | 0.1 | | 14 | 4.7 |
| 55 | I | 73.5 | 5.3 | 79.1 | 0.3 | | | 64.3 | 0.1 | 72.6 | 5 | | 10 | 3.4 |
| 55 | MainCh | 86.4 | | 90.5 | | | | 70.7 | | 86.7 | | | | |
| 56 | D! | 69.8 | 0.4 | 83.6 | 0.3 | 76.2 | 0.7 | 54* | 3.5 | 73.2 | 3.6 | 2.25 | 27 | 9.2 |
| 56 | N | 69.4 | 0.5 | 86.4 | 2.8 | 87.2 | 8 | 54.5 | 0.5 | 80.7 | 7.5 | | 11 | 3.7 |
| 56 | L | 68.9 | 6.7 | 83.3 | 6.8 | 79.2 | 0.7 | 47.3 | 0.1 | 69 | 2.4 | | 0 | 0 |
| 56 | MainCh | 85.5 | | 92.6 | | 91.8 | | 72.8 | | 77.1 | | | | |
| 58 | N! | | | 92 | 0.2 | 92 | 0.2 | | | | | 2.25 | 70 | 23.6 |
| 58 | L | | | 90 | 8.3 | 90 | 8.3 | | | | | | 22 | 7.4 |
| 58 | D | | | 91.8 | 1.8 | 91.8 | 1.8 | | | | | 1.29 | 31 | 10.4 |
| 58 | MainCh | | | 93.5 | | 92.1 | | | | | | | | |
| 59 | P! | | | 88.1 | 2.5 | | | | | | | 2.25 | 191 | 64.1 |
| 59 | MainCh | | | 89.1 | | | | | | | | | | |
| 60 | K! | 68.7 | 0.2 | 87 | 2.7 | 76.8 | 4 | n/a* | n/a | | | 2.25 | 127 | 42.5 |
| 60 | A | 68.5 | 0.9 | 81.5 | 2.6 | 80.9 | 4.1 | 61.9 | 9.9 | | | | 1 | 0.3 |
| 60 | G | 67.6 | 3.9 | 69.2 | 1.8 | 71.6 | 0.3 | 67.5 | 5.6 | | | | 0 | 0 |
| 60 | MainCh | 86.7 | | 87.8 | | 93.1 | | 85.1 | | 76.5 | | | | |
| 63 | R! | | | 81.1 | 0.1 | 87.9 | 1.1 | | | | | 2.42 | 36 | 12.1 |
| 63 | K | | | 78.6 | 2.7 | 88.7 | 0.4 | | | | | | 17 | 5.7 |
| 63 | MainCh | | | 89.9 | | 94.7 | | | | | | | | |
| 65 | S! | | | 88.7 | 2.4 | | | | | | | 2.42 | 89 | 29.9 |
| 65 | MainCh | | | 95.9 | | | | | | | | | | |
| 67 | S! | | | 87 | 1.6 | | | | | 82* | 5.1 | 2.42 | 116 | 39.1 |

| | | | | | | | | | | | | | | |
|---|---|---|---|---|---|---|---|---|---|---|---|---|---|---|
| 67 | A | | | 87 | 0 | | | | | 85.8 | 3.8 | | 5 | 1.7 |
| 67 | MainCh | | | 90.3 | | | | | | 78.7 | | | | |
| 69 | A | 75.4 | 3.1 | 84.5 | 0.5 | 81.4 | 3.8 | 80.9 | 1.9 | 56.8 | 1.2 | | 9 | 3.1 |
| 69 | S | 72.3 | 3.6 | 84 | 4.9 | 82.7 | 0 | 74.3 | 0.6 | 58.9 | 0.4 | | 79 | 27 |
| 69 | L | 68.7 | 2.5 | 78.3 | 2.7 | 82.7 | 1.3 | 79 | 3.5 | 50.8 | 0 | 0.98 | 7 | 2.4 |
| 69 | K! | 62.5 | 2.1 | 75.6* | 3.3 | 70.1* | 3.1 | 68.5* | 2.6 | 53.7* | 0.4 | 2.42 | 22 | 7.5 |
| 69 | MainCh | 92.2 | | 93.8 | | 94.3 | | 92.6 | | 87.4 | | | | |
| 70 | N! | 84.4 | 5.8 | 86.7 | 0.6 | 92.5 | 0.8 | 77.4 | 0.2 | 74 | 0.4 | | 96 | 32.5 |
| 70 | L | 78.6 | 0.3 | 85.1 | 4.1 | 89.7 | 0.8 | 74.2 | 3.2 | 62.2 | 0.1 | | 0 | 0 |
| 70 | D | 78.3 | 1.8 | 86.1 | 1 | 91.7 | 1.9 | 71 | 1.7 | 60.9 | 0.1 | 2.42 | 44 | 14.9 |
| 70 | MainCh | 92.8 | | 95.9 | | 96.2 | | 88 | | 85.5 | | | | |
| **72** | T! | | | 94.2 | 2.7 | 93.6 | 2.5 | | | | | 2.51 | 62 | 21 |
| **72** | V | | | 91.5 | 2.8 | 91.1 | 5.4 | | | | | | 15 | 5.1 |
| **72** | MainCh | | | 94.9 | | 95.5 | | | | | | | | |
| 73 | I! | | | 92.3 | 1.8 | | | | | | | 2.51 | 146 | 49 |
| 73 | MainCh | | | 95.8 | | | | | | | | | | |
| 74 | V! | | | | | 93.8 | 2.2 | | | 78.5 | 8.2 | 2.51 | 55 | 18.5 |
| 74 | T | | | | | 91.6 | 4.8 | | | 79.5 | 1 | 1.27 | 107 | 35.9 |
| 74 | MainCh | | | | | 94.5 | | | | 78.4 | | | | |
| 76 | G! | | | | | | | 78.8 | 2.8 | 53.8 | 1.6 | 2.51 | 80 | 26.8 |
| 76 | MainCh | | | | | | | 87.8 | | 86.2 | | | | |
| 80 | S | 61.1 | 3.8 | 67.5 | 2.3 | 68.9 | 0.6 | 55.6 | 2.6 | 75.5 | 7.2 | | 27 | 9.1 |
| 80 | A! | 57.3 | 0.7 | 68.1 | 0.6 | 67.6 | 0.4 | n/a | n/a | 65.1 | 1 | 2.51 | 25 | 8.4 |
| 80 | MainCh | 81.8 | | 88 | | 74.2 | | 80 | | 86 | | | | |
| 82 | L! | | | | | 95.2 | 2.3 | 91.9 | 1.9 | 78.2 | 0.9 | 2.51 | 12 | 4.1 |
| 82 | S | | | | | 89.2 | 2.6 | 90 | 3.3 | 79.1 | 0.9 | | 4 | 1.4 |
| 82 | N | | | | | 92.9 | 0.6 | 86.7 | 0.7 | 77.3 | 0.1 | 1.77 | 0 | 0 |
| 82 | D | | | | | 92.3 | 3.1 | 86 | 1.6 | 76.1 | 1.4 | 2.14 | 16 | 5.4 |

| | | | | | | | | | | | | | | |
|---|---|---|---|---|---|---|---|---|---|---|---|---|---|---|
| 82 | MainCh | | | | | 94 | | 88.1 | | 76.5 | | | | |
| 83 | K! | | | | | 68.1* | 0 | | | | | 2.51 | 81 | 27.3 |
| 83 | A | 77.4 | 2.6 | 78.9 | 0.5 | 80.6 | 0.3 | 75.9 | 3.2 | 75.2 | 0.8 | | 42 | 14.1 |
| 83 | MainCh | 90.8 | | 94.8 | | 91.4 | | 89.7 | | 81.3 | | | | |
| 84 | E! | 67.7 | 1.9 | 87.4 | 1.5 | 75.8 | 0.4 | 53.2* | 4.1 | 77 | 0.3 | 2.52 | 24 | 8.1 |
| 84 | Q | 65.7 | 3.2 | 89.5 | 2.1 | 75.4 | 3.2 | 56.4 | 0.9 | 83.3 | 5.9 | 1.49 | 75 | 25.3 |
| 84 | MainCh | 90.4 | | 94.8 | | 89.9 | | 88.4 | | 85.8 | | | | |
| 85 | L! | | | | | 95.4 | 2.5 | | | | | 2.52 | 87 | 29.3 |
| 85 | D | | | | | 92.9 | 0.4 | | | | | | 0 | 0 |
| 85 | N | | | | | 92.5 | 8.2 | | | | | | 2 | 0.7 |
| 85 | MainCh | | | | | 95.8 | | | | | | | | |
| 87 | E! | | | 96.8 | 2.3 | | | | | | | 2.52 | 151 | 50.8 |
| 87 | Q | | | 94.5 | 3.2 | | | | | | | | 18 | 6.1 |
| 87 | MainCh | | | 96.9 | | | | | | | | | | |
| 89 | D! | 93.7 | 2.6 | 94.8 | 0.8 | 95.4 | 2.1 | 94 | 1.8 | 87.5 | 1.9 | 2.52 | 294 | 99 |
| 89 | N | 91.1 | 6.8 | 94 | 2.3 | 93.3 | 4.3 | 92.2 | 9.8 | 85.6 | 3.3 | | 1 | 0.3 |
| 89 | L | 84.3 | 3.1 | 87.6 | 3.8 | 83.9 | 1.7 | 82.4 | 1.6 | 82.3 | 2.5 | | 0 | 0 |
| 89 | MainCh | 93.4 | | 92.8 | | 94.8 | | 94.5 | | 91.1 | | | | |
| 90 | K! | | | | | | | | | 92.3 | 1.9 | 2.52 | 107 | 36 |
| 90 | MainCh | | | | | | | | | 92.1 | | | | |
| 92 | V! | 93.4 | 2.1 | 95.3 | 2 | 93.5 | 5.3 | | | | | 2.52 | 32 | 10.8 |
| 92 | T | 91.3 | 11.1 | 93.3 | 7.7 | 94.4 | 0.9 | | | | | | 1 | 0.3 |
| 92 | MainCh | 93.7 | | 93.9 | | 93.9 | | | | | | | | |
| 94 | I! | | | | | 92.9 | 1.2 | | | | | 2.52 | 53 | 18 |
| 94 | MainCh | | | | | 93.3 | | | | | | | | |
| 97 | R! | | | | | | | | | 90 | 0.3 | 2.52 | 64 | 22.1 |
| 97 | MainCh | | | | | | | | | 94.3 | | | | |

| | | | | | | | | | | | | | | |
|---|---|---|---|---|---|---|---|---|---|---|---|---|---|---|
| 98 | E! | 91.4 | 2.8 | | | | | 87.2 | 0.4 | 92.8 | 2.1 | 2.37 | 27 | 9.9 |
| 98 | Q | 88.6 | 2.9 | | | | | 86.3 | 3.5 | 93 | 0.2 | 2.01 | 23 | 8.4 |
| 98 | MainCh | 93.1 | | | | | | 90 | | 95.3 | | | | |
| 99 | N! | 93.3 | 2.7 | 94.5 | 2.4 | | | | | 93 | 0.1 | 2.37 | 97 | 38.8 |
| 99 | D | 90.6 | 0.2 | 91 | 4.1 | | | | | 92.9 | 1.9 | 2.01 | 0 | 0 |
| 99 | L | 90.4 | 5.6 | 92.1 | 1.1 | | | | | 94.1 | 1.1 | | 0 | 0 |
| 99 | MainCh | 95.2 | | 96.7 | | | | | | 95.1 | | | | |
| 100 | L! | 90.8 | 2.1 | | | 88.2 | 0.9 | | | 93.6 | 0.3 | 2.37 | 79 | 56 |
| 100 | N | 88.7 | 0.6 | | | 87.1 | 4.5 | | | 93.3 | 0.3 | 1.75 | 4 | 2.8 |
| 100 | D | 88.1 | 4 | | | 87.3 | 0.2 | | | 93 | 7.7 | | 2 | 1.4 |
| 100 | MainCh | 92.1 | | | | 93.2 | | | | 91.8 | | | | |
| 101 | S | 74.8 | 1.5 | 75.7 | 0.9 | 67.4 | 2.2 | 79.8 | 0.3 | 83.8 | 3.4 | | 8 | 5.7 |
| 101 | N | 73.3 | 0.6 | 84.1 | 0.5 | 79 | 1.6 | 67.9 | 1 | 79 | 0.8 | 1.95 | 14 | 9.9 |
| 101 | D! | 72.7 | 0.9 | 83.6 | 6.2 | 76.1 | 1.3 | 66.9 | 1.1 | 77.9 | 0.8 | 2.37 | 27 | 19.1 |
| 101 | MainCh | 79.6 | | 84.4 | | 74 | | 75.7 | | 89.4 | | | | |
| 102 | T! | | | | | | | 94.2 | 2.6 | | | 2.37 | 108 | 45.4 |
| 102 | S | | | | | | | 91.6 | 4.1 | | | | 42 | 17.6 |
| 102 | V | | | | | | | 87.5 | 0.2 | | | | 2 | 0.8 |
| 102 | MainCh | | | | | | | 89.9 | | | | | | |
| 104 | N! | 90.9 | 2.7 | 87.8 | 0.3 | | | | | 90.7 | 2.6 | 2.37 | 135 | 57.2 |
| 104 | D | 88.2 | 1.3 | 87.5 | 3.5 | | | | | 88.1 | 0.8 | 2.32 | 22 | 9.3 |
| 104 | L | 86.9 | 4.1 | 84 | 0.9 | | | | | 87.3 | 3 | | 1 | 0.4 |
| 104 | MainCh | 94.1 | | 90 | | | | | | 96.7 | | | | |
| 105 | S | 75 | 2.9 | | | 77.1 | 1.2 | 69.2 | 1.8 | 68.3 | 2.4 | 1.22 | 13 | 5.6 |
| 105 | A | 72.1 | 2.3 | | | 69 | 0 | 75 | 5.8 | 71.5 | 0.6 | 1.21 | 1 | 0.4 |
| 105 | K! | 69.8 | 0.2 | | | 70.2 | 0.7 | 65.2* | 1.6 | 73.2 | 1.7 | 2.37 | 85 | 36.3 |
| 105 | MainCh | 90.4 | | | | 91.8 | | 80.1 | | 93.9 | | | | |
| 106 | S | 80.1 | 4 | | | 78.4 | 8.9 | 80.9 | 1.1 | 86.5 | 3.3 | | 18 | 7.9 |

| | | | | | | | | | | | | | | |
|---|---|---|---|---|---|---|---|---|---|---|---|---|---|---|
| 106 | K! | 68.1 | 0.4 | | | 64.7* | 0.1 | 70.1* | 0.8 | 72.3* | 2.5 | 2.85 | 91 | 39.7 |
| 106 | MainCh | 90.8 | | | | 91.6 | | 90.7 | | 89.9 | | | | |
| 107 | Y! | | | | | 87.2 | 2.2 | 77.4 | 0.6 | | | 2.85 | 150 | 67 |
| 107 | K | | | | | 87.6 | 0.4 | 87.1 | 1.8 | | | | 1 | 0.4 |
| 107 | R | | | | | 89.1 | 1.5 | 87.9 | 0.8 | | | | 0 | 0 |
| 107 | MainCh | | | | | 90.3 | | 83.5 | | 90.8 | | | | |
| 108 | K! | 72.5 | 2.7 | | | 76.5 | 0.4 | 57.9 | 1.3 | 81.3 | 0.9 | 2.85 | 63 | 30.9 |
| 108 | A | 69.8 | 0.1 | | | 79.9 | 1.7 | 62.1 | 0.6 | 75.4 | 0.1 | | 9 | 4.4 |
| 108 | MainCh | 83.3 | | | | 89.7 | | 72.4 | | 91.3 | | | | |
| 110 | L | 71 | 4.3 | 76 | 0.3 | 68.6 | 1.7 | 70.3 | 4.6 | 65.3 | 1.4 | 2.16 | 22 | 16.2 |
| 110 | D | 66.7 | 2.6 | 80.8 | 4.8 | 66.9 | 1.6 | 65.7 | 4.3 | 51.8 | 1.3 | | 0 | 0 |
| 110 | N! | 59.7 | 0.9 | 86 | 5.2 | 69.1 | 0.5 | n/a | n/a | 50.2 | 0.5 | 3.09 | 3 | 2.2 |
| 110 | G | 58.8 | 0.7 | 58.9 | 0.6 | 70.7 | 1 | 74 | 3.7 | 75.6 | 10.3 | 1.3 | 0 | 0 |
| 110 | MainCh | 69.4 | | 73 | | 62.9 | | 68.7 | | 68.2 | | | | |
| 111 | F | 62.1 | 0.5 | | | n/a | n/a | 62.5 | 0.3 | 71.6 | 0.3 | | 14 | 11.1 |
| 111 | Y! | 61.6 | 0.4 | | | n/a | n/a | 59.7 | 1.6 | 69.9 | 2 | 3.09 | 5 | 4 |
| 111 | H | 61.2 | 0.4 | | | n/a | n/a | 61.1 | 0.1 | 74.2 | 1.6 | | 2 | 1.6 |
| 111 | A | 52.2 | last | | | n/a | n/a | 63.3 | 0.2 | 62.5 | 2.3 | | 1 | 0.8 |
| 111 | MainCh | 59.6 | | | | 60.2 | | 75 | | 42.7 | | | | |
| 112 | G | 67.8 | 0.5 | 74 | 2.2 | n/a | n/a | n/a | n/a | n/a | n/a | 1.3 | 1 | 3 |
| 112 | A | 67.3 | 0.8 | 67.9 | 0.2 | 68 | 2 | 74.3 | 1.1 | 68.3 | 2.2 | | 0 | 0 |
| 112 | S | 65.7 | 1 | 53.9 | 1.4 | 66 | 0.8 | 77 | 2.7 | 78.7 | 2.7 | 2.15 | 7 | 21.2 |
| 112 | L! | 63.5 | 0.9 | 59.4* | 0.5 | 65.2 | 0.5 | 73.2* | 3.3 | 73.9* | 1.5 | 3.09 | 2 | 6.1 |
| 112 | N | 62 | 0.4 | 51.8 | 1.5 | 69.7 | 0.7 | 68.8 | 1.9 | 72.2 | 3.9 | | 1 | 3 |
| 112 | D | 61.6 | 6.8 | 50.3 | 2.2 | 69 | 1 | 66.9 | 1 | 76 | 2.1 | | 0 | 0 |
| 112 | MainCh | 67.7 | | 71.2 | | 56.2 | | 82.4 | | 57 | | | | |
| 113 | L | 63.6 | 2.5 | 49.2 | 0.5 | n/a | n/a | 64.5 | 0.6 | n/a | n/a | | 1 | 0.8 |
| 113 | A | 61.1 | 1.1 | n/a | n/a | n/a | n/a | 68.7 | 4.2 | n/a | n/a | 1.3 | 5 | 4.2 |
| 113 | K! | n/a | n/a | 48.7* | 0.9 | n/a* | n/a | n/a* | n/a | 65.4 | last | 3.09 | 38 | 31.9 |
| 113 | MainCh | 82.4 | | 85.3 | | 77.7 | | 92.1 | | 85.1 | | | | |

| | | | | | | | | | | | | | | |
|---|---|---|---|---|---|---|---|---|---|---|---|---|---|---|
| 114 | P! | 70 | 0.6 | 68.7 | 1.1 | 81.5 | 1.1 | | | | | 3.09 | 17 | 14.4 |
| 114 | V | 69.4 | 3.1 | 67.6 | 2.1 | 83.1 | 0.5 | | | | | | 8 | 6.8 |
| 114 | MainCh | 87.3 | | 91 | | 92.7 | | | | | | | | |
| 115 | F! | 77 | 1.9 | | | 83 | 0.7 | 78.2 | 4.5 | | | 3.09 | 10 | 8.8 |
| 115 | MainCh | 91.3 | | | | 93.4 | | 92.3 | | | | | | |
| 117 | D! | 82.8 | 2.9 | 82.3 | 1.9 | 85.2 | 1.4 | | | 87.1 | 1.3 | | 3 | 3.9 |
| 117 | MainCh | 92.7 | | 93.4 | | 91.6 | | | | 95.2 | | | | |
| 120 | L! | 78.7 | 1.8 | 83.2 | 1.3 | 82.5 | 1.5 | 79.5 | 0.6 | 81 | 0.8 | 1.3 | 3 | 8.6 |
| 120 | S | 76.9 | 0.4 | 81.9 | 1 | 79.1 | 4 | 76.3 | 2.9 | 80.2 | 1.3 | | 2 | 5.7 |
| 120 | N | 76.5 | 1 | 80.9 | 0.9 | 81 | 0.9 | 79.7 | 0.2 | 77.7 | 1 | | 2 | 5.7 |
| 120 | D | 75.5 | 5.2 | 80 | 1.7 | 80.1 | 1 | 78.9 | 2.4 | 76.7 | 1 | | 3 | 8.6 |
| 120 | MainCh | 81.5 | | 79.3 | | 80.6 | | 82.4 | | 90.1 | | | | |
| 121 | P! | 72 | 1 | 81 | 2.7 | 78.4 | 0.5 | 67.8* | 5.1 | 71.2 | 2.7 | 1.3 | 23 | 79.3 |
| 121 | G | 71 | 5.6 | 67.8 | 1.1 | 77.1 | 0.1 | 70.6 | 2.8 | 80.7 | 9.5 | | 0 | 0 |
| 121 | MainCh | 82.6 | | 88.9 | | 83.1 | | 86.4 | | 87.5 | | | | |

Res# stands for residue number; ResType for residues type; Cryst for crystallography, mon for monomer, RSCC for real-space correlation coefficient; ΔCon for contrast difference within other RSCC; MS for mass spectrometry; ! for resolved residue. Residues in bold are the side chain with hydrogen bond with neighbour atom(s). Residues with * had their side chain atoms omitted due to absence of electron density. Residues in bold are the side chain with hydrogen bond with neighbour atom(s).

Supplementary Table 11: Residues of MjTX-I having ambiguity assignment from both crystallographic and mass spectrometry data but being resolved by phylogenetic analysis

| | | Cryst mon ABCD | | Cryst mon A | | Cryst mon B | | Cryst mon C | | Cryst mon D | | MS | ConSurf | |
|---|---|---|---|---|---|---|---|---|---|---|---|---|---|---|
| Res# | ResType | RSCC | ΔCon | RSCC | ΔCon | RSCC | ΔCon | RSCC | ΔCon | RSCC | ΔCon | PrimScore | n | % |
| 16 | N! | | | | | 91.4 | 1.2 | | | | | 2.45 | 72 | 25.6 |
| 16 | D | | | | | 88.1 | 0.2 | | | | | 2.6 | 16 | 5.7 |
| 16 | MainCh | | | | | 96 | | | | | | | | |
| 52 | G | 77.8 | 1.3 | 82.4 | 0.3 | 77.2 | 0.7 | 72.9 | 2.9 | 74.3 | 1.3 | | 66 | 22.1 |
| 52 | S | 76.5 | 0.8 | 84.5 | 2.1 | 76.4 | 1.2 | 76.9 | 1.2 | 71.5 | 1.1 | | 35 | 11.7 |

| | | | | | | | | | | | | | | |
|---|---|---|---|---|---|---|---|---|---|---|---|---|---|---|
| 52 | K! | 73.7 | 0.8 | 86.5 | 2 | 76.5 | 0.1 | 68.8 | 0.8 | 73* | 0.3 | | 28 | 9.4 |
| 52 | I | 72.8 | 0.9 | 75.1 | 4.3 | 78.2 | 1 | 79.8 | 2.9 | 70.2 | 2.3 | 0.86 | 2 | 0.7 |
| 52 | MainCh | 87.9 | | 94.5 | | 89.7 | | 83.8 | | 89 | | | | |
| 61 | A | 70 | 1.4 | 87.6 | 7.3 | | | n/a | n/a | 74.5 | 5.2 | | 4 | 1.3 |
| 61 | G | 68.6 | 0.2 | 67.9 | 2.8 | | | n/a | n/a | 77.1 | 2.6 | | 0 | 0.0 |
| 61 | S | 68.4 | 2 | 80.3 | 0.7 | | | 67.7 | 0.3 | 59.1 | 0.6 | | 13 | 4.3 |
| 61 | K! | 66.4 | 2.6 | 79,6* | 0.6 | | | n/a* | n/a | 67,3* | 3.7 | | 8 | 2.7 |
| 61 | MainCh | 91.5 | | 89.8 | | | | 83.9 | | 92.1 | | | | |
| 70 | N! | 84.4 | 5.8 | 86.7 | 0.6 | 92.5 | 0.8 | 77.4 | 0.2 | 74 | 0.4 | | 96 | 32.5 |
| 70 | L | 78.6 | 0.3 | 85.1 | 4.1 | 89.7 | 0.8 | 74.2 | 3.2 | 62.2 | 0.1 | | 0 | 0.0 |
| 70 | D | 78.3 | 1.8 | 86.1 | 1 | 91.7 | 1.9 | 71 | 1.7 | 60.9 | 0.1 | 2.42 | 44 | 14.9 |
| 70 | MainCh | 92.8 | | 95.9 | | 96.2 | | 88 | | 85.5 | | | | |
| 77 | Q | 61.2 | 2.3 | 79.6 | 1.1 | 65.7 | 0.1 | 59 | 1.2 | 63.5 | 2.7 | 2.14 | 3 | 1.0 |
| 77 | E! | 58.9 | 1.1 | 80.4 | 0.8 | 64.8 | 0.9 | 60.1 | 1.1 | 60,8* | 0.5 | 2.51 | 24 | 8.0 |
| 77 | MainCh | 81.7 | | 93.3 | | 73 | | 56 | | 84.7 | | | | |
| 78 | D | 75.2 | 1.1 | 91.4 | 0 | 74 | 1.2 | 65.5 | 1 | 83.1 | 0.2 | 1.27 | 20 | 6.7 |
| 78 | N! | 74.1 | 0 | 91.4 | 2.3 | 72.8 | 0.5 | 64.3 | 0 | 82.9 | 1.1 | 1.74 | 90 | 30.2 |
| 78 | L | 74.1 | 2.8 | 89.1 | 5.8 | 75.1 | 0.4 | 63.2 | 1.6 | 81.8 | 2.1 | 1.29 | 4 | 1.3 |
| 78 | E | 71.3 | 0.2 | 81.1 | 1.2 | 78.2 | 1.7 | 67.9 | 1.8 | 76.2 | 1.2 | 2.51 | 26 | 8.7 |
| 78 | Q | 66.6 | 0.3 | 67 | 0.4 | 76.5 | 1.4 | 69.7 | 1.1 | 75 | 1.3 | 2.14 | 29 | 9.7 |
| 78 | MainCh | 79 | | 85.3 | | 73.5 | | 72 | | 79.8 | | | | |
| 79 | N! | | | 84.3 | 0.7 | | | 83.2 | 2.3 | 82.4 | 2.4 | 1.77 | 65 | 21.8 |
| 79 | D | | | 83.6 | 2.8 | | | 80.9 | 2.1 | 78.7 | 0.4 | 2.51 | 81 | 27.2 |
| 79 | L | | | 80.8 | 2.9 | | | 78.8 | 1.2 | 80 | 1.3 | | 1 | 0.3 |
| 79 | MainCh | | | 78.6 | | | | 79.6 | | 79.8 | | | | |
| 109 | D | 67.0 | 4.7 | 80.5 | 1.2 | 71.0 | 1.3 | n/a | n/a | 77.2 | 0.2 | 2.85 | 11 | 7.1 |
| 109 | L | 58.9 | 3.1 | 82.6 | 0.9 | 68.6 | 3.7 | n/a | n/a | 77.0 | 0.7 | | 3 | 1.9 |
| 109 | N! | 50.2 | 0.4 | 81.7 | 1.2 | 73.5 | 1.2 | n/a* | n/a | 76.3 | 4.2 | 3.09 | 45 | 29.2 |
| 109 | MainCh | 71.1 | | 66.4 | | 74.2 | | 62.7 | | 84.4 | | | | |

Res# stands for residue number; ResType for residues type; Cryst for crystallography, mon for monomer, RSCC for real-space correlation coefficient; ΔCon for contrast difference within other RSCC; MS for mass spectrometry; PrimScore for Primary Score ! for resolved residue. Residues with * had their side chain atoms omitted due to absence of electron density.

Supplementary Table 12: Residues of crotoxin resolved by crystallography and MS and/or phylogenetic analysis

| Crystallography monomer A | | | | | Crystallography monomer B | | | | | MS | ConSurf | |
|---|---|---|---|---|---|---|---|---|---|---|---|---|
| Res# | ResType | RSCC | ΔCont | RSCC* | Res# | ResType | RSCC | ΔCont | RSCC* | PrimaryScore | n | % |
| | | | | | 1 | H | 97.1 | 4.5 | 97 | 3.52 | 0 | 0.0 |
| 2 | L | 93.6 | 4 | 96.9 | 2 | L | 91.4 | 5.9 | 95.2 | 3.52 | 123 | 89.8 |
| 5 | F | 96.1 | 4.2 | 96.5 | 5 | F | 95.3 | 6 | 96.7 | 3.52 | 127 | 65.5 |
| 8 | M | 97.4 | 5.8 | 95 | 8 | M | 97.3 | 4.7 | 96.4 | 3.52 | 167 | 82.7 |
| 9 | I | 95.5 | 6.3 | 95.2 | 9 | I | 96.1 | 3.5 | 96 | 3.52 | 175 | 85.4 |
| 10 | K | 91 | 9 | 96.8 | 10 | K | 92 | 6.8 | 95.3 | 3.52 | 83 | 40.3 |
| | | | | | 11 | F | 96.1 | 6.1 | 95.1 | 3.52 | 4 | 1.9 |
| 13 | T | 94.8 | 5.8 | 94.1 | 13 | T | 96.8 | 3.6 | 96 | 3.52 | 144 | 68.6 |
| 14 | R | 88.5 | 6.4 | 91 | 14 | R | 84.9 | 3.7 | 94.9 | 3.52 | 12 | 5.6 |
| 15 | K | 84.9 | 8.6 | 93.6 | 15 | K | 90.2 | 4 | 96.3 | 6.32 | 89 | 40.1 |
| 16 | N | 88.1 | 3.9 | 94.4 | | | | | | 6.32 | 59 | 26.3 |
| 17 | A | 92.5 | 11.9 | 92 | 17 | A | 92.1 | 21 | 93.3 | 6.32 | 98 | 43.0 |
| 19 | P | 96.3 | 16.1 | 97 | 19 | P | 96.1 | 16 | 97 | 6.32 | 10 | 4.2 |
| 20 | F | 96 | 4.5 | 96.9 | 20 | F | 95 | 4.3 | 96.8 | 6.32 | 6 | 2.3 |
| 21 | Y | 95.8 | 6 | 96.4 | 21 | Y | 96.4 | 6.6 | 95.2 | 6.32 | 236 | 81.1 |
| 22 | A | 83.8 | 11.3 | 92.1 | 22 | A | 88.4 | 14 | 92.9 | 6.32 | 43 | 14.7 |
| 23 | F | 94 | 5.3 | 95.5 | 23 | F | 95.9 | 2.7 | 84.3 | 6.32 | 37 | 12.4 |
| 24 | Y | 97.1 | 7.8 | 96 | 24 | Y | 95.2 | 6 | 95.4 | 6.32 | 296 | 99.3 |
| 25 | G | 93.2 | 17.4 | 95.9 | 25 | G | 90.5 | 19 | 95.3 | 6.32 | 297 | 99.7 |
| 26 | C | 95.7 | 16.8 | 94.2 | 26 | C | 94.5 | 13 | 94.4 | 6.32 | 294 | 98.7 |
| 27 | Y | 94.1 | 5.3 | 95 | 27 | Y | 95.6 | 7.7 | 95.9 | 6.32 | 225 | 75.5 |
| 28 | C | 97.2 | 18.1 | 94.9 | 28 | C | 95.6 | 10 | 86.2 | 6.32 | 297 | 99.7 |
| 29 | G | 61.8 | 4.6 | 89.9 | 29 | G | 37.9 | 4.7 | 79.1 | 6.32 | 289 | 97.0 |
| 30 | W | 95 | 8.4 | 96.4 | 30 | W | 81 | 3.9 | 83.4 | 6.32 | 43 | 14.4 |
| 31 | G | 90.5 | 31.7 | 95.1 | 31 | G | 64.7 | 30 | 83.1 | 6.32 | 261 | 87.6 |
| 32 | G | 87.6 | 13.3 | 95.3 | 32 | G | 81.3 | 17 | 90.4 | 6.32 | 281 | 94.3 |
| 33 | Q | 91.4 | 6.5 | 94.5 | 33 | A | 78 | 4.8 | 92.5 | 6.32 / - | 41 / 8 | 13.8 / 2.7 |

| | | | | | | | | | | | | |
|---|---|---|---|---|---|---|---|---|---|---|---|---|
| 34 | G | 88.3 | 16.7 | 94.7 | 34 | G | 91 | 17 | 96.6 | 6.32 | 275 | 92.3 |
| 36 | P | 89 | 14.8 | 89.6 | 36 | P | 94.5 | 10 | 95 | 4.95 | 290 | 97.3 |
| 37 | A | 89.6 | 13.4 | 91.7 | 37 | K | 90.9 | 4.5 | 93.2 | - / 4.95 | 3 / 72 | 1.0 / 24,2 |
| 39 | A | 91.8 | 21.1 | 95.7 | 39 | A | 94.4 | 25 | 95.6 | 3.27 | 97 | 32.6 |
| | | | | | 41 | D | 96.9 | 3.2 | 95.2 | 3.27 | 298 | 99.7 |
| | | | | | 42 | R | 87 | 8.6 | 94.2 | 3.55 | 163 | 54.3 |
| 43 | C | 95.6 | 7.6 | 95.7 | 43 | C | 95 | 8.5 | 94 | 3.55 | 297 | 99.7 |
| 44 | C | 95.7 | 16.8 | 96.5 | 44 | C | 95.2 | 8 | 94.4 | 3.55 | 298 | 100.0 |
| 45 | F | 94.3 | 4.2 | 95.4 | 45 | F | 95.2 | 3.4 | 93 | 3.55 | 60 | 20.1 |
| 46 | V | 92.6 | 6.2 | 95 | 46 | V | 92.4 | 4.6 | 96.5 | 3.55 | 80 | 26.8 |
| 47 | H | 94 | 5 | 93.8 | 47 | H | 96.7 | 3.9 | 95.5 | 3.55 | 294 | 98.3 |
| 48 | D | 87.6 | 3.8 | 95.5 | | | | | | 3.55 | 261 | 87.3 |
| 49 | C | 95.5 | 14.2 | 94.8 | 49 | C | 95.7 | 11 | 95 | 3.55 | 169 | 56.5 |
| 50 | C | 94.3 | 10.7 | 94.1 | 50 | C | 95.1 | 7.5 | 96.2 | 3.55 | 298 | 99.7 |
| 51 | Y | 93.5 | 6 | 93.5 | 51 | Y | 96.7 | 6.7 | 95.5 | 3.55 | 281 | 94.0 |
| 52 | G | 82.3 | 34.1 | 93.1 | 52 | G | 82.7 | 23 | 92 | 3.55 | 79 | 26.4 |
| 53 | K | 90.6 | 5.7 | 91.5 | 53 | K | 91.1 | 6.6 | 92.7 | 3.55 | 81 | 27.1 |
| 54 | L | 92.2 | 4.7 | 92.6 | 54 | L | 92.6 | 5.6 | 95.1 | 1.39 | 109 | 36.5 |
| 56 | K | 91 | 6.7 | 90.4 | 56 | K | 87.6 | 9.3 | 93 | 1.39 | 14 | 4.8 |
| 57 | C | 92.7 | 13.6 | 90 | 57 | C | 96.8 | 16 | 97.2 | 3.94 | 294 | 98.7 |
| | | | | | 60 | N | 89.9 | 4.9 | 95 | | 2 | 0.7 |
| 61 | W | 93.1 | 7.7 | 87.1 | 61 | W | 95.7 | 9.7 | 95 | 3.94 | 15 | 5.0 |
| 62 | N | 72.4 | 3.8 | 84.4 | 62 | N | 95.3 | 3.9 | 96.7 | | 26 | 8.7 |
| 64 | Y | 94.9 | 4.4 | 95.4 | 64 | Y | 95.2 | 5.9 | 94.1 | 3.94 | 277 | 92.3 |
| 65 | R | 75.9 | 4.4 | 86.1 | 65 | R | 90 | 4.7 | 95.1 | 2.77 | 23 | 7.7 |
| 66 | Y | 93.1 | 6.7 | 83.5 | 66 | Y | 96.5 | 3.6 | 94.5 | 3.94 | 142 | 47.5 |
| 67 | S | 86.2 | 6.4 | 92.2 | 67 | S | 89.8 | 3.6 | 95 | 3.94 | 104 | 34.8 |
| 68 | L | 93.2 | 5.9 | 93.5 | 68 | L | 93.4 | 3.9 | 95 | 3.94 | 8 | 2.7 |
| | | | | | 69 | A | 77.5 | 7 | 88.5 | | 19 | 6.3 |
| | | | | | 70 | S | 95.8 | 13 | 94.8 | 3.00 | 25 | 8.3 |
| 71 | G | 87 | 35.1 | 91.7 | 71 | G | 86.4 | 23 | 94.1 | 3.00 | 113 | 37.7 |
| 72 | Y | 93.6 | 7.6 | 90.6 | 72 | Y | 93.5 | 3.7 | 93.4 | 3.00 | 3 | 1.0 |
| | | | | | 73 | I | 95.5 | 5.8 | 95.7 | 3.00 | 143 | 47.7 |
| 74 | T | 84.3 | 3.9 | 88.5 | | | | | | 3.00 | 113 | 37.7 |
| 75 | C | 92 | 10.3 | 89.3 | 75 | C | 95.8 | 18 | 95.3 | 3.00 | 298 | 99.7 |
| 76 | G | 69 | None | 87.8 | 76 | G | 88.5 | 32 | 94.5 | 3.00 | 53 | 17.9 |

| | | | | | | | | | | | | |
|---|---|---|---|---|---|---|---|---|---|---|---|---|
| 77 | G | 73.5 | 13.4 | 87.8 | 77 | K | 82.5 | 7 | 91.9 | - / 3.00 | 40 / 42 | 13.7 / 14.3 |
| 78 | G | 65.8 | 8.7 | 75.6 | 78 | G | 83 | 14 | 94.3 | 3.97 | 52 | 17.9 |
| | | | | | 79 | T | 92.5 | 4.3 | 93.8 | 3.97 | 52 | 17.9 |
| 80 | W | 92.4 | 9.7 | 93.2 | 80 | W | 96.5 | 12 | 95.1 | 3.97 | 26 | 8.9 |
| 81 | C | 91.4 | 8.1 | 91.2 | 81 | C | 93.6 | 8.4 | 94.5 | 3.97 | 280 | 96.2 |
| 82 | E | 85.8 | 7.4 | 94.2 | | | | | | 3.97 | 90 | 31.0 |
| 83 | E | 88.3 | 3.1 | 95.2 | | | | | | 3.97 | 11 | 3.8 |
| 84 | Q | 89 | 5.2 | 96.7 | 84 | Q | 95.4 | 3.9 | 96.6 | 3.97 | 59 | 20.3 |
| 86 | C | 93.5 | 3.2 | 93.8 | 86 | C | 96.6 | 15 | 94.2 | 3.97 | 289 | 100.0 |
| | | | | | 87 | E | 93 | 4.8 | 95.6 | 3.97 | 126 | 43.9 |
| 88 | C | 96.3 | 8.5 | 95.6 | 88 | C | 96.6 | 14 | 96.7 | 3.97 | 287 | 99.3 |
| 90 | R | 95 | 6.5 | 95.7 | 90 | R | 94.3 | 3.6 | 96 | 3.97 | 124 | 43.1 |
| | | | | | 91 | V | 95.2 | 4 | 97 | 3.80 | 24 | 8.4 |
| 92 | A | 95.8 | 17.9 | 97.7 | 92 | A | 93.8 | 11 | 96.9 | 3.80 | 199 | 69.3 |
| 93 | A | 93.3 | 14.1 | 95.8 | 93 | A | 95.1 | 20 | 97 | 3.80 | 236 | 82.2 |
| 94 | E | 93 | 6.4 | 95.8 | 94 | E | 90.2 | 2.6 | 93.6 | 3.80 | 52 | 18.1 |
| | | | | | 95 | C | 94 | 14 | 95.9 | 3.80 | 284 | 99.0 |
| 96 | L | 95.6 | 3 | 94.3 | 96 | L | 96.4 | 3.2 | 97.8 | 3.80 | 83 | 28.9 |
| 97 | R | 92.5 | 3.8 | 95.5 | 97 | R | 92.5 | 3.8 | 95.9 | 3.80 | 53 | 18.7 |
| 98 | A | 86.3 | 5.3 | 91.3 | 98 | R | 95.4 | 5.5 | 94.6 | 3.64 / 2.99 | 24 / 70 | 9.5 / 27,8 |
| 99 | S | 94 | 11 | 95.4 | 99 | S | 96.2 | 11 | 96.4 | 3.74 | 35 | 15.0 |
| 101 | S | 82.8 | 8.7 | 90.6 | 101 | S | 81.5 | 4.8 | | 3.74 | 7 | 6.0 |
| 102 | T | 94.1 | 3.9 | 93.9 | 102 | T | 94.4 | 5 | 95.1 | 3.74 | 89 | 41.2 |
| 103 | Y | 95.3 | 5.7 | 90.1 | 103 | Y | 94.7 | 4.6 | 95.6 | 3.74 | 172 | 83.5 |
| 104 | T | 89.1 | 6 | 92.9 | | | | | | 3.64 | 3 | 1.5 |
| 105 | N | 91 | 4.2 | 90.8 | 105 | Y | 93.2 | 1.9 | 96.7 | 3.74 / 2.99 | 22 / 0 | 11.1 / 0,0 |
| 106 | G | 72.5 | 7 | 90.4 | 106 | G | 92.6 | 25 | 94.2 | 3.74 | 4 | 2.1 |
| 107 | Y | 91.2 | 3.1 | 94 | 107 | Y | 94.9 | 3 | 96.1 | 3.74 | 119 | 61.7 |
| 108 | M | 95.4 | 6 | 95.6 | 108 | M | 95 | 0.8 | 92.6 | 3.74 | 9 | 4.8 |
| 109 | F | 96.4 | 3.2 | 94.3 | 109 | F | 96.2 | 3.5 | 90.6 | 3.74 | 46 | 24.7 |
| 110 | Y | 96.4 | 4.8 | 94.8 | 110 | Y | 95 | 3 | 96.1 | 3.74 | 92 | 51.1 |
| 111 | P | 83 | 12.4 | 90 | 111 | P | 91.5 | 16 | 92.2 | 3.74 | 104 | 58.4 |
| | | | | | 112 | D | 88.1 | 6.4 | 93.7 | 3.74 | 32 | 18.1 |
| 113 | S | 77.9 | 8 | 83.2 | | | | | | 3.74 | 23 | 12.8 |
| 114 | R | 84.4 | 3.6 | 88.6 | 114 | R | 83 | 1.4 | 93 | 3.74 | 33 | 18.5 |
| 115 | C | 96 | 18.2 | 83.3 | 115 | C | 89.1 | 7 | 88.6 | 1.25 | 179 | 100.0 |

| | | | | | | | | | | | | |
|---|---|---|---|---|---|---|---|---|---|---|---|---|
| 116 | G | 57.7 | 5.2 | 65.7 | 116 | G | 53.9 | None | 73.1 | 1.25 | 41 | 27.5 |
| 118 | A | 84.4 | 11.4 | 80.9 | 118 | P | 92 | 18 | 91.2 | - / 1.25 | 4 / 35 | 3.0 / 26,1 |
| 119 | S | 86.8 | 8.6 | 90.5 | | | | | | 1.25 | 23 | 18.7 |
| 122 | C | 85.4 | 12.4 | 77.2 | 122 | C | 61.1 | 7.9 | 76.7 | 1.25 | 108 | 100.0 |

Res# stands for residue number; ResType for residues type; RSCC for real-space correlation coefficient; ΔCont for contrast difference within other RSCC; *for main chain atoms; MS for mass spectrometry.

Supplementary Table 13: Residues of crotoxin having assignment ambiguity from crystallographic data being resolved by mass spectrometry

| Crystallography monomer A | | | | | MS | ConSurf | | Crystallography monomer B | | | | | | ConSurf | |
|---|---|---|---|---|---|---|---|---|---|---|---|---|---|---|---|
| Res# | ResType | RSCC | ΔCo | Observation | PrScore | n | % | Res# | ResType | RSCC | ΔCo | Observation | PrScore | n | % |
| 1 | S! | 73.2 | 0.5 | | 3.09 | | | | | | | | | | |
| 1 | K | 72.7 | 0.1 | | | | | | | | | | | | |
| 1 | T | 72.6 | 1.3 | | | | | | | | | | | | |
| 1 | V | 71.3 | 1.5 | | | | | | | | | | | | |
| 1 | MainCh | 90.1 | | | | | | | | | | | | | |
| 3 | V | 74.2 | 1.3 | | | 20 | 12.2 | 3 | L! | 93 | 0.2 | | 3.52 | 79 | 48.2 |
| 3 | L! | 72.9 | 0.7 | | 3.52 | 79 | 48.2 | 3 | N | 92.9 | 1.4 | | | 0 | 0.0 |
| 3 | A | 72.2 | 2 | | | 4 | 2.4 | 3 | D | 91.4 | 6.2 | | | 0 | 0.0 |
| 3 | MainCh | 90.6 | | | | | | 3 | MainCh | 94.8 | | | | | |
| 4 | E | 93.2 | 2.1 | | | 30 | 16.0 | 4 | Q! | 96.8 | 2 | | 3.52 | 109 | 58.3 |
| 4 | Q! | 91.2 | 5 | | 3.52 | 109 | 58.3 | 4 | E | 94.8 | 8.9 | | | 30 | 16.0 |
| 4 | MainCh | 95 | | | | | | 4 | MainCh | 97.1 | | | | | |
| 6 | D | 88.8 | 3.5 | | | 3 | 1.5 | 6 | N! | 94.5 | 0.5 | 3 H-bonds | 3.52 | 16 | 8.2 |
| 6 | N! | 85.3 | 3.8 | | 3.52 | 16 | 8.2 | 6 | D | 94 | 5.8 | | | 3 | 1.5 |
| 6 | L | 81.5 | 0.6 | | | 0 | 0.0 | 6 | MainCh | 96.8 | | | | | |
| 7 | T | 89.3 | 5.6 | | | 11 | 5.6 | 7 | V | 87.8 | 1.6 | | | 6 | 3.0 |
| 7 | V | 83.7 | 3.9 | | | 6 | 3.0 | 7 | T | 86.2 | 1.5 | | | 11 | 5.6 |
| 7 | S | 79.8 | 1.3 | | | 37 | 18.7 | 7 | S | 84.7 | 1.6 | | | 37 | 18.7 |
| 7 | K! | 62.1 | 0.5 | | 3.52 | 38 | 19.2 | 7 | K! | 83.1 | 0.4 | | 3.52 | 38 | 19.2 |
| 7 | MainCh | 94.3 | | | | | | 7 | MainCh | 95.9 | | | | | |
| 11 | F! | 94.5 | 2.8 | | 3.52 | 4 | 1.9 | | | | | | | | |

| | | | | | | | | | | | | | | |
|---|---|---|---|---|---|---|---|---|---|---|---|---|---|---|
| 11 | H | 91.7 | 3.1 | | 19 | 9.2 | | | | | | | | |
| 11 | MainCh | 94.7 | | | | | | | | | | | | |
| 12 | E! | 93.9 | 1.8 | 3.52 | 12 | 5.8 | 12 | E! | 94.7 | 1 | | 3.52 | 12 | 5.8 |
| 12 | Q | 92.2 | 5.4 | | 0 | 0.0 | 12 | Q | 93.7 | 5.4 | | | 0 | 0.0 |
| 12 | MainCh | 92.6 | | | | | 12 | MainCh | 96.3 | | | | | |
| | | | | | | | 16 | N! | 93.8 | 1.7 | | 6.32 | 59 | 26.3 |
| | | | | | | | 16 | L | 92.1 | 2.1 | | | 3 | 1.3 |
| | | | | | | | 16 | S | 90.0 | 2.8 | | | 68 | 30.4 |
| | | | | | | | 16 | D | 87.2 | 6.5 | | | 10 | 4.5 |
| | | | | | | | 16 | MainCh | 95.8 | | | | | |
| 18 | V! | 90.9 | 0.4 | 6.32 | 23 | 13.6 | 18 | I! | 92.2 | 2.4 | | 5.47 | 27 | 16.0 |
| 18 | T | 90.5 | 3.9 | | 6 | 3.6 | 18 | V | 89.8 | 2.2 | | 6.32 | 23 | 13.6 |
| 18 | MainCh | 95.2 | | | | | 18 | T | 87.6 | 2.3 | | | 6 | 3.6 |
| | | | | | | | 18 | MainCh | 95.9 | | | | | |
| 35 | A | 77.9 | 1.7 | | 5 | 1.7 | 35 | R! | 86.2 | 1.8 | | 6.32 | 21 | 7.0 |
| 35 | R!* | 67.3 | 2.1 | 6.32 | 21 | 7.0 | 35 | K | 84.4 | 4.9 | | | 14 | 4.7 |
| 35 | MainCh | 95.7 | | | | | | | 95.7 | | | | | |
| 38 | D! | 95.2 | 1.2 | 3.27 | 297 | 99.7 | 38 | D! | 97 | 2.2 | | 3.27 | 297 | 99.7 |
| 38 | N | 94 | 8.3 | | 0 | 0.0 | 38 | N | 94.8 | 6.4 | | | 0 | 0.0 |
| 38 | MainCh | 94.7 | | | | | 38 | MainCh | 95 | | | | | |
| 40 | T! | 94.8 | 1.5 | 3.27 | 126 | 42.3 | 40 | T! | 95.7 | 1.8 | 1 H-bond | 3.27 | 126 | 42.3 |
| 40 | V | 93.3 | 8 | | 29 | 9.7 | 40 | V | 93.9 | 4.9 | | | 29 | 9.7 |
| 40 | MainCh | 96.3 | | | | | 40 | MainCh | 96.2 | | | | | |
| 42 | R! | 84 | 0.3 | 3.55 | 163 | 54.3 | | | | | | | | |
| 42 | A | 92 | | | 4 | 1.3 | | | | | | | | |
| 42 | MainCh | 95.1 | | | | | | | | | | | | |
| | | | | | | | 48 | D! | 89.5 | 1 | | 3.55 | 261 | 87.3 |
| | | | | | | | 48 | S | 88.5 | 3.5 | | | 0 | 0.0 |

| | | | | | | | | | | | | | | | |
|---|---|---|---|---|---|---|---|---|---|---|---|---|---|---|---|
| | | | | | | | | 48 | MainCh | 94.3 | | | | | |
| 55 | A! | 70.2 | 0.8 | | 1.39 | 14 | 4.7 | 55 | G | 79.2 | 0.7 | | | 7 | 2.3 |
| 55 | G | 69.5 | 8.6 | | | 7 | 2.3 | 55 | A! | 78.5 | 27 | | 1.39 | 14 | 4.7 |
| 55 | MainCh | 86.6 | | | | | | 55 | MainCh | 94.4 | | | | | |
| 58 | N! | 81.4 | 2.9 | 1 H-bond | 3.94 | 41 | 13.8 | 58 | N! | 96.4 | 2.5 | 2 H-bonds | 3.94 | 41 | 13.8 |
| 58 | D | 78.5 | 4.8 | | | 21 | 7.0 | 58 | D | 93.9 | 3.8 | | | 21 | 7.0 |
| 58 | MainCh | 80.2 | | | | | | 58 | MainCh | 96.3 | | | | | |
| 59 | V | 88 | 0.7 | 1 H-bond | | 8 | 2.7 | 59 | T! | 95.8 | 1.9 | 1 H-bond | 3.94 | 36 | 12.1 |
| 59 | T! | 87.3 | 3.1 | | 3.94 | 36 | 12.1 | 59 | V | 93.9 | 5.6 | | | 8 | 2.7 |
| 59 | MainCh | 86.7 | | | | | | 59 | MainCh | 95.9 | | | | | |
| 63 | V | 89.1 | 1.6 | | | 12 | 4.0 | 63 | T | 91.7 | 0.1 | | | 32 | 10.7 |
| 63 | T | 87.5 | 3.1 | | | 32 | 10.7 | 63 | V | 91.6 | 1.4 | | | 12 | 4.0 |
| 63 | I! | 84.4 | 2.7 | | 3.94 | 20 | 6.7 | 63 | I! | 90.2 | 5.1 | | 3.94 | 20 | 6.7 |
| 63 | MainCh | 90.4 | | | | | | 63 | MainCh | 97.2 | | | | | |
| 70 | S! | 83.5 | 0.8 | | 3.00 | 25 | 8.3 | | | | | | | | |
| 70 | G | 82.7 | 1.2 | | | 44 | 14.7 | | | | | | | | |
| 70 | A | 81.5 | 11.2 | | | 8 | 2.7 | | | | | | | | |
| 70 | MainCh | 87 | | | | | | | | | | | | | |
| 73 | I! | 93.7 | 2.7 | | 3.00 | 143 | 47.7 | | | | | | | | |
| 73 | T | 91.0 | 0.2 | | | 7 | 2.3 | | | | | | | | |
| 73 | V | 90.8 | 7.6 | | | 74 | 24.7 | | | | | | | | |
| 73 | MainCh | 87.6 | | | | | | | | | | | | | |
| 79 | T! | 82.9 | 1.4 | | 3.97 | 52 | 17.9 | | | | | | | | |
| 79 | S | 81.5 | 5.6 | | | 55 | 18.9 | | | | | | | | |
| 79 | MainCh | 86.7 | | | | | | | | | | | | | |
| | | | | | | | | 83 | E! | 94.3 | 1.4 | | 3.97 | 11 | 3.8 |
| | | | | | | | | 83 | Q | 92.9 | 5.9 | | | 21 | 7.2 |
| | | | | | | | | 83 | MainCh | 91.8 | | | | | |

| | | | | | | | | | | | | | | | |
|---|---|---|---|---|---|---|---|---|---|---|---|---|---|---|---|
| 85 | I! | 94.3 | 2.3 | | 3.97 | 77 | 26.6 | 85 | I! | 92.5 | 0.1 | | 3.97 | 77 | 26.6 |
| 85 | T | 91.9 | 0.7 | | | 23 | 7.9 | 85 | V | 92.4 | 2 | | | 54 | 18.6 |
| 85 | V | 91.2 | 5.3 | | | 54 | 18.6 | 85 | T | 90.4 | 1.5 | | | 23 | 7.9 |
| 85 | MainCh | 96.5 | | | | | | 85 | MainCh | 96.4 | | | | | |
| 87 | E! | 93.7 | 2.1 | | 3.97 | 126 | 43.9 | 87 | E! | 93 | 4.8 | | 3.97 | 126 | 43.9 |
| 87 | Q | 91.6 | 7.7 | | | 20 | 7.0 | 87 | MainCh | 95.6 | | | | | |
| 87 | MainCh | 96.3 | | | | | | | | | | | | | |
| 89 | D! | 96.9 | 1.5 | 4 H-bonds | 3.97 | 283 | 97.9 | 89 | D! | 96.9 | 1.7 | 4 H-bonds | 3.97 | 283 | 97.9 |
| 89 | N | 95.4 | 3.6 | | | 2 | 0.7 | 89 | N | 95.2 | 7 | | | 2 | 0.7 |
| 89 | MainCh | 96.2 | | | | | | 89 | MainCh | 97.9 | | | | | |
| 91 | V! | 95.3 | 1.3 | | 3.97 | 24 | 8.4 | | | | | | | | |
| 91 | T | 94 | 2.8 | | | 42 | 14.6 | | | | | | | | |
| 91 | MainCh | 96.5 | | | | | | | | | | | | | |
| 95 | C! | 91.3 | 0.2 | | 3.97 | 284 | 99.0 | | | | | | | | |
| 95 | S | 91.1 | 27.0 | | | 1 | 0.3 | | | | | | | | |
| 95 | MainCh | 93.6 | | | | | | | | | | | | | |
| | | | | | | | | 104 | L | 84.1 | 0.8 | | 3.74 | 0 | 0.0 |
| | | | | | | | | 104 | A | 83.2 | 0.7 | | | 0 | 0.0 |
| | | | | | | | | 104 | S | 82.6 | 5.5 | | | 13 | 6.5 |
| | | | | | | | | 104 | K! | 77 | 1.4 | | 2.99 | 13 | 6.5 |
| | | | | | | | | 104 | N | 75.7 | 1.3 | | | 106 | 52.7 |
| | | | | | | | | 104 | D | 74.4 | 4.1 | | | 12 | 6.0 |
| | | | | | | | | 104 | MainCh | 95.1 | | | | | |
| 112 | K | 68.7 | 1.8 | | | 16 | 9.0 | | | | | | | | |
| 112 | R | 66.8 | 0.8 | | | 26 | 14.7 | | | | | | | | |
| 112 | M | 66.1 | 0.1 | | | 0 | 0.0 | | | | | | | | |
| 112 | A | 65.9 | 0.3 | | | 3 | 1.7 | | | | | | | | |
| 112 | D! | 49.1 | 4.4 | | 3.74 | 32 | 18.1 | | | | | | | | |
| 112 | MainCh | 86.2 | | | | | | | | | | | | | |

| | | | | | | | | | | | | | |
|---|---|---|---|---|---|---|---|---|---|---|---|---|---|
| 117 | G! | | | 1.25 | 72 | 50.0 | 117 | G! | 57.4 | | 1.25 | 72 | 50.0 |
| 117 | MainCh | 60.3 | | | | | 117 | MainCh | 74.7 | | | | |

Res# stands for residue number; ResType for residues type; RSCC for real-space correlation coefficient; ΔCo for contrast difference within other RSCC; MainCh for main chain atoms; MS for mass spectrometry; H-bond for hydrogen bond; ! for resolved residue.

Supplementary Table 14: Residues of crotoxin having ambiguity assignment from both crystallographic and mass spectrometry data but being resolved by phylogenetic analysis

| Crystallography monomer A | | | | MS | ConSurf | | Crystallography monomer B | | | | MS | ConSurf | |
|---|---|---|---|---|---|---|---|---|---|---|---|---|---|
| Res# | ResType | RSCC | ΔCont | PrimaryScore | n | % | Res# | ResType | RSCC | ΔCont | PrimaryScore | n | % |
| 60 | N! | 83 | 2.4 | | 2 | 0.7 | | | | | | | |
| 60 | L | 81.1 | 0.5 | | 6 | 2.0 | | | | | | | |
| 60 | MainCh | 88 | | | | | | | | | | | |
| 69 | A! | 68 | 0.1 | | 19 | 6.3 | | | | | | | |
| 69 | N | 68 | 0.9 | | 9 | 3.0 | | | | | | | |
| 69 | D | 67 | 2.1 | | 7 | 2.3 | | | | | | | |
| 69 | MainCh | 75 | | | | | | | | | | | |
| | | | | | | | 74 | T! | 94.1 | 0.8 | 3.00 | 113 | 37.7 |
| | | | | | | | 74 | V | 93.3 | 7.6 | | 46 | 15.3 |
| | | | | | | | 74 | MainCh | 95.6 | | | | |
| 82 | E! | 82 | 2 | 3.97 | 90 | 31.0 | 82 | E! | 93.3 | 0.6 | 3.97 | 90 | 31.0 |
| 82 | Q | 80 | 1.3 | 3.80 | 63 | 21.7 | 82 | Q | 92.6 | 4.1 | 3.80 | 63 | 21.7 |
| 82 | MainCh | 95 | | | | | | | | | | | |
| 100 | D | 95 | 1 | 3.74 | 0 | 0.0 | 100 | D | 95.6 | 1.2 | 3.74 | 0 | 0.0 |
| 100 | L! | 94 | 0.8 | 2.99 | 55 | 47.4 | 100 | L! | 94.4 | 0.3 | 2.99 | 55 | 47.4 |
| 100 | N | 93 | 5.5 | 3.64 | 0 | 0.0 | 100 | N | 94 | 8.8 | 3.64 | 0 | 0.0 |
| 100 | MainCh | 95 | | | | | 100 | MainCh | 95.7 | | | | |
| 120 | E! | 93 | 2.3 | 1.25 | 19 | 16.4 | 120 | E! | 93.4 | 1.4 | 1.25 | 19 | 16.4 |
| 120 | Q | 91 | 4.6 | | 1 | 0.9 | 120 | Q | 92 | 4.8 | | 1 | 0.9 |

| | | | | | | | | | | | | | |
|---|---|---|---|---|---|---|---|---|---|---|---|---|---|
| 120 | MainCh | 88 | | | | | 120 | MainCh | 95.3 | | | | |
| 121 | T! | 73 | 1.4 | 1.25 | 30 | 26.8 | 121 | T! | 89.6 | 2.9 | 0.38 | 30 | 26.8 |
| 121 | I | 71 | 0.3 | | 0 | 0.0 | 121 | V | 86.7 | 7.1 | | 0 | 0.0 |
| 121 | V | 71 | 2.3 | | 0 | 0.0 | 121 | MainCh | 90.7 | 5.1 | | | |

Res# stands for residue number; ResType for residues type; RSCC for real-space correlation coefficient; ΔCont for contrast difference within other RSCC; MainCh for main chain atoms; MS for mass spectrometry; PrScore for PrimaryScore; H-bond for hydrogen bond; ! for resolved residue.
